# Evolutionary rescue of bacterial populations by heterozygosity on multicopy plasmids

**DOI:** 10.1101/2024.05.29.596466

**Authors:** Ian Dewan, Hildegard Uecker

**Affiliations:** Research Group Stochastic Evolutionary Dynamics, Department of Theoretical Biology, Max Planck Institute for Evolutionary Biology, 24306 Plön, Germany

**Keywords:** evolutionary rescue, heterozygosity, multicopy plasmids, bacterial evolution, plasmid copy number, fitness trade-off

## Abstract

Bacterial plasmids and other extrachromosomal DNA elements frequently carry genes with important fitness effects for their hosts. Multicopy plasmids can additionally carry distinct alleles of host-fitness-relevant genes on different plasmid copies, allowing for heterozygosity not possible for loci on haploid chromosomes. Plasmid-mediated heterozygosity may increase the fitness of bacterial cells in circumstances where there is an advantage to having multiple distinct alleles (heterozyogote advantage); however, plasmid-mediated heterozygosity is also subject to constant loss due to random segregation of plasmid copies on cell division. We analyze a multitype branching process model to study the evolution and maintenance of plasmid-mediated heterozygosity under a heterozygote advantage. We focus on an evolutionary rescue scenario in which a novel mutant allele on a plasmid must be maintained together with the wild-type allele to allow population persistance (although our results apply more generally to the maintenance of heterozygosity due to heterozygote advantage). We determine the probability of rescue and derive an analytical expression for the threshold on the fitness of heterozygotes required to overcome segregation and make rescue possible; this threshold decreases with increasing plasmids copy number. We further show that the formation of cointegrates from the fusion of plasmid copies increases the probability of rescue. Overall, our results provide a rigorous quantitative assessment of the conditions under which bacterial populations can adapt to multiple stressors through plasmid-mediated heterozygosity. Many of the results are furthermore applicable to the related problem of the maintenance of incompatible plasmids in the same cell under selection for both.

**MSC Classification:** 92D15, 60J85

## 1 Introduction

Many bacteria carry, in addition to the bacterial chromosome, extrachromosomal genetic elements called plasmids, which replicate independently of the chromosome and often exist in multiple copies. The simplest plasmids have only the core backbone genes required to ensure their stable maintenance in the bacterial host, but many plasmids also carry genes that have important effects on the host phenotype (Garcillán-Barcia et al, 2011). Perhaps the most famous of these are antibiotic resistance genes, which pose a serious threat to the effectiveness of the clinical treatment of bacterial infections (Carattoli, 2013), but phenotypes contributed by plasmid-borne genes also include heavy-metal resistance, virulence, metabolism of novel carbon sources, and symbiotic interactions with other organisms, among many others (see e.g. Portnoy and Martinez, 1985; Silver, 1992; Beijersbergen et al, 1994; Yu et al, 1996; Anda et al, 2015; Wardell et al, 2021). The plasmids carried by a bacterium form an important part of the bacterial genome, and the contribution of plasmids must be considered in explaining and predicting bacterial evolution. In this context, much attention has been given to the ability of many plasmids to transfer horizontally between bacteria, infecting new hosts, in a process called conjugation (Falkow, 1975; Smillie et al, 2010); however, very many plasmids are transmitted only vertically, or depend on other plasmids for horizontal transfer (Coluzzi et al, 2022), and purely vertically transmitted plasmids can play an important role in bacterial adaptation.

The backbone genes of a plasmid are responsible for ensuring the stable maintenance of the plasmid by vertical transmission. This requires ensuring that both daughter cells receive copies of the plasmid at host cell division and limiting the possible reduction in host growth imposed by fitness costs of plasmid carriage. To this end, the number of copies of a plasmid per host cell is regulated by the plasmid itself to remain approximately constant; the plasmid copy number is therefore an intrinsic property of the plasmid-host system which plays an important role in the biology of the plasmid (Rodríguez-Beltrán et al, 2021). Although for some plasmids the copy number is kept quite low, at only one or a few copies (e.g. Gustafsson and Nordström, 1980; Casjens et al, 2000; Nordström, 2006; Ismail et al, 2014; Brovedan et al, 2019; Smith et al, 2021), many plasmids are found in their hosts in high numbers of copies, often in tens or sometimes even hundreds (e.g. Kiyosawa et al, 1993; Burian et al, 1997; Projan et al, 1987; Prangishvili et al, 1998; Anda et al, 2015; San Millan et al, 2016; Santos-Lopez et al, 2016). Interest in the evolution of these multicopy plasmids and their contribution to bacterial adaptation has recently increased, particularly due to their contribution to antibiotic resistance (San Millan et al, 2009; Gama et al, 2018). Existence in multiple copies per cell has multiple effects on the contribution of multicopy plasmids to bacterial evolution. The copy numbers of genes carried on a multicopy plasmid are also higher than those of genes on the (for many species haploid) bacterial chromosome, so genes on plasmids have both higher mutational input and dosage (these effects have been shown in experiments by, e.g., San Millan et al (2016)). The presence of the plasmid in multiple copies also provides the possibility of heterozygosity for loci on a multicopy plasmid which is not possible for loci on a haploid chromosome.

Once there are multiple variants of the plasmid in the cell, the process of plasmid segregation at host cell division will have important effects on the distribution of plasmid variants among cells in the population (San Millan et al, 2016; Ilhan et al, 2018; Santer and Uecker, 2020; Rodríguez-Beltrán et al, 2021). At host cell division, the copies of a plasmid in the host are distributed between the two daughter cells: for low copy number plasmids this is often aided by an active partitioning system to ensure each daughter gets at least one copy of the plasmid, while high copy number plasmids may simply rely on random distribution of the plasmids between daughter cells (Zielenkiewicz and Cegłowski, 2001). Random assortment of the plasmid copies at host cell division (i.e. segregation independent of the allele at the heterozygous locus) tends to eliminate heterozygosity, since every cell division has some nonzero probability of producing a homozygote daughter cell by chance, and the homozygotes are absorbing states of this process (see Figure 1A). An analogous situation appears when two different (multicopy) plasmids that share the same replication system reside within the same cell—the plasmid variants are said to be *incompatible*, since they will not be maintained together in the same cell line indefinitely, and over time separate lines containing either one or the other will emerge (Novick, 1987). Such segregative loss of one or the other plasmid was already observed and quantified in early studies on plasmid incompatibility; this loss can be counteracted by selection for maintenance of the plasmid variants together (see e.g. Uhlin and Nordström, 1975; Cullum and Broda, 1979; Rodriguez-Beltran et al, 2018).

**Fig. 1.**
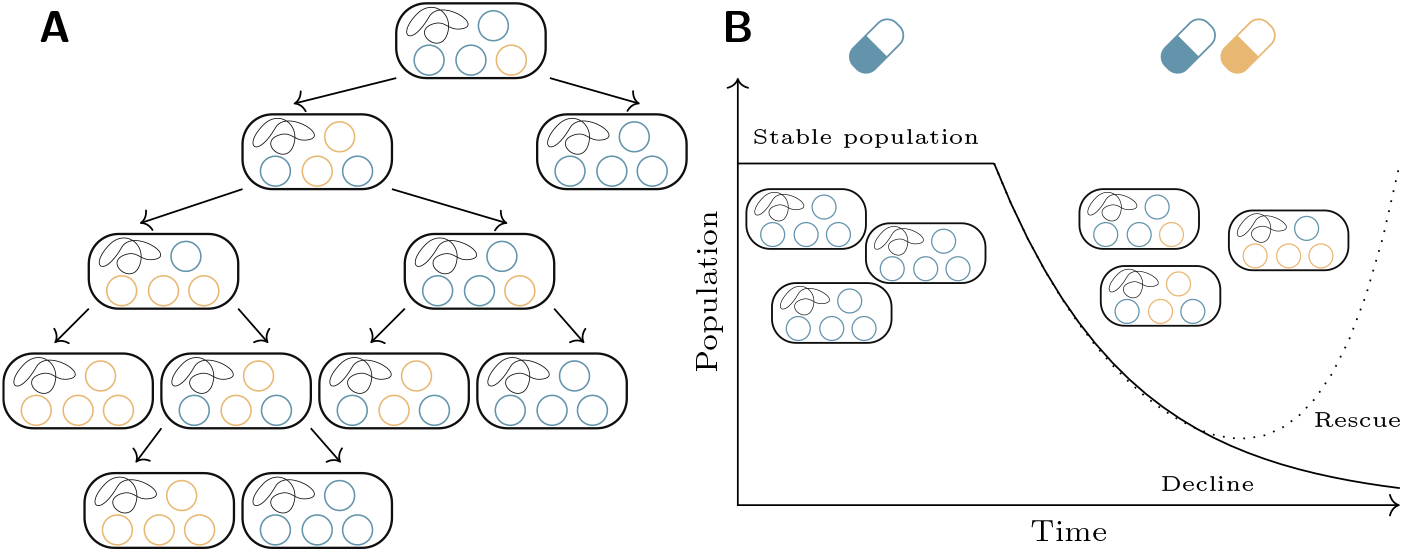
A model of the rescue of a bacterial population by plasmid-mediated heterozygosity. (A) The loss of plasmid-mediated heterozygosity due to segregation. From an initial heterozygote cell, random segregation results in the two plasmid types ending up in different, homozygous cells. (B) The evolutionary rescue scenario. An initial stable population carrying a plasmid with an adaptive gene on it is exposed to a sudden environmental change, which leads the population to decline. A mutation on the plasmid (orange plasmids) can rescue the population, but only if it is maintained together with the wild-type (blue) plasmid. Based on Figure 1 of Santer and Uecker (2020).

Driven by these empirical observations, early models were developed to describe the kinetics of the loss of heterozygosity in multicopy plasmids (or loss of incompatible plasmids) in the absence of selection under varying assumptions about the mechanisms of segregation and replication (Ishii et al, 1978; Novick and Hoppensteadt, 1978; Cullum and Broda, 1979). Recently, interest in the evolution of multicopy plasmids and their hosts has gained momentum (e.g. San Millan et al, 2016; Rodriguez-Beltranet al, 2018; Santer and Uecker, 2020; Rodríguez-Beltrán et al, 2021; Garoña et al, 2021; Hernandez-Beltran et al, 2022; Santer et al, 2022; Garoña et al, 2023). This includes experimental studies of multicopy plasmids with accompanying models: some of these focus on the fixation process of novel beneficial alleles on plasmids (Ilhan et al, 2018; Garoña et al, 2023), while others focus on the effect of selection for heterozygotes on maintaining plasmid variants together despite segregation (Rodriguez-Beltran et al, 2018). Recent modelling studies have also examined the effects of plasmid replication and segregation on the fate of novel beneficial alleles that appear on a single copy of a multicopy plasmid (Halleran et al, 2019; Santer and Uecker, 2020).

We here focus on scenarios of heterozygote advantage, where the optimal bacterial fitness is obtained by having multiple variants of a single plasmid carrying distinct alleles. This could be because, for example, the two alleles confer resistance to distinct antibiotics present in the environment. The models we present are of an *evolutionary rescue* scenario, of the kind introduced by Gomulkiewicz and Holt (1995) and shown in Figure 1B, extending previous models of rescue on plasmids (Tazzyman and Bonhoeffer, 2014; Santer and Uecker, 2020). In such a scenario, an environmental change exposes a population to new conditions to which it is maladapted, and the population therefore enters a demographic decline which would lead to extinction. However, if a novel mutation emerges which adapts individuals to the new conditions, and this mutation survives to spread in the population, the population can be rescued from extinction. Since this rescue depends on the mutation occurring and surviving the initial period in which it is rare in the population, rescue is inherently a stochastic process, and the key question is the probability that rescue will occur in a given population. When maintenance of both plasmid variants is required for rescue, rescue requires not only the establishment of the new mutation against the force of genetic drift, but also the persistence of both variants against the force of segregation at all times. It is intuitively clear, that such persistence is only possible if the fitness of heterozygote cells is large enough to bear the production of unfit homozygous cells. Based on a multitype branching process model, we determine the probability of evolutionary rescue and derive analytical conditions on the plasmid copy number and the fitness of heterozygous cells that need to be fulfilled for maintenance of heterozygosity, and thus rescue, to be possible at all. While we derive this condition in the context of evolutionary rescue, it holds more broadly for the maintenance of heterozygosity or of incompatible plasmids within the same cell.

## 2 The model

Consider a demographically stable population of bacteria which carries a plasmid present at a fixed copy number *n* in each bacterial cell. This population is then exposed to novel environmental conditions to which it is maladapted, and the population begins a demographic decline which will eventually lead to its extinction. Evolution, however, may rescue this population from extinction. A mutation might appear at a locus on the plasmid which, *if maintained together with the wild-type allele at that locus*, will adapt the population to the novel environmental conditions; that is, mutant *heterozygotes* can survive in the novel environmental conditions, but mutant and wild-type homozygotes are both maladapted. We assume that the population is homozygous for this particular plasmid before the environmental change; that is, that there is only one variant of the plasmid in the population, at least with respect to the locus of interest. These conditions constitute the evolutionary-rescue-due-to-heterozygote-advantage scenario we wish to describe in the model. We expect that heterozygote advantage of plasmid-borne loci could occur in many circumstances, but as a motivating example we consider a bacterial population resistant to antibiotic A due to a resistance gene located on a plasmid. If the population is exposed to antibiotics A and B simultaneously, then lack of resistance to antibiotic B will cause the population to decline. But if a mutation occurs on a plasmid that converts the A-resistance allele into a B-resistance allele, its host is now resistant to antibiotic B, and (provided that *n >* 1) the remaining wild-type plasmids continue to confer resistance to antibiotic A: therefore the host can survive and grow in the novel conditions. Its descendants are resistant to both antibiotics and the population might be rescued from extinction, provided that both plasmid types, and therefore both resistances, are maintained. Rodriguez-Beltran et al (2018) found very similar situation arose in an evolution experiment where a mutation in a plasmid-borne TEM-1 *β*-lactamase shifted it from being mostly effective against ampicillin to mostly effective against ceftazidime. More generally, this scenario could be particularly likely with *β*-lactam antibiotics, as common small mutations can alter the affinity of *β*-lactamases for different drugs (Farr et al, 2023). Such a combination of two closely-related antibiotics would usually not be used in a clinical context to treat a patient (although recent research has suggested that sequential treatment with different *β*-lactams—including both penicillins, to which the wild-type TEM allele provide resistance, and cephalosporins, to which mutant extended-spectrum TEM alleles provide resistance—may be more effective than previously thought: Batra et al (2021)). However, our scenario might arise in accidental environmental exposure of bacterial populations to antibiotics, for example from agricultural sources, where nothing prevents two related antibiotics from cooccurring.

Our model extends that of Santer and Uecker (2020) to the case of heterozygote advantage. Suppose that cells with *i* mutant plasmids (and therefore *n − i* wild-type plasmids) reproduce at a rate *λ*_*i*_ and die at a rate *µ*_*i*_. We assume throughout that the plasmid composition only affects the replication rate but not the death rate. Since the unit of time is arbitrary, we may fix *µ*_*i*_ = 1. If we then set *λ*_*i*_ = 1 + *s*(*n, i*), the function *s*(*n, i*) gives the net growth rate (or Malthusian fitness) of cells with *i* mutant plasmids out of *n*. Since maintenance of both wild-type and mutant plasmids is necessary for the population to persist, we have that *s*(*n*, 0) *<* 0 and *s*(*n, n*) *<* 0; beyond this, the choice of fitness function is free.

We will consider two different possible fitness functions. The simpler is

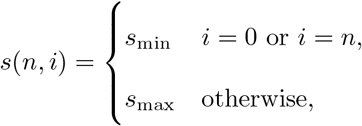

where *s*_min_ *<* 0 and *s*_max_ *>* 0. We shall call this the *dominant fitness function* (shown in Figure 2A), since a single copy of either allele confers the full fitness effect of that allele. It is also possible to imagine that dominance is intermediate or that there is a gene dosage effect, in which additional copies of a gene increase fitness. Combined with the necessity to maintain both alleles, this might produce a fitness function like

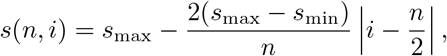

with *s*_min_ *<* 0 and *s*_max_ *>* 0 as before; this function linearly interpolates between a maximum fitness at an exact half-and-half mixture of wild-type and mutant plasmids, and a minimum fitness for the homozygotes (shown in Figure 2B). We shall call this the *peaked fitness function*. If there are gene dosage effects, *s*_max_ would increase with *n*; we do not consider this in our numerical examples, but all results for a given *n* also apply to this scenario.

**Fig. 2.**
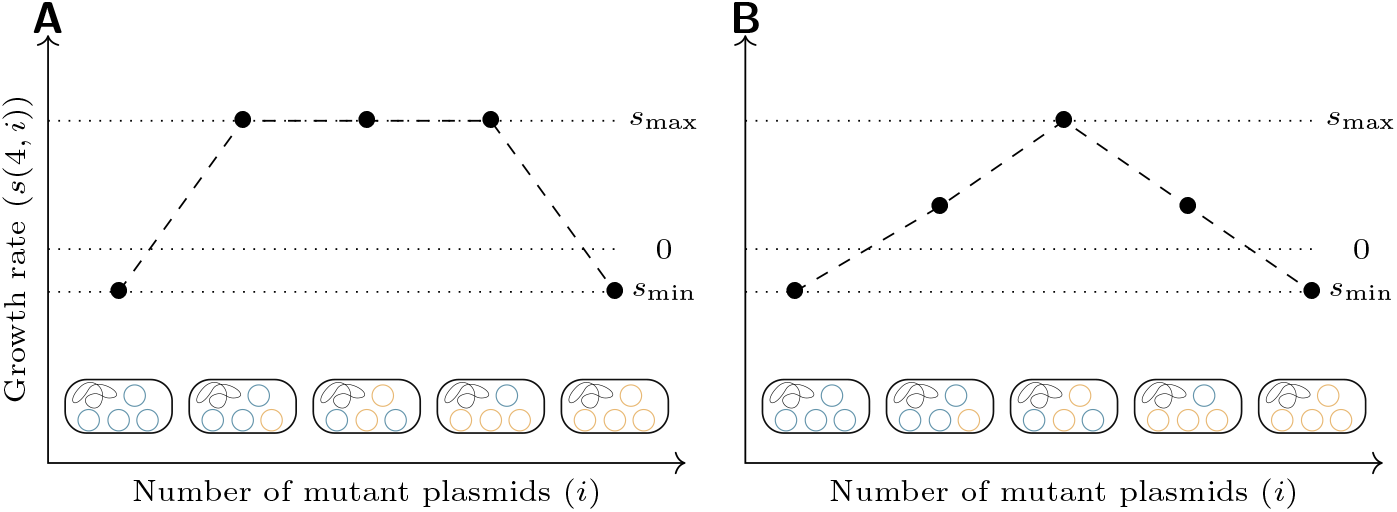
Examples of the two fitness functions considered, for *n* = 4: (A) the dominant fitness function; (B) the peaked fitness function.

We have assumed above that every cell has *n* copies of the plasmid of interest. To maintain this number, we assume the cell replicates its plasmids to a copy number of 2*n* before reproduction, and then segregates equal numbers into each daughter cell. At replication of wild-type plasmids, mutations happen with probability *u*, but we disregard this for the moment. We denote by *P* (*i* ⟶ {*j, k*}) the probability that when a cell with *i* mutant plasmids (out of *n* total plasmids) divides it produces daughters with *j* and *k* mutant plasmids (again out of *n* total). We will consider two models of replication, called *regular* and *random replication*, after the taxonomy of Novick and Hoppensteadt (1978).

In regular replication, every plasmid copy in the parent cell is replicated once, and then the copies are segregated randomly, but maintaining the copy number, into each daughter cell. The number of mutant plasmids in the daughter cell then has a hypergeometric distribution, giving a segregation probability function

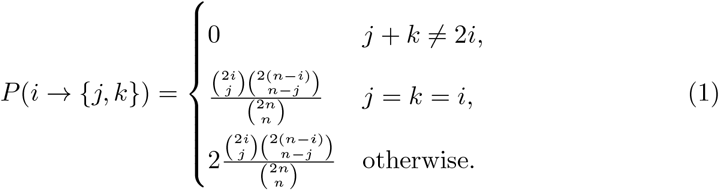

In random replication, each new plasmid to be replicated is randomly chosen from the pool of the initial plasmids together with the products of previous replications. This proceeds until there are 2*n* total plasmids, when the number of mutants will be distributed according to a Pólya urn scheme. These 2*n* plasmids are then randomly segregated into the daughter cells as before. The number of mutant plasmids in a daughter cell is then distributed as

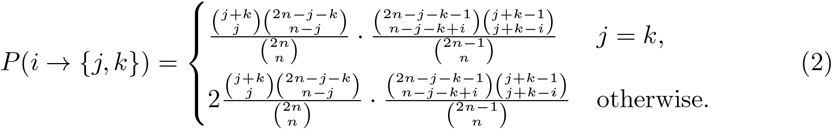

The first fraction in the expression is the same hypergeometric distribution for segregation as in the regular replication model, representing the probability that the *j* + *k* mutant plasmids produced by replication are divided into *j* in one daughter cell and *k* in the other. The second fraction is the probability of producing *j* + *k* mutant plasmids from *i* mutant plasmids during replication under the Pólya urn process model: for the derivation see appendix A.

For rescue to happen, a mutation must occur on the plasmid and be maintained together with the wild-type plasmid in the same cell indefinitely. To determine the probability of this occurring in a given population, we split the process into two parts. In the first, we look at the descendants of a single novel mutant, and determine the probability that these descendants will survive indefinitely rather than suffering stochastic loss. This is called the *establishment probability* of the mutant. We ignore additional mutations that might recurrently generate the mutant from the wild-type plasmid, which is a rare event in a small cell line. In the second part, we can then estimate the number of mutants which will occur in wild-type homozygous cells before the initial population goes extinct, and determine the probability that at least one of them will establish; this is called the *rescue probability*.

We have described our birth-death model thus far as a continuous-time multitype branching process, in which cells divide and die independently of each other, excluding in particular resource competition. Of course, a population cannot grow indefinitely. However, we are here considering a population that is, at least initially, declining due to maladaptation. Mutant cells are rare in the early stages of establishment in which we are interested and therefore independent from each other to a good approximation; this approximation is commonly made even in more complicated models and dates back to the very early calculations of establishment probabilities of beneficial alleles (Haldane, 1927). We further assume that we are far enough away from carrying capacity that we can also neglect competition with the declining wild-type population.

## 3 The establishment probability

### 3.1 Analysis

To determine the establishment probability for the descendants of a given bacterial cell, we first determine the extinction probability, the probability that the cell’s descendants will at some point go extinct. While our model was described above in continuous time, we transition to a discrete-time branching process for the analysis of the establishment probability. We can do this because we are only interested in the final outcome of the process—extinction or survival—and not in the timing of events.

A given cell will have either zero daughter cells (if it dies before reproducing), with probability *µ*_*i*_*/*(*λ*_*i*_ + *µ*_*i*_), or two daughter cells (if it reproduces), with probability *λ*_*i*_*/*(*λ*_*i*_ + *µ*_*i*_). Since the survival of the cell depends on its complement of plasmids, we will need to track the number of mutant and wild-type plasmids in each daughter cell. The extinction probabilities *Q*_0_, *Q*_1_, …, *Q*_*n*_ of the descendants of cells with, respectively, 0, 1, …, *n* mutant and *n, n −* 1, …, 0 wild-type plasmids are given by the least fixed point of the generating function of the distribution of daughter cells of a single cell (Sewastjanow, 1975, Folgerung V.1.1). This function is given by *f* (*z*_0_, …, *z*_*n*_) = (*f*_0_(*z*_0_, …, *z*_*n*_), …, *f*_*n*_(*z*_0_, …, *z*_*n*_)), where each component *f*_*i*_ is given by

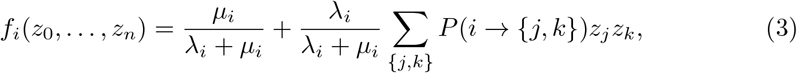

the sum being taken over all unordered pairs *{j, k}* of numbers of plasmids. The first term of this expression corresponds to the probability of immediate cell death, while the sum gives the probabilities of each possible pair of daughter cell types upon reproduction. The fixed point of the generating function is then the solution to the system of *n* + 1 equations (3) in *n* + 1 unknowns *z*_*i*_ = *Q*_*i*_, which can be solved numerically. This is also intuitive: the initial cell has either zero or two daughter cells. In the case that the initial cell has zero daughter cells, its descendants go extinct immediately (corresponding to the first term in the fixed point equation); in the case it has two, its descendants go extinct if and only if the descendants of both daughter cells go extinct (corresponding to the second term).

The fixed point gives us the probability that the descendants of a bacterial cell eventually go extinct, but what we really want is the probability that the descendants of a novel mutant never go extinct: the establishment probability. Under the regular replication model, this is very simple to determine: a novel mutant allele appears at first on a single plasmid in a single host cell, so the establishment of the mutant allele occurs only when the descendants of that cell do not go extinct, and

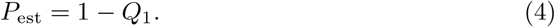

In the random replication model the situation is slightly more complex. When a mutation occurs during plasmid replication, the novel mutant plasmid is available to be replicated during the same bacterial generation. If it is replicated, there will be multiple mutant plasmids to segregate between daughter cells, and each daughter cell might get one or more mutant plasmids. Thus the establishment process of the mutation gets a head start, possibly starting from multiple mutant plasmids which may possibly be in two separate cells. The establishment probability becomes a quadratic form in the extinction probabilities of the individual types

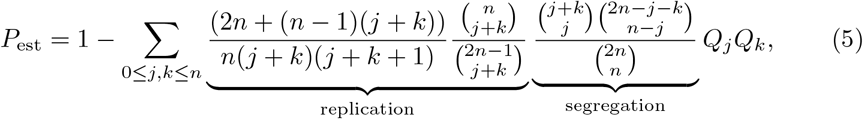

where the first factor inside the sum is the probability of going from no mutant to *j* +*k* mutant plasmids during replication, conditional on exactly one mutation occurring (for the derivation of this probability and its assumptions, see appendix A), and the second factor is the probability of the *j* + *k* mutant plasmids being segregated into daughter cells with *j* and *k* mutant plasmids.

### 3.2 Results

The establishment probabilities for the regular replication model with a dominant fitness function are shown in Figure 3A. Perhaps unsurprisingly, increasing the fitness of heterozygotes (*s*_max_) increases the establishment probability. More interesting is the effect of the copy number of the plasmid. At low copy number, the establishment probability increases with copy number: this is due to the loss of heterozygosity to segregation, which is most pronounced for small copy numbers. For larger copy numbers, the establishment probability stabilizes—indeed, a close examination shows that it begins to decrease for larger copy numbers, probably because at high copy numbers it takes more generations to go from a single mutant plasmid copy to an approximately equal number of wild-type and mutant plasmids, where the probability of producing a homozygote daughter cell is lowest (a similar effect was observed in the non-heterozygote-advantage case by Santer and Uecker (2020)).

**Fig. 3.**
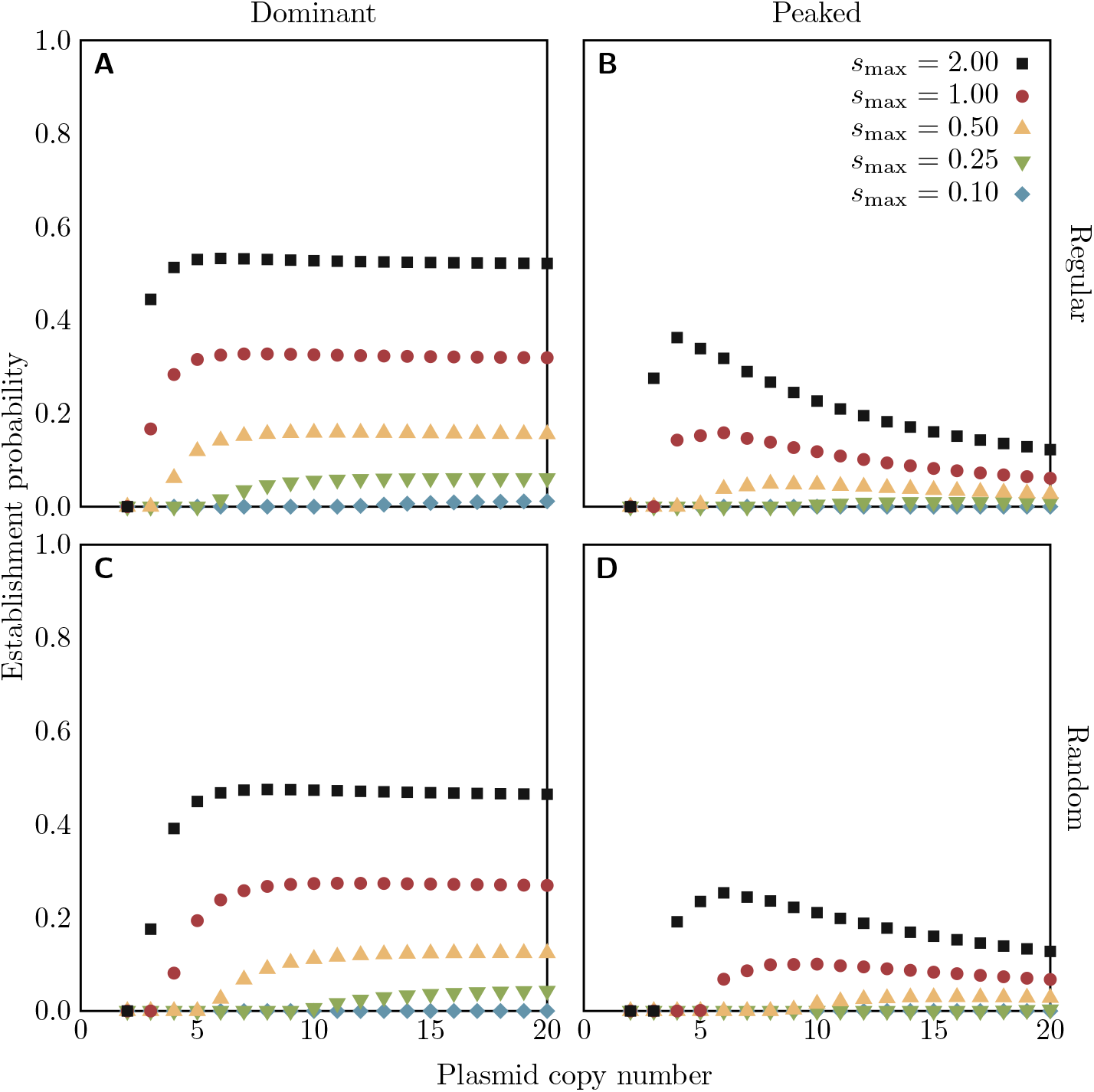
Establishment probabilities of rescue mutations on a plasmid of a given copy number in the heterozygote advantage scenario, under the regular (top row, from equation (4)) or random (bottom row, from equation (5)) replication assumption. Colours of points indicate the fitness *s*_max_ of all heterozygotes (with the dominant fitness function, left column) or the maximum fitness of heterozygotes (with the peaked fitness function, right column). For all cases, *s*_min_ = −0.1. The code for numerical calculation of establishment probabilities is available as supplementary material.

The establishment probabilities for the regular replication model with a peaked fitness function are shown in Figure 3B. The establishment probabilities are lower than with the dominant fitness function, and decline more precipitously for high copy numbers, since the fitness of most cell types, other than the perfectly balanced heterozygotes, has been reduced. In fact, for large enough copy numbers, the fitness of cells with positive but small numbers of one or the other plasmid type becomes negative.

The establishment probabilities for the random replication model are shown in Figures 3C (with the dominant fitness function) and 3D (with the peaked fitness function). The general trend is the same as under the regular replication model, but the establishment probabilities are all reduced. This is because random replication increases the rate of loss of heterozygosity to segregation relative to regular replication: the “rich-get-richer” behaviour of the Pólya urn scheme means that the proportion of mutant plasmids is likely to be more unbalanced after replication, and the probability of a homozygote daughter cell is increased.

For comparison, we performed stochastic computer simulations, which are in excellent agreement with equations (4) and (5) (see supplementary material).

Note that although the establishment probabilities appear to level off for the random replication model with dominant fitness function, in fact the establishment probability goes to zero for arbitrarily large copy number under random replication. This is because the mutant allele appears initially in a single copy: as the copy number increases, the probability of that mutant plasmid being selected for replication decreases. In the limit, the mutant plasmid is never replicated at all, and eventually the cell line containing it is lost to demographic stochasticity, and the mutant goes extinct.

## 4 The rescue probability

### 4.1 Analysis

Now let us consider the overall probability of rescue for the entire population. We follow the typical approach for approximating rescue probabilities from branching process models (used by, e.g, Orr and Unckless, 2008; Alexander and Bonhoeffer, 2012; Martin et al, 2013; Tazzyman and Bonhoeffer, 2014; Santer and Uecker, 2020): we assume that the initial population of homozygote wild-type cells is large, and that mutations are sufficiently rare that their occurrence and establishment can be treated as being independent. The first assumption allows us to treat the initial population of homozygote wild-type cells deterministically; it has an initial size *N*_0_ and growth rate *λ*_0_ *− µ*_0_ = *s*(*n*, 0) *<* 0, so its size at time *t* is given by

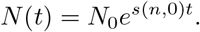

If the per locus mutation rate is *u*, then at each cell division the probability that a mutation occurs and will successfully establish is *unP*_est_ (we assume that at most one mutation occurs per cell division). Thus the instantaneous rate of occurrence at time *t* of mutations that will successfully establish is *unP*_est_*λ*_0_*N* (*t*). Our second assumption, of independence of mutations, means that successfully establishing mutations form an inhomogeneous Poisson process with intensity function *unP*_est_*λ*_0_*N* (*t*). The rescue probability is just the probability that this process has at least one event (i.e., at least one mutation occurs and successfully establishes), and is given by

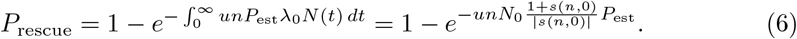

The Poisson process approximation also allows us to find the distribution of the time until the first successful rescue mutation occurs (conditional on one occurring at all): the time of occurrence *T* is distributed as

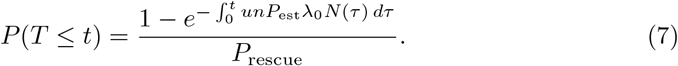

Our choice to set *µ*_*i*_ = 1 means that the unit of time here is the mean time to death of host cells. Note, however, that visible recovery of the population will in general occur much later: even after a successful mutation has occurred, the total population size still keeps decreasing initially, since the majority of cells are still maladapted wild-type cells.

### 4.2 Results

The rescue probabilities are shown in Figure 4. The major difference to the trends in the establishment probabilities in Figure 3 is that the rescue probability increases with copy number even when the establishment probability is stable or declining; this is because the increased mutational input with a greater copy number results in more mutants arising before wild-type population extinction, and thus a higher probability that at least one establishes; this effect turns out to be stronger than that of the decrease in the establishment probability. As for the establishment probability, we additionally performed stochastic computer simulations, which confirm the results (see supplementary material).

**Fig. 4.**
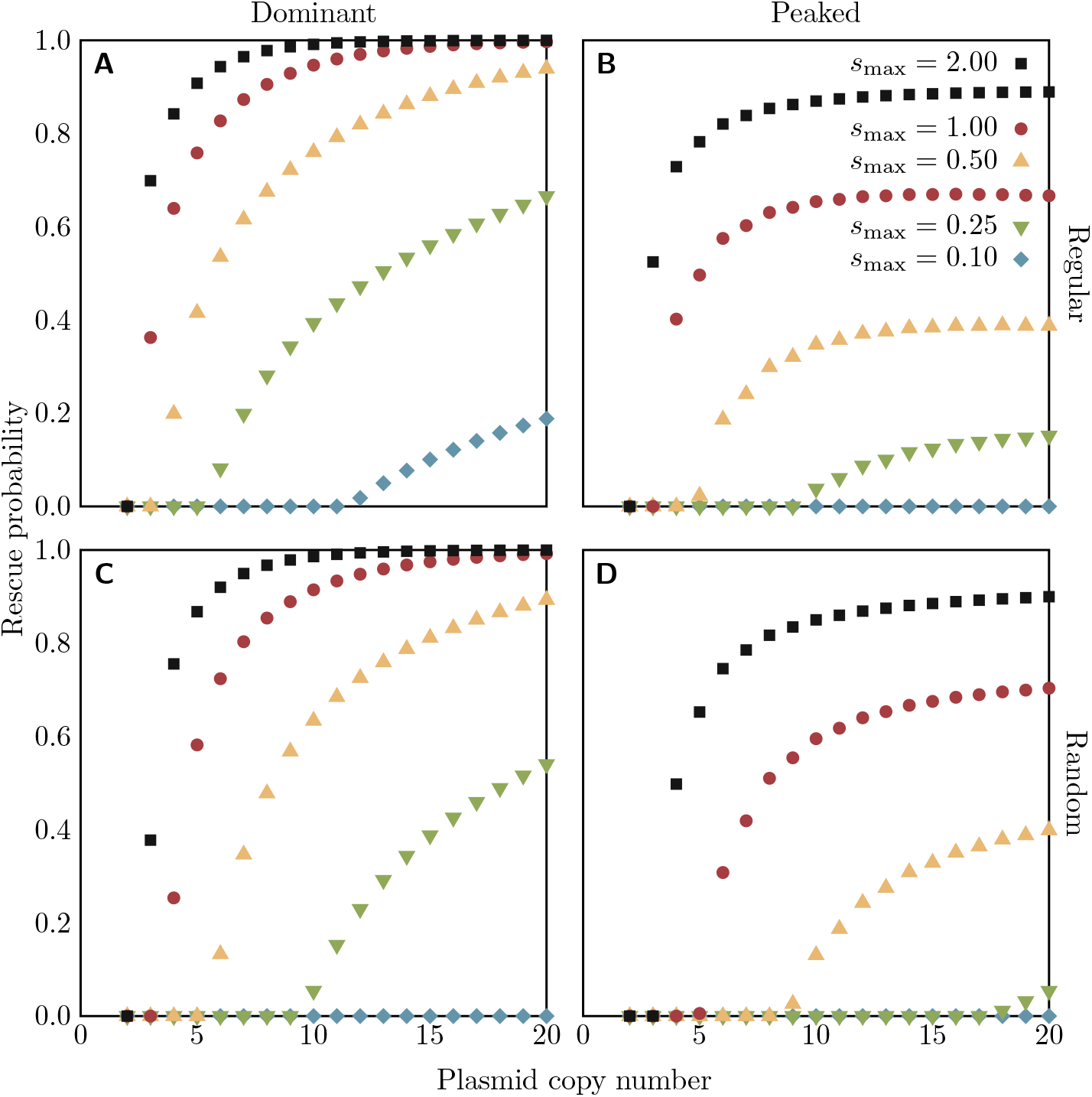
Probabilities (from equation (6)) of a bacterial population being rescued by a mutation on a plasmid of a given copy number in the heterozygote advantage scenario, under the regular (top row) or random (bottom row) replication assumption. Colours of points indicate the fitness *s*_max_ of all heterozygotes (with the dominant fitness function, left column) or the maximum fitness of heterozygotes (with the peaked fitness function, right column). For all cases, *s*_min_ = *−*0.1 and *uN*_0_ = 0.1. The code for numerical calculation of rescue probabilities is available as supplementary material.

Exemplary trajectories of the population size over the course of the rescue process from stochastic simulations following the Gillespie algorithm (Gillespie, 1976) are shown in Figure 5. Given rescue occurs, population recovery occurs earlier and is faster for higher copy numbers: population growth turns from negative to positive at earlier times, the population size at the minimum is larger, and subsequent growth is more rapid. We calculate the (long-run) growth rate of the rescued population for the dominant fitness function at the end of the next section.

**Fig. 5.**
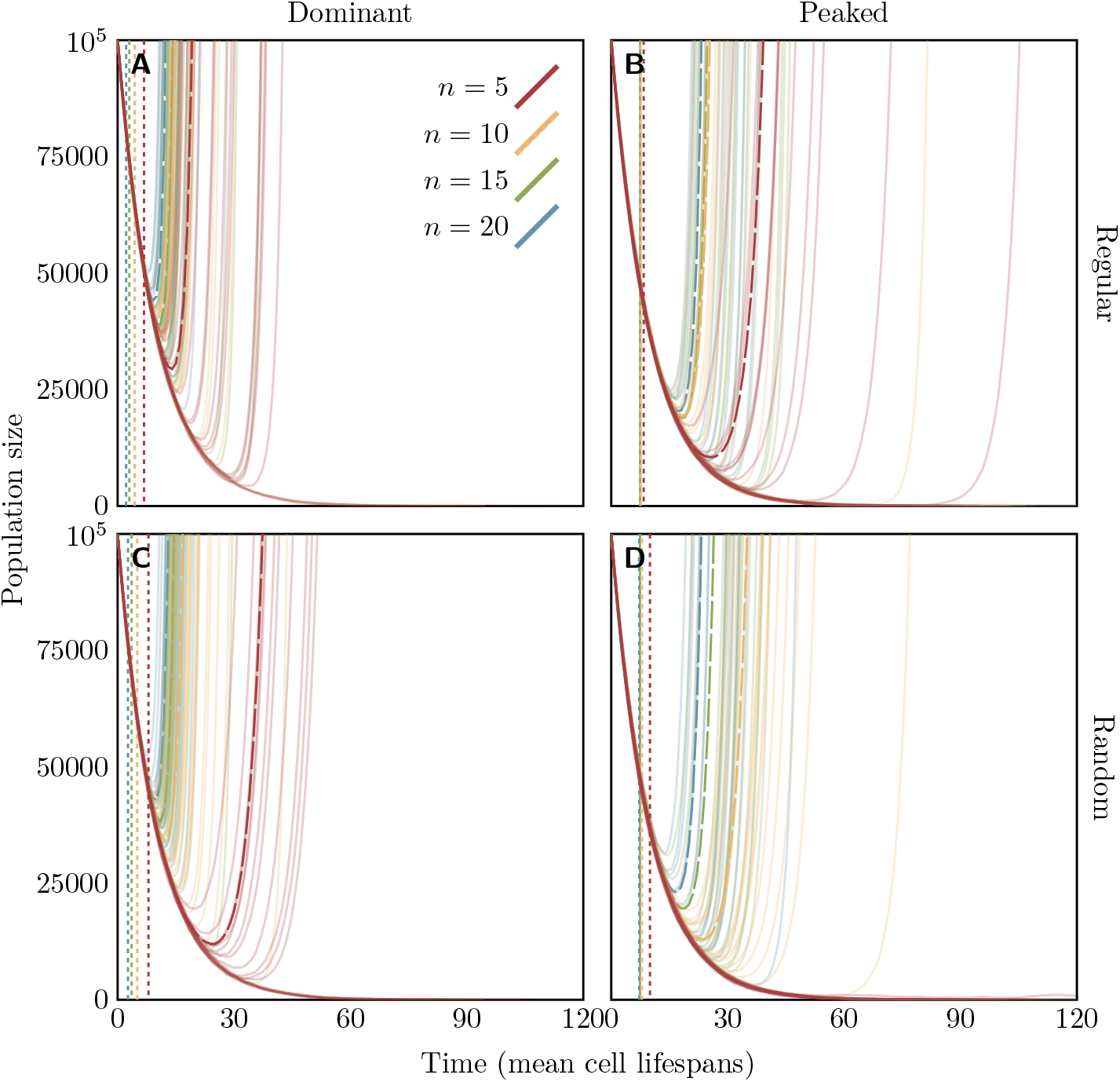
Simulated population size trajectories of a bacterial population being rescued by a mutation on a plasmid in the heterozygote advantage scenario, under the regular (top row) or random (bottom row) replication assumption. Time (on the *x*-axis) is measured in units of the mean lifespan of host cells 1*/µ*_*i*_. Colours of lines indicate the plasmid copy number: for each copy number, there are 20 simulated trajectories. The dashed curves show the mean of the simulations which did not go extinct; the dotted vertical lines indicate the mean time of occurrence of the first successful rescue mutation conditioned on rescue (from equation (7)). For all cases, *s*_max_ = 1.0, *s*_min_ = *−* 0.1, and *uN*_0_ = 0.1. For the peaked fitness function and random replication (panel D), the establishment probability with *n* = 5 is very small (although non-zero), and none of the 20 replicate populations survived. The code for the stochastic simulations is available as supplementary material.

## 5 Critical values of *n* and *s*_max_

Examining Figures 3 and 4 shows that not only does the effect of loss of heterozygosity during segregation cause the establishment and rescue probabilities to decline sharply at low copy numbers, but below a certain threshold (which depends on *s*_max_) establishment and therefore rescue becomes impossible (*P*_rescue_ = *P*_est_ = 0). We can, in fact, describe analytically the relation between the heterozygote advantage *s*_max_ and the copy number *n* which is required to hold in order for establishment to be possible with the dominant fitness function, in both the regular and random replication models. We take the opportunity here to give names to the probabilities that a single bacterial cell reproduces before it dies, which will be important in deriving the analytical relations. With the dominant fitness function, there are only two distinct values of this probability,

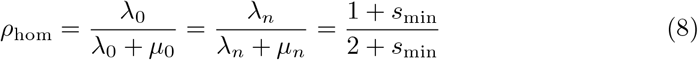

for homozygote cells and

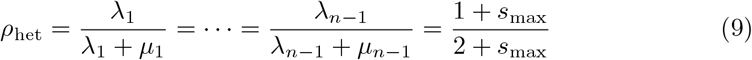

for heterozygotes.

### Theorem 1.

*In the model with regular replication and a dominant fitness function, the establishment probability (and therefore the rescue probability) is nonzero if and only if n ≥* 2 *and only if n ≥ 2 and*

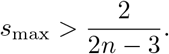

*Proof*. As in the derivation of the establishment probability, we consider the branching process in discrete time. Let *M* be the matrix the *i, j* element of which is the expected number of daughter cells with *j* mutant plasmids from a cell with *i* mutant plasmids. In the model with regular replication and a dominant fitness function, the elements of this matrix have the form

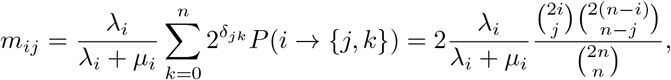

where we have substituted the definition of *P* (*i ⟶ {j, k}*) for regular replication given in equation (1). By a general result of branching process theory, a discrete-time multitype branching process goes extinct with probability one if and only if it has no final classes and the largest eigenvalue of *M* is less than or equal to one (Sewastjanow, 1975, Satz V.2.5). The first condition is a technical one, to exclude the degenerate case where there is no stochasticity in the branching process and the population stays at a fixed size: a final class is a subset of the types in the multitype branching process such that an individual of one of those types will have, with probability one, exactly one offspring and that offspring will be of one of the types in the class. In our case, there can be no final classes because every individual has a nonzero probability of having no offspring. Therefore, there is a nonzero probability of establishment if and only if *M* has an eigenvalue greater than one.

To find the eigenvalues of *M*, we factor it as a product *M* = 2*RP*, where *P* is the matrix with *i, j* element 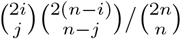 and *R* a diagonal matrix containing the probabilities of reproduction, then find the eigenvalues of each factor individually—it will turn out, luckily, that all the eigenvalues of *M* are products of the eigenvalues of these factors. Finding the eigenvalues of *R* is easy, since it is diagonal: in fact, it has only two distinct eigenvalues, which are given by the two possible probabilities *ρ*_hom_ and *ρ*_het_ of reproduction for a cell with the dominant fitness function given in equations (8) and (9). The eigenvalue *ρ*_hom_ has an eigenspace consisting of all of the “pure homozygote” vectors, that is those with nonzero entries only in the first and last component, and the eigenvalue *ρ*_het_ has an eigenspace consisting of the “pure heterozygote” vectors (those with zero first and last components).

The eigenvalues of *P* were calculated by Schensted (1958); we present here an elaboration of her argument. The trick will be to find a basis of ℝ^*n*+1^ in which the matrix that represents the same linear transformation represented by *P* in the standard basis is upper triangular, which makes reading off the eigenvalues trivial (in other words, we show that *P* is similar to certain upper triangular matrix; Strang (2009, § 6.6)). This new basis will arise from considering the vector space ℝ[*X*] of polynomials in an arbitrary unknown *X* with real coefficients. The map *T* : ℝ[*X*] *⟶* ℝ^*n*+1^ that takes a polynomial *p*(*X*) to the vector

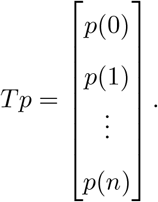

is clearly linear, and if we restrict the domain to polynomials of degree at most *n*, it becomes an isomorphism (since any *n* + 1 points define a polynomial of degree at most *n*). For any integer 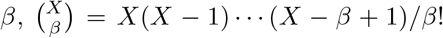 is a polynomial in *X* of degree *β*, and the sequence 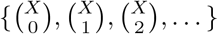 forms a basis for ℝ[*X*]; moreover, if we truncate the sequence at 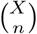, we get a basis for the subspace of polynomials of degree at most *n*, which we can pass through the isomorphism *T* to obtain a basis for ℝ^*n*+1^.

Looking carefully at the binomial coefficient identity

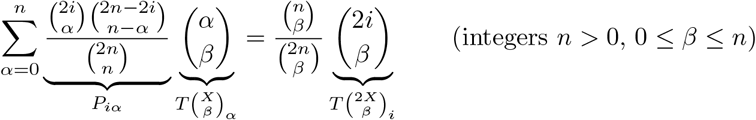

(proven in appendix A), we see that it is the *i*-th component of the equation

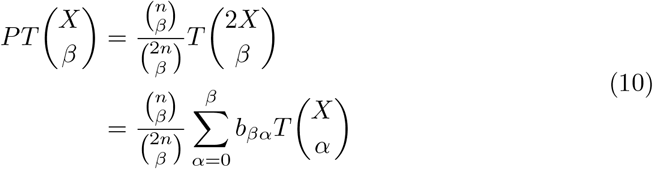

where the second equality follows from the fact that 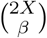 is a polynomial of degree *β*, and so must itself be a linear combination of 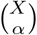 with *α ≤ β*.

From equation (10), we can see that 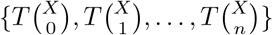 is the basis we are looking for: the linear transformation represented by *P* in the standard basis is represented in this basis by an upper triangular matrix with diagonal elements 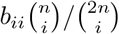. Since the eigenvalues are a property of the linear transformation, not its matrix representation in a particular basis, the eigenvalues *χ*_0_, …, *χ*_*n*_ of *P* are exactly these diagonal elements. To calculate *b*_*ii*_, we note that the coefficient of *X*^*i*^ in the polynomial 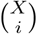 is 1*/i*!, while in the polynomial 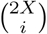 it is 2^*i*^*/i*!. None of the 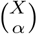 for *α < I* can contribute an *X*^*i*^ term, so it must be the case that *b*_*ii*_ = 2^*i*^. Thus we have that

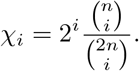

The ratio *χ*_*i*+1_*/χ*_*i*_ is less than or equal to 1, showing the sequence of eigenvalues is nonincreasing. The first three values are *χ*_0_ = 1, *χ*_1_ = 1, and 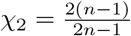.

Finally we get to the eigenvalues of *M* itself. In general, of course, it is not the case that the eigenvalues of a product of matrices are the products of the eigenvalues of the factors. However, if a pair of eigenvalues of the two factors share a common eigenspace, then their product is an eigenvalue of the product. This condition is not true for all the pairs of eigenvalues of *R* and *P*, but it *is* true for *ρ*_hom_ (an eigenvalue of *R*) and 1 (an eigenvalue of *P*). Recall that the *ρ*_hom_-eigenspace of *R* consists of the “pure homozygote” vectors: these are also exactly the *left* eigenvectors of *P* corresponding to the eigenvalue 1 (this can be seen from the fact that the first and last rows of *P* have a one on the diagonal and zeros everywhere else). Therefore 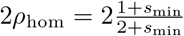 is an eigenvalue of *M* = 2*RP* (with multiplicity 2). But since *s*_min_ *<* 0, this eigenvalue is always less than 1, and we can ignore it: either it is not the dominant eigenvalue, in which case our conclusion does not depend on it, or it is the dominant eigenvalue, in which case the next largest eigenvalue will also be less than 1, and our conclusion will be the same as if that were the largest eigenvalue.

We now turn to the remaining eigenvalues of *M*. Let *χ ≠* 1 be another eigenvalue of *P* and *x* a corresponding *left* eigenvector. We now show that 2*ρ*_het_*χ* is an eigenvalue of *M*. Unfortunately, it will not be so easy this time: *x* does not have to lie in the *ρ*_het_- eigenspace of *R*. However, we can split *x* into a pure homozygote component *x*_hom_, which is the component of *x* in the *ρ*_hom_-eigenspace of *R*, and a pure heterozygote component *x − x*_hom_, which lies in the *ρ*_het_-eigenspace. Of these three vectors, *x* is a (left) eigenvector of *P* (by definition), and *x − x*_hom_ is an eigenvector of *R* (by construction), but *x*_hom_ is an eigenvector of *both:* this enables us to construct a vector that will be a left eigenvector of *RP* corresponding to *ρ*_het_*χ*, namely *ρ*_het_(1 *− χ*)*x*_hom_ + (*ρ*_hom_ *− χρ*_het_)(*x − x*_hom_). To see it is an eigenvector of *RP*, note that

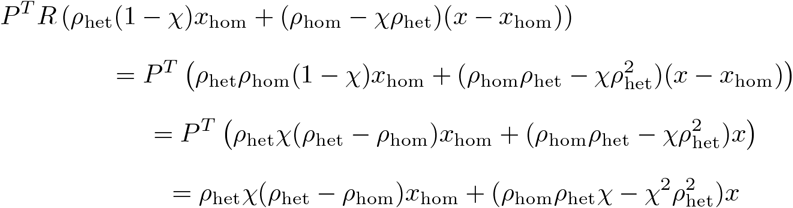

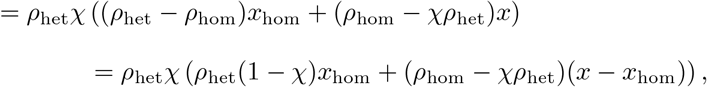

where the first equality uses that *x*_hom_ and *x−x*_hom_ are eigenvectors of *R* and the third uses that *x* and *x*_hom_ are left eigenvectors of *P*. From the eigenvalues *χ*_2_, *χ*_3_, …, *χ*_*n*_ of *P* we obtain in this way the remaining *n −* 1 eigenvalues of *M* = 2*RP*, and have thus found all eigenvalues of *M*.

Since 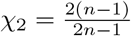 is the largest of the remaining eigenvalues of *P*, to check if there is at least one eigenvalue of *M* greater than 1 we need only check if 2*ρ*_het_*χ*_2_ *>* 1. We have that

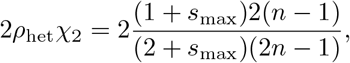

which is greater than one if and only if 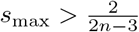

### Theorem 2.

*In the model with random replication and a dominant fitness function, the establishment probability (and therefore the rescue probability) is nonzero if and only if n ≥* 2 *and*

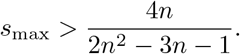

*In addition, in the long term limit, the heterozygote types are all equally abundant*.

*Proof*. Let *M* once again be the matrix the *i, j* element of which is the expected number of daughter cells with *j* mutant plasmids from a cell with *i* mutant plasmids. In the model with random replication and a dominant fitness function, the elements of this matrix have the form

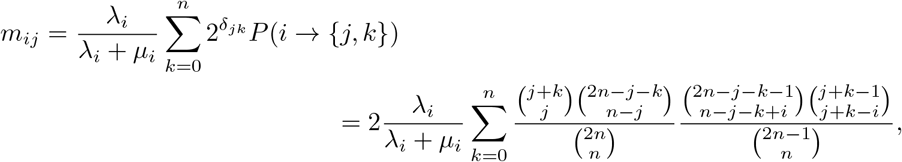

where we have substituted the definition of *P* (*i ⟶ {j, k}*) for random replication from equation (2). As in the proof for the regular replication case, we use general branching process theory (Sewastjanow, 1975, Satz V.2.5) to determine that there is a nonzero probability of establishment if and only if *M* has an eigenvalue greater than one.

The elements of the first and last rows of *M*, the numbers *m*_0*j*_ and *m*_*nj*_, must all be zero except the diagonal elements *m*_00_ = *m*_*nn*_ = 2*ρ*_hom_. This can be seen either from biological considerations—a homozygote cell will have only homozygote daughters—or by substitution into the definition of *m*_ij_ —if *i* = 0, then 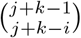 will be zero unless *j* = *k* = 0, in which case it is 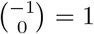, and if *i* = *n*, then the same is true of 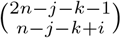. This means *M* has the structure

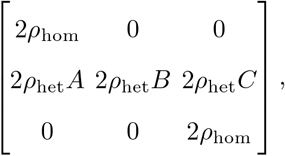

where *B* is an (*n −* 1) *×* (*n −* 1) matrix and *A* and *C* are *n −* 1 component column vectors. Then 2*ρ*_hom_ is an eigenvalue of *M* ; this eigenvalue, however, is always less than one, since *s*_min_ *<* 0. Every other eigenvalue of *M* will be 2*ρ*_het_ times an eigenvalue of *B*, and its corresponding left eigenvectors are of the form [*x*_0_ *v x*_1_], where *v* is a left eigenvector of *B*.

We now show that (*n −* 1)(2*n* + 1)*/*(*n* + 1)(2*n −* 1) is such an eigenvalue of *B* with acorresponding left eigenvector *v* with all components equal to 1 (the identification of this eigenvalue is due to Novick and Hoppensteadt (1978)). The *j*th element of *vB* is

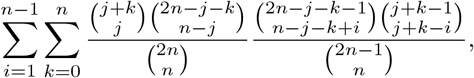

which by equation (A2) in appendix A is equal to (*n −* 1)(2*n* + 1)*/*(*n* + 1)(2*n −* 1); so *vB* = (*n −* 1)(2*n* + 1)*/*(*n* + 1)(2*n −* 1)*v*. This must also be the largest eigenvalue of *B*: *B* is a nonnegative irreducible matrix (since a cell with 1 *≤ i ≤ n −* 1 mutant plasmids can produce a descendant cell with 1 *≤ k ≤ n −* 1 mutant plasmids after at most |*i − k*| generations, *B*^*n−*1^ is a positive matrix), so by the Perron-Frobenius

Theorem (Frobenius, 1912; Sewastjanow, 1975, Satz IV.5.2) its only eigenvalue with a strictly positive eigenvector is the largest eigenvalue.

Thus 2*ρ*_het_(*n −* 1)(2*n* + 1)*/*(*n* + 1)(2*n −* 1) is the largest eigenvalue of *M* (other than possibly 2*ρ*_hom_) and its corresponding left eigenvector has all heterozygote types equally abundant. By simple rearrangement, we have that

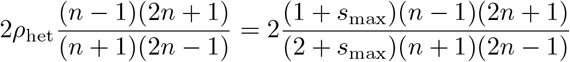

is greater than one if and only if 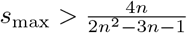.

Although we have given the conditions in the statements of the theorems as thresholds on *s*_max_, they can be equivalently restated as thresholds on *n* necessary for establishment: for regular replication, the threshold is 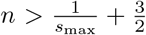, and for random replication it is 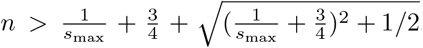. Note that as a direct consequence of these two theorems, if establishment is impossible in the regular case for a given *n* and *s*_max_, then is also impossible in the random case (the threshold *s*_max_ is always larger in the regular case, since 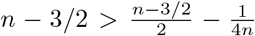). We have derived the threshold condition only for the dominant fitness function: we relied on having one common reproduction probability *ρ*_het_ for all heterozygote types, which made the interaction between host reproduction and the combinatorics of plasmid replication and segregation particularly simple.

With the insights obtained in this section, we can now also determine the exponential growth rate of the population, once rescued. For this, we switch back to the birth-death process in continuous time. At long enough times, growth is well described deterministically by a system of ODEs. This system will be linear, with a matrix of coefficients given by Λ(2*P − I*) *− D*, where *P* has the same meaning as in the proof of Theorem 1, the combinatorial part of the expected progeny matrix, Λ is a diagonal matrix of birth rates *λ*_*i*_, *I* is the identity matrix, and *D* is the diagonal matrix of death rates, which by assumption are the same for all cell types and given by *µ*_*i*_ = 1. By the same argument as in the proof of Theorem 1, the eigenvalues of Λ(2*P − I*) are products of an eigenvalue of Λ and an eigenvalue of 2*P − I* (the arguments in the proof of Theorem 1 also apply to random replication). The largest eigenvalue, which determines the growth rate of the rescued population is (1 + *s*_max_)(2*χ −* 1) *−* 1, where *χ* is the largest eigenvalue of *P* other than 1: this is 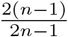 for regular replication and 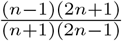 for random replication, where the latter result is obtained with the same reasoning as applied to the matrix *M* in the proof of Theorem 2.

## 6 The final distribution of cell types

If establishment occurs, we can ask what the relative abundances of cells with different numbers of mutant plasmids will be. The distribution of the number of mutant plasmids in a cell in the infinite time limit can be found for the discrete-time branching process from the expected progeny matrix *M*, with the expected number of daughters with *j* mutant plasmids from a cell with *i* mutant plasmids as its *i, j* element. If establishment is possible (*P*_est_ *>* 0) then *M* has a dominant eigenvalue, and a corresponding left eigenvector to that eigenvalue gives the distribution of cell types in the long run (Mode, 1971, Theorem 4.1). This eigenvector can be found numerically.

Examples of the final stable distribution of the number of mutant plasmids per cell under the four models considered (regular and random replication and dominant and peaked fitness functions) are shown in Figure 6. Even after establishment, considerable numbers of homozygote cells are present in the population, because of their constant replenishment by segregation. Heterozygotes are more abundant under regular replication, where there is a lower rate of loss of heterozygosity to segregation than with random replication, and with the dominant fitness function where the intermediate heterozygote types have higher fitness than with the peaked fitness function. Note that in the model with random replication and dominant fitness function, all heterozygote cell types are equally abundant in the long-term limit (as shown by Theorem 2).

**Fig. 6.**
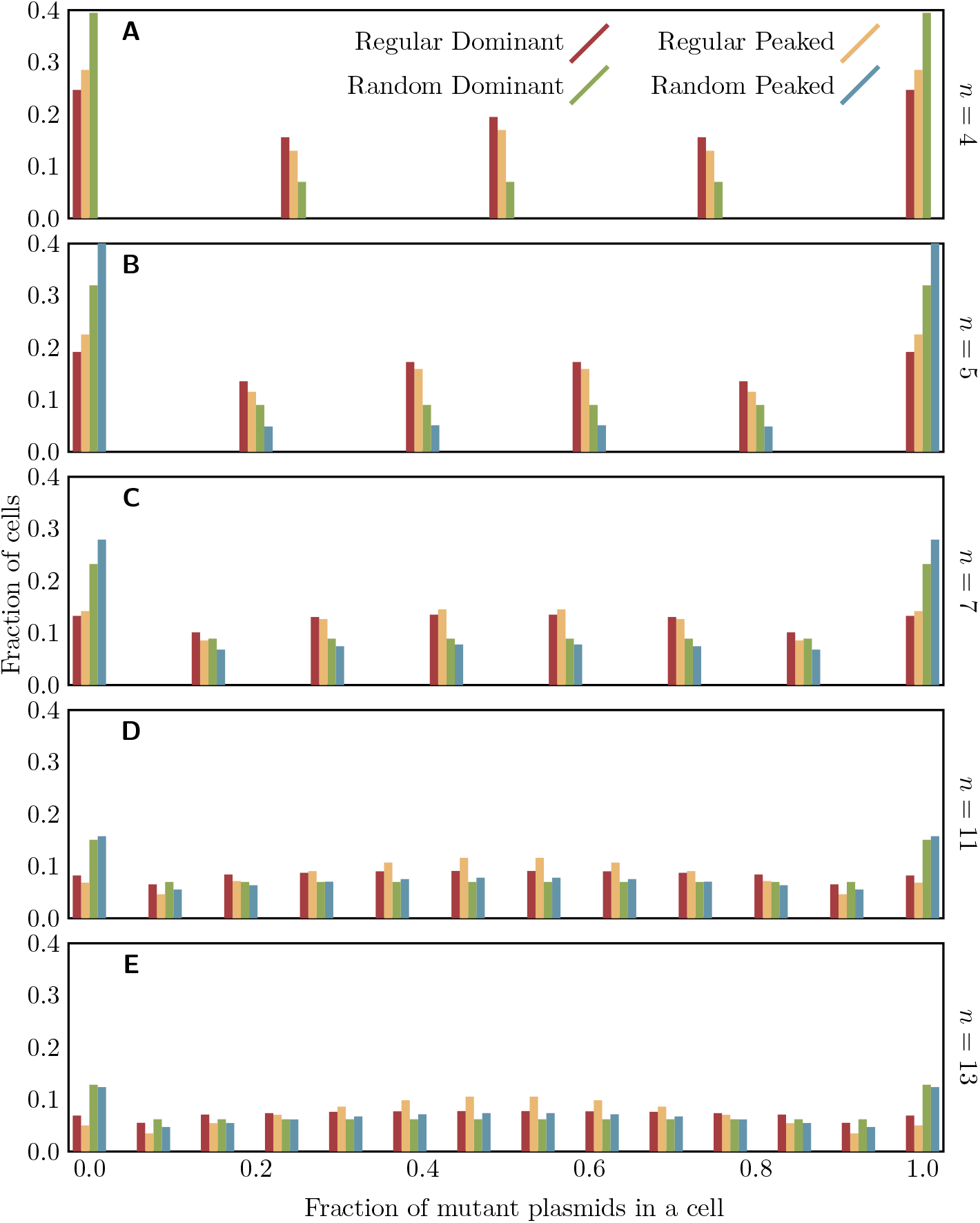
The final distribution of the proportion of mutant plasmids in a cell for plasmids of given copy numbers, under each of the four combinations of replication assumption and fitness function considered, and with *s*_min_ = 0.1 and *s*_max_ =− 1.0. The eigenvector corresponding to the leading eigenvalue of *M* was calculated numerically (code available as supplementary material); note that for *n* = 4 with random replication and the peaked fitness function, the population goes extinct with probability one, and there is no well-defined final distribution.

## 7 Cointegration

As we have seen, segregation plays an important role in limiting the possibility of the population being rescued. One process which alters the rate of segregation is the fusion of plasmids to form cointegrates. When plasmids fuse, the resulting multimer has multiple copies of the plasmid backbone, including multiple copies of the copy number control system, so we expect that an *m*-mer still “counts against” the copy number *m* times, even though it is only one molecule (Chiang and Bremer, 1991; Summers and Sherratt, 1984; Summers et al, 1993). Cointegration can eliminate the possibility of losing heterozygosity to segregation in the descendants of cells in which it occurs (experimentally observed by Hülter et al, 2020; Garoña et al, 2021): if plasmids with distinct alleles fuse, the resulting cointegrate provides the heterozygote phenotype on its own and can no longer be separated by segregation (unless the cointegrate is resolved into independent plasmids again). But conversely, if plasmids with identical alleles fuse, the number of independently segregating units in the cell is reduced, and the rate of loss will increase. We now present an extension of the model to include cointegration. We have chosen to use a very simplified cointegration model, with no resolution of multimers, only regular replication and the dominant fitness function, and novel mutations only appearing in cells with all monomers, to provide a comparison to the model without cointegration without a great deal of complexity.

### 7.1 The cointegration model

In a model which incorporates cointegration, there are a much larger number of possible cell types, since not only are there multiple loci that may have one allele or the other, but also these loci may be distributed among plasmids in a variety of ways. In the following, a cell type will be written in the form `(AA)(B)(AB)`, where each letter represents a locus carrying one of the two alleles, `A` and `B`, that must be maintained for the population to survive, loci within a pair of parentheses are on the same plasmid, and the total number of loci is fixed at the copy number *n*. In the example, the cell thus contains a cointegrate of two plasmids with allele A, a plasmid with allele B, and a cointegrate of a plasmid with allele A and a plasmid with allele B. Each cell type *a* then has a death rate *µ*_*a*_, which will be fixed to 1 as before, and a reproduction rate *λ*_*a*_, which is determined by the number of mutant and wild-type alleles in the cell (irrespective of their distribution on different plasmids). Incorporating cointegration into our model requires slight changes to the reproduction probabilities to account for the greater number of cell types, and adds a new process of cointegration which needs to be modelled.

#### 7.1.1 Reproduction

The reproduction probabilities are determined by the regular replication model used previously. The probability that a dividing cell of type *a* has daughters of types *b* and *c* will be denoted *P* (*a* ⟶ {*b, c*}) All plasmids are duplicated and then divided into two subsets having the same copy number *of the copy number control system*. This last part is important because now plasmids contribute differently to the total copy number. This means that some assortments of plasmids into daughter cells are no longer possible: for example, an `(AAA)(B)` cell can only have daughters identical to itself, since a `(B)(B)` cell has too few and an `(AAA)(AAA)` cell too many copies of the copy number control system. The probabilities *P* (*a* ⟶ {*b, c*}) are calculated by enumeration of the possible pairs of daughter cells and the number of ways to produce each one.

#### 7.1.2 Cointegration

The cointegration rate is divided into two parts: the overall probability of a cointegration event occurring in a given cell, which is the biological part, and the probability, conditional on cointegration occurring, that two particular plasmids fuse, which is purely combinatorial. The probability, conditional on a fusion occurring, that it takes the cell from type *a* to type *a*^*′*^ is denoted *P* (*a ⇒ a*^*′*^). This probability is the fraction of all pairs of plasmids in the *a*-cell that are pairs of the two types that need to fuse for this transition to occur. For example, 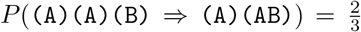 (there are 3 pairs of plasmids in the original cell, of which two are `{(A), (B)}` pairs). We only allow for fusion of plasmids but not for the resolution of cointegrates, therefore *P* ((AA)(A)(B) *⇒* (A)(A)(AB)) = 0.

The probability of cointegration occurring in one generation in cells of type *a* is denoted *κ*_*a*_. In the model presented here, this will be simply a constant *κ*, except in cells that have only a single large multimer, where it is 0 because no further cointegration can take place. We assume that at most one cointegration event occurs in each cell cycle.

### 7.2 Analysis and results

Combining the reproduction and resolution models, the extinction probability *Q*_*a*_ of type *a* is given by

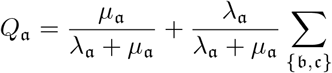

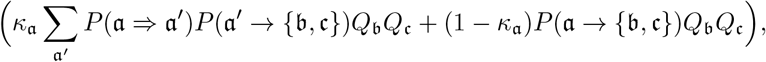

where the sums are taken over all types or unordered pairs of types. Note that this expression is equally compatible with selection happening before or after the fusion of plasmid copies, since the values of *λ*_*a*_ and *µ*_*a*_ are determined only by the alleles present and not their distribution between plasmid copies. Since we are using the regular replication model and make the assumption that the mutation first appears in a cell with only monomers present, the establishment probability is then given by

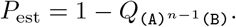

Establishment probabilities for the cointegrate model are shown in Figure 7. An important qualitative difference is that for non-zero *κ*, the establishment probability no longer exhibits a threshold effect: establishment is theoretically possible for any value of *s*_max_.

**Fig. 7.**
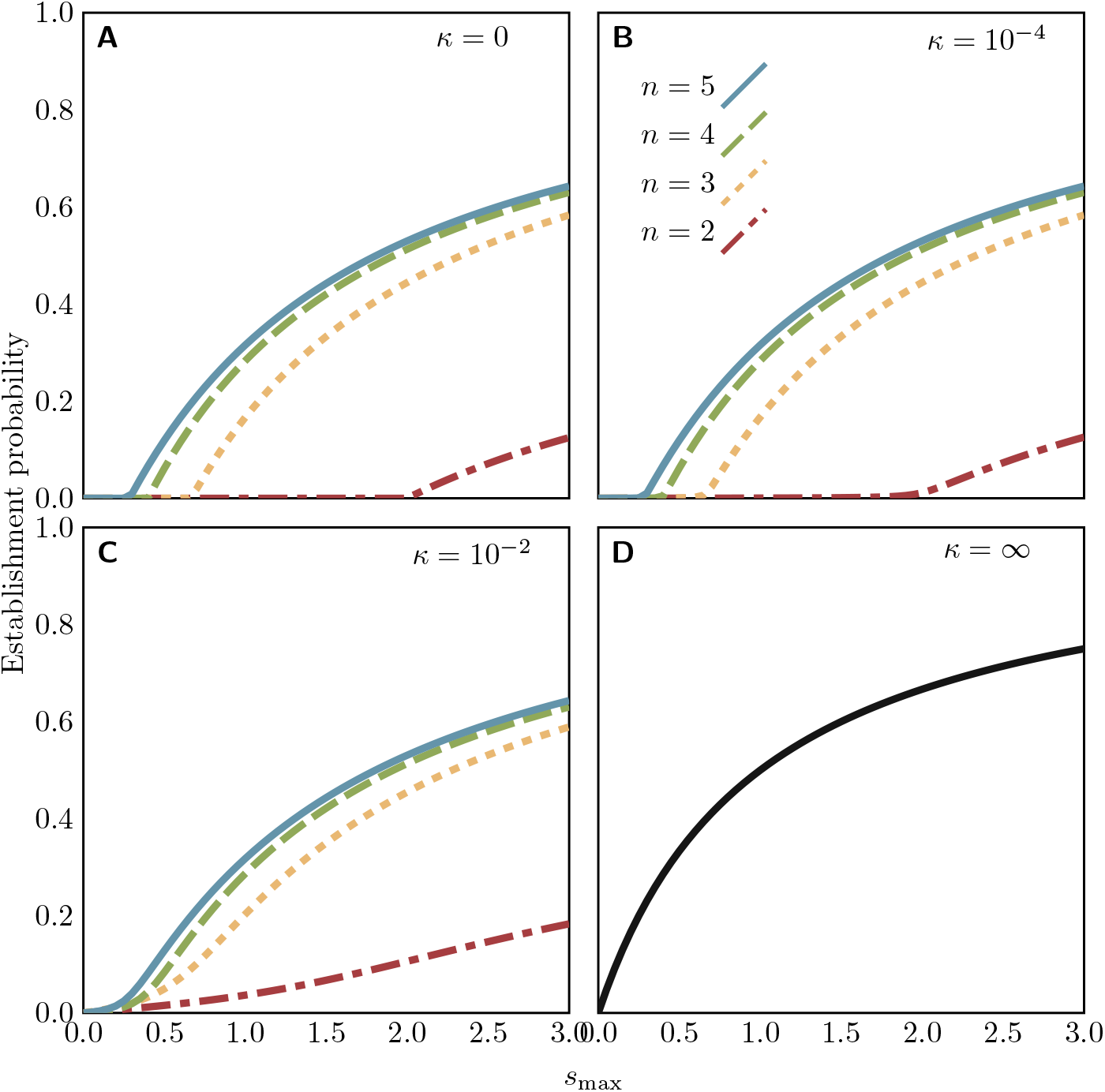
Establishment probabilities of rescue mutations on possibly-cointegrating plasmids of a given copy number, for varying values of the heterozygote fitness *s*_max_ and the cointegration probability *κ* (for all cases, *s*_min_ = *−* 0.1). The *κ* =∞ panel corresponds to a model in which the mutation is immediately followed by the fusion of all plasmids into a single multimer, and thus is the theoretical maximum establishment probability. The code for numerical calculation of establishment probabilities is available as supplementary material.

## 8 Discussion

Multicopy plasmids are a frequent component of bacterial genomes, where they may be cryptic passengers or play an important role in bacterial adaptation and evolution. One way in which they can play such a role is through their inherently polyploid nature: loci on a multicopy plasmid can exhibit heterozygosity which is not usually possible for loci on the (for many species) haploid chromosome. Here we have examined one scenario in which heterozygosity of plasmids may provide an adaptive advantage to their hosts, by enabling rapid evolution in scenarios of heterozygote advantage.

It is apparent that a key factor in determining the fate of heterozygosity on plasmids is plasmid segregation at host cell division. The effects of segregation on the establishment of novel alleles located on multicopy plasmids has already been explored both in models (Santer and Uecker, 2020; Garoña et al, 2023) and experiments (Ilhan et al, 2018; Garoña et al, 2023), showing that segregation reduces the establishment probability of adaptive alleles on multicopy plasmids. Here, we have shown that the effect of segregation is even more drastic in scenarios of heterozygote advantage: the constant loss of heterozygosity to segregation puts strong limits on the probability of successful adaptation unless the fitness of heterozygotes is sufficiently large. The condition that we derived for the required fitness of heterozygotes more generally holds for the maintenance of heterozygosity on multicopy plasmids, even outside a rescue scenario. While we have phrased our model in terms of wild-type and mutant plasmids (i.e. variants of the same plasmid), the results carry over to the establishment and the maintenance of two incompatible plasmids variants sharing the same replication mechanism.

The importance of plasmid-mediated heterozygosity in allowing populations to escape from fitness trade-offs was previously explored by Rodriguez-Beltran et al (2018) in both experiments and models. In a system where different plasmid alleles provide resistance to different antibiotics, they find that fluctuating selection is capable of maintaining heterozygosity, and intermediate antibiotic concentrations maintained heterozygosity the longest (note that unlike in our model, there is always selection for only one plasmid variant at a time). Their model is simpler than ours in one important respect: they bundle all heterozygotes into a single compartment of their ODE model, ignoring the precise number of mutant and wild-type plasmids. To estimate the rate of segregation, Rodriguez-Beltran et al (2018) assume that all heterozygote cells have half-and-half mutant and wild-type composition, so that the probability of having a homozygote daughter cell is 2^1*−n*^. This means that the expected number of heterozygote daughters of a heterozygote cell is 2(1 *−* 2^1*−n*^)(1 + *s*_max_)*/*(2 + *s*_max_), and therefore the branching process founded by a heterozygote cell is supercritical, and the establishment probability of a mutant on a plasmid is positive, if and only if

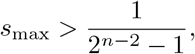

which is a much weaker condition than our Theorem 1 (provided that *n >* 3): ignoring the unbalanced heterozygotes overestimates heterozygote stability. The extension of our model to fluctuating selection remains for future work.

The important role of segregation on multicopy plasmids contrasts strongly with the state of affairs on a haploid bacterial chromosome, where true heterozygosity is impossible, so that our two focal alleles would have to either replace each other completely or be located at different loci and not subject to segregation. Cells in rapidly dividing populations will contain multiple copies of the chromosome and thus be effectively polyploid; although this exposes them to the effects of dominance (Sun et al, 2018) their segregation is not random and heterozygosity cannot be maintained long-term. There is also a strong contrast to the role of segregation in the system one normally thinks of when considering heterozygosity, sexually reproducing diploid organisms. Homologous chromosomes in diploids segregate at meiosis, but heterozygosity is maintained by sexual reproduction; unlike with multicopy plasmids, no selection for heterozygotes is required to maintain heterozygosity at Hardy-Weinberg proportions in the population indefinitely, provided homozygous individuals can thrive and both homozygous types are equally fit. However, if homozygous individuals have fitness less than one as in our rescue scenario and heterozygotes are required for population persistence, there is also a threshold on the heterozygote fitness below which the population cannot survive; unlike for multicopy plasmids, this threshold depends on the fitness of homozygotes. The closest analogue in diploids to the role of segregation in multicopy plasmids is inbreeding, which also reduces heterozygosity in the population without altering allele frequencies. Segregation is in a sense a stronger force in reducing heterozygosity than any degree of partial selfing, since the equilibrium heterozygosity is zero in the absence of selection for the heterozygote (unlike partial selfing, which produces a nonzero equilibrium), but less strong than complete selfing; this can be seen by comparing the *n* = 2 case (when segregation has the largest effect on multicopy plasmids, and also the case most directly comparable to diploids), where 2*/*3 of the daughter cells of heterozygotes are heterozygotes, to a selfing diploid, where only 1*/*2 of the offspring of heterozygotes are heterozygotes. Nevertheless, constraints on heterozygote fitness are stronger for plasmid-mediated rescue of bacterial populations than for rescue of selfing populations: division of a heterozygote cell into two homozygote cells leads to loss of a heterozygote from the population, while offspring reproduction by a selfing heterozygote leaves the heterozygote parent individual intact. This means that the rate of reproduction 1 + *s*_max_ only needs to be greater than two for the selfing individual (or for a random mating population in which homozygotes are lethal or infertile) rather than three as in the bacterial population. The nonplasmid systems that most closely resemble the genetics of multicopy plasmids are the organelle (mitochondrial and plastid) genomes of eukaryotes (Birky, 1983) and the macronuclei of ciliate protists (Allen and Nanney, 1958)—indeed, it was in the latter context that Schensted (1958) developed a model of chromosomal segregation which is equivalent our regular replication model for plasmids and first determined the eigenvalues of the matrix we refer to as *P* in the proof of Theorem 1.

We have also seen that the mechanism or mode of replication has a substantial effect on the fate of a novel mutation in heterozygote advantage scenarios. In the regular replication model, there are no additional stochastic effects in replication, while in the random replication model the rich-get-richer property of the replication process is an additional mechanism reducing heterozygosity in the population. Our understanding of the biological mechanisms of plasmid replication would seem to suggest that random replication is a more realistic model of the replication process of most plasmids; however, some empirical studies have found the regular replication model to be a better fit to the data (Garoña et al, 2023). Here we have followed Novick and Hoppensteadt (1978) in presenting both models to permit a comparison.

We have kept properties of the plasmid such as copy number fixed over the course of rescue. While this may be a realistic assumption over the short timescale of the rescue process, over the longer term (for example in the rescued population) these properties are also subject to evolution. Given the important role of copy number in the fate of bacterial populations depending on plasmid-mediated heterozygosity, the copy number might well be under selection. Some experimental studies have shown rapid evolution of plasmid copy number in response to selection on plasmid-borne traits in other contexts (Dimitriu et al, 2021; San Millan et al, 2015; San Millan et al, 2016). A mutation in the plasmid replication control system that produces a new plasmid type compatible with the existing plasmid—a plasmid speciation event—would also confer a selective advantage if it allowed the mutant and wild-type alleles to exist on plasmids of distinct compatible types, no longer sharing a common copy number between them, and no longer subject to loss of heterozygosity by segregation. Rapid plasmid speciation has also been observed in experiments (Santos-Lopez et al, 2016). Another change in the plasmid that would affect the maintenance of plasmid-mediated heterozygosity, which we have modelled, is the formation of plasmid cointegrates (Hülter et al, 2020; Garoña et al, 2021). Our results showed that stable multimerization of plasmids can increase the probability of successful establishment of the novel allele. Our model of cointegration is deliberately simple and limited, as it is intended as a short demonstration of the importance of cointegration in the heterozygote advantage scenario and not a deep exploration of the cointegration process. Many aspects of the cointegration process not explored here, particularly the precise behaviour of multimers during replication and segregation and the resolution of cointegrates into smaller multimers or monomers, are no doubt important to the fate of alleles on cointegrates and merit further investigation. We especially have not included consequences of multimerization on the stability of plasmid inheritance, which has been examined in a model by Summers et al (1993). The possibility of evolution of a multidrug resistance plasmid from two incompatible single-resistance plasmids has been previously demonstrated by Condit and Levin (1990) both experimentally and in a model; it is left open in their model whether the plasmid carrying both resistance genes has been formed through plasmid fusion or exchange of DNA between plasmids, and they do not account for the plasmid copy number. A detailed model that combines elements of their and our models and allows for dependence of recombination on the plasmid copy number or the ratio of variant frequencies could be interesting to further understand the evolution of multidrug resistance on plasmids.

Broadly speaking, whether rescue occurs or not depends on the availability of rescue mutations and their probability to escape stochastic loss while rare, which is reflected in equation (6) by the product of the mutational input and the establishment probability. In our model, all rescue mutations are assumed to appear after the population decline begins. However, if the mutation were to occur before the environmental change that causes the population decline, then there could already be heterozygotes present in the population at the beginning of the evolutionary rescue process. Their contribution relative to that of *de novo* mutations strongly depends on the fitness effect of the mutation prior to the environmental change and the rate of decay of the wild-type population (Orr and Unckless, 2014). Rescue from the standing genetic variation has been explored in many models of evolutionary rescue (e.g. Orr and Unckless, 2008), including rescue on plasmids (Santer and Uecker, 2020). Plasmids also create avenues for the arrival of rescue mutations through horizontal gene transfer: a mutant plasmid could be introduced into the host population by transformation, or by conjugation (where heterozygosity can only be generated if transfer of the mutant plasmid is not excluded by entry or surface exclusion from the wild-type plasmid variant). While horizontal transfer from a source population to the target population increases the availability of the rescue mutation, horizontal transfer within the target population affects its establishment probability. There has been work on modelling rescue from genes on plasmids arriving and spreading by horizontal transfer, splitting the problem up into the rate of appearance and the establishment probability of the (single-copy) rescue plasmid analogous to our equation (6) (Tazzyman and Bonhoeffer, 2014; Geoffroy and Uecker, 2023).

Plasmids are pervasive in natural populations of bacteria, and play an important role in adaptation and evolution of bacteria. Understanding bacterial evolution therefore involves a good understanding of the population genetics of plasmids. The properties of plasmids as independently replicating units within their hosts make them an intriguing genetic system, with complexities not present for haploid or diploid chromosomes. We have explored one aspect of plasmid genetics—heterozygosity on multicopy plasmids—in the context of evolutionary rescue. We have shown that, as intuitively expected, the maintenance of heterozygosity and thus rescue is impossible below a threshold copy number for a given heterozygote fitness. Going beyond this intuition, our concise criterion in terms of the plasmid copy number and the fitness of heterozygous cells quantifies the conditions for the persistence of plasmid-mediated heterozygosity through heterozygote advantage, and thus contributes to a population genetic theory of bacterial evolution.

## Supporting information

Numerical and simulation code

## Acknowledgments

The authors thank Félix Geoffroy and Mario Santer for helpful discussions, and Reinhard Bürger and two anonymous reviewers for their valuable comments on the manuscript.

## Statements and Declarations

### Availability of data and materials

No new data was generated in the course of this theoretical work.

### Code availability

The programs used to numerically solve the models and produce the figures and the simulation code are available as supplementary material.

### Authors’ contributions

ID: Conceptualization, formal analysis, methodology, software, writing—original draft; HU: Conceptualization, methodology, supervision, writing—review & editing.

## Appendix A Mathematical notes

In this appendix, we derive several combinatorial results needed in the main text of the paper.

First, for completeness, we derive the distribution of outcomes of a Pólya urn process (Eggenberger and Pólya, 1923). Suppose we start with an urn of *n* balls, *i* blue and *n − i* white. We pick a ball out of the urn, then replace it together with another ball of the same colour. After repeating this process *m − n* times, there are *m* balls in the urn: what is the probability that *k* of them are blue?

### Proposition 1.

*Denote the probability described above by* 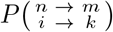. *Then for m ≥ n > 0 and k ≥ i ≥ 0*,

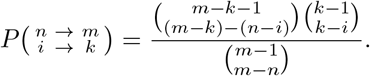

*Proof*. First we prove the special case

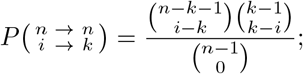

since no balls are drawn in this case, the left-hand side is 1 if *i* = *k* and 0 otherwise. The lower indices of the binomial coefficients in the numerator are negatives of each other, so the coefficients can only both be nonzero if *k − i* = 0, and in that case all three coefficients are equal to 1, satisfying the equality.

Next we prove another special case

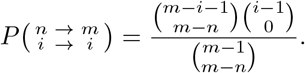

Suppose that this equality is established for 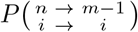. In order to reach *m* balls in the urn without increasing the number of blue balls, we must first reach *m −* 1 balls, then draw a white ball on the final draw. Therefore

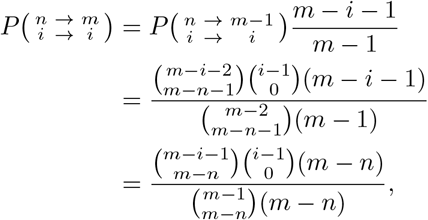

and by induction on *m* (with our previous special case as the base case), the equality is established.

Finally, we prove the general case. Suppose we know the equality holds for 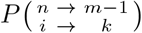 and 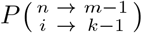. In order for there to be *k* blue balls in the urn when we get to the point of having *m* total balls, then either the last ball drawn was white, and there were *k* blue out of *m −* 1 total balls before, or the last ball drawn was blue, and there were *k −* 1 blue out of *m −* 1 total balls before. Thus

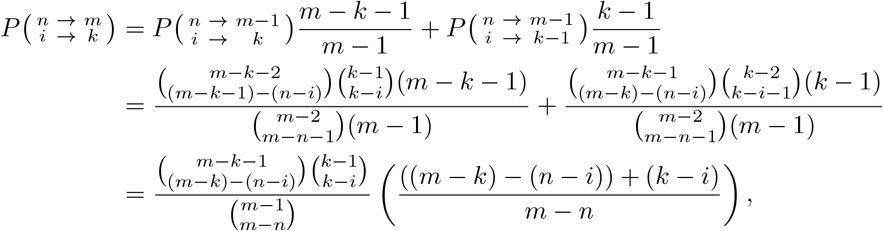

and by induction on *m* and *k*, the equality is established.

Obviously if we replace blue and white balls in the urn with mutant and wild-type plasmids in a cell, this process describes the replication of plasmids under the random replication model. The probability that starting with a cell with *i* mutant plasmids there are *j* + *k* mutant plasmids after replication is then 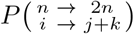.

When a mutation occurs during plasmid replication in the random replication model, it is possible for the mutant plasmid to be replicated again the same generation, and so the initial mutant cell can end up with multiple mutant plasmids. Suppose that a mutation occurs during each replication of a plasmid with probability *u*; then the probability that in a certain generation exactly one mutation occurs, it occurs during the (*ℓ* + 1)-th replication, and the cell ends up with *i* mutant plasmids after all of the replication has occurred but before cell division is

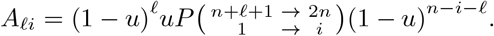

We assume that *u* is small, so that 1 *− u ≈* 1; then the probability that a mutation occurs in a given generation is

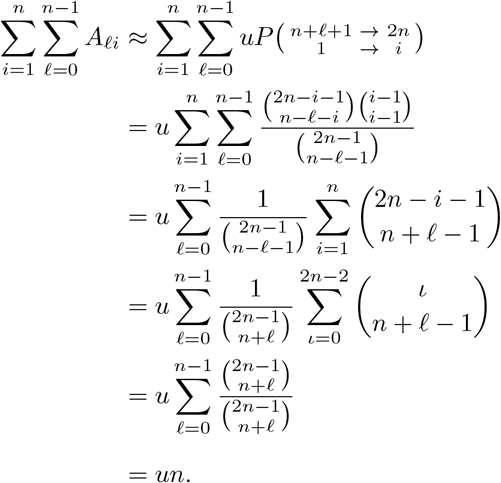

The probability of ending up with *i* mutant plasmids conditional on a mutation occurring is then

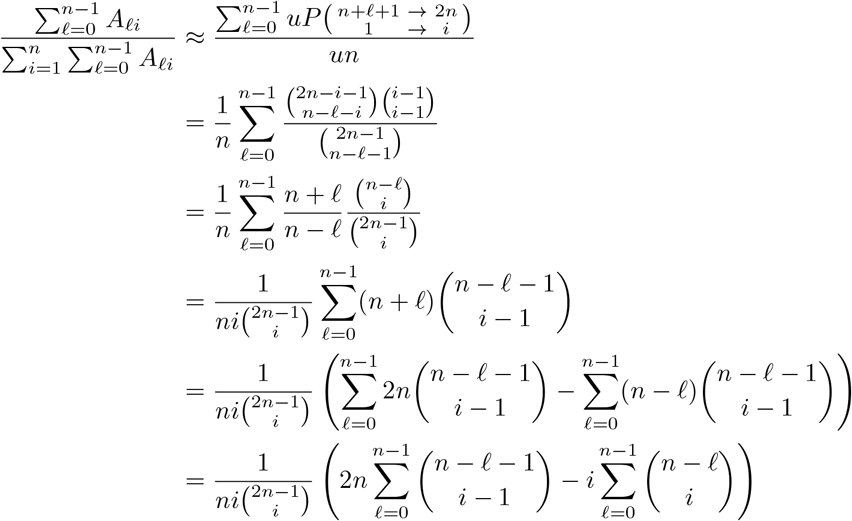

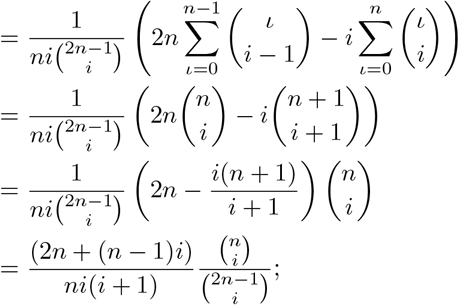

by substituting *i* = *j* + *k*, we obtain the replication part of equation (5).

### Lemma 1.

*For all integers n >* 0, 0 *≤ β ≤ n, and γ >* 0,

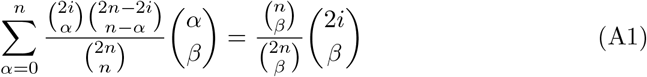

*and*

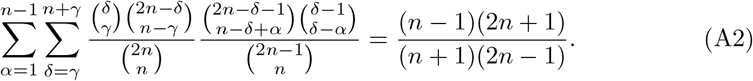

*Proof*. For (A1), we have that

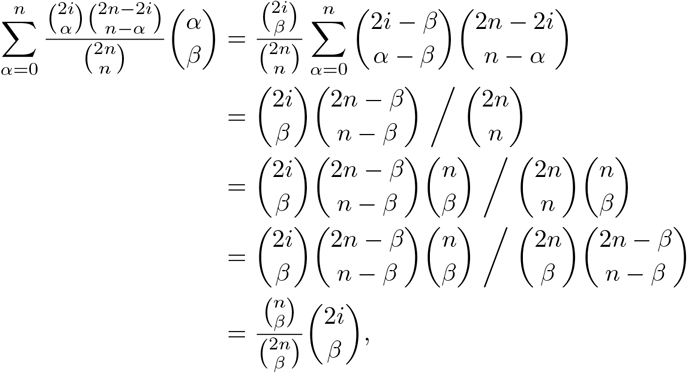

where the first and fourth equalities use trinomial revision and the third uses Vandermonde convolution (with an implicit change of index to *α − β*). For (A2), we have that

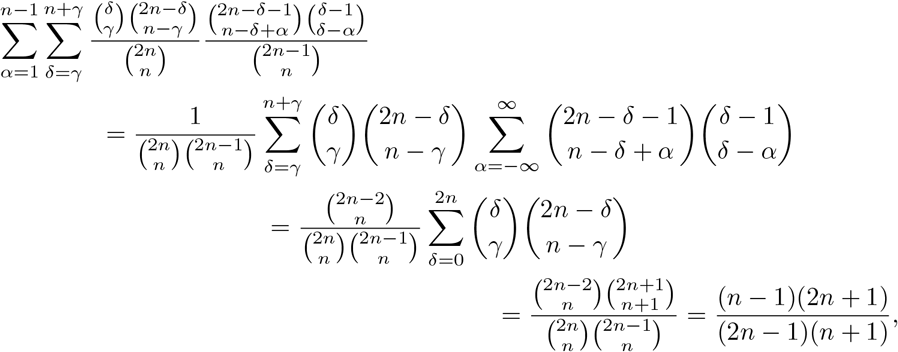

where the second equality uses Vandermonde convolution (with an implicit change of index to *δ − α*) and the third uses equation (5.26) of Graham et al (1994).

## References

Alexander HK, Bonhoeffer S (2012) Pre-existence and emergence of drug resistance in a generalized model of intra-host viral dynamics. Epidemics 4(4):187–202. 10.1016/j.epidem.2012.10.001

Allen SL, Nanney DL (1958) An analysis of nuclear differentiation in the selfers of tetrahymena. The American Naturalist 92(864):139–160. 10.1086/282022

Anda M, Ohtsubo Y, Okubo T, et al (2015) Bacterial clade with the ribosomal RNA operon on a small plasmid rather than the chromosome. Proceedings of the National Academy of Sciences 112(46):14343–14347. 10.1073/pnas.1514326112

Batra A, Roemhild R, Rousseau E, et al (2021) High potency of sequential therapy with only β-lactam antibiotics. eLife 10:e68876. 10.7554/elife.68876

Beijersbergen A, Smith SJ, Hooykaas PJ (1994) Localization and topology of VirB proteins of Agrobacterium tumefaciens. Plasmid 32(2):212–218. 10.1006/plas.1994.1057

Birky CW Jr. (1983) Relaxed cellular controls and organelle heredity. Science 222(4623):468–475. 10.1126/science.6353578

Brovedan M, Repizo GD, Marchiaro P, et al (2019) Characterization of the diverse plasmid pool harbored by the bla_NDM-1_-containing Acinetobacter bereziniae HPC229 clinical strain. PLOS ONE 14(11):e0220584. 10.1371/journal.pone.0220584

Burian J, Guller L, Mavčor M, et al (1997) Small cryptic plasmids of multiplasmid, clinical Escherichia coli. Plasmid 37(1):2–14. 10.1006/plas.1996.1273

Carattoli A (2013) Plasmids and the spread of resistance. International Journal of Medical Microbiology 303:298–304. 10.1016/j.ijmm.2013.02.001

Casjens S, Palmer N, van Vugt R, et al (2000) A bacterial genome in flux: the twelve linear and nine circular extrachromosomal DNAs in an infectious isolate of the Lyme disease spirochete Borrelia burgdorferi. Molecular Microbiology 35(3):490–516

Chiang CS, Bremer H (1991) Maintenance of pBR322-derived plasmids without functional RNAI. Plasmid 26(3):186–200. 10.1016/0147-619x(91)90042-u

Coluzzi C, Garcillán-Barcia MP, de la Cruz F, et al (2022) Evolution of plasmid mobility: Origin and fate of conjugative and nonconjugative plasmids. Molecular Biology and Evolution 39(6):msac115. 10.1093/molbev/msac115

Condit R, Levin BR (1990) The evolution of plasmids carrying multiple resistance genes: The role of segregation, transposition, and homologous recombination. The American Naturalist 135(4):573–596. 10.1086/285063

Cullum J, Broda P (1979) Rate of segregation due to plasmid incompatibility. Genetical Research 33(1):61–79. 10.1017/s0016672300018176

Dimitriu T, Matthews AC, Buckling A (2021) Increased copy number couples the evolution of plasmid horizontal transmission and plasmid-encoded antibiotic resistance. Proceedings of the National Academy of Sciences 118(31). 10.1073/pnas.2107818118

Eggenberger F, Pólya G (1923) Über die statistik verketteter vorgänge. Zeitschrift für Angewandte Mathematik und Mechanik 3(4):279–289. 10.1002/zamm.19230030407

Falkow S (1975) Infectious Multiple Drug Resistance.

Pion Farr AD, Pesce D, Das SG, et al (2023) The fitness of beta-lactamase mutants depends nonlinearly on resistance level at sublethal antibiotic concentrations. mBio 14(3). 10.1128/mbio.00098-23

Frobenius G (1912) Ü ber matrizen aus nicht negativen elementen. Sitzungsberichte der Königlich Preussischen Akademie der Wissenschaften pp 456–477

Gama JA, Zilhão R, Dionisio F (2018) Impact of plasmid interactions with the chromosome and other plasmids on the spread of antibiotic resistance. Plasmid 99:82–88. 10.1016/j.plasmid.2018.09.009

Garcillán-Barcia MP, Alvarado A, de la Cruz F (2011) Identification of bacterial plasmids based on mobility and plasmid population biology. FEMS Microbiology Reviews 35(5):936–956. 10.1111/j.1574-6976.2011.00291.x

Garoña A, Hülter NF, Picazo DR, et al (2021) Segregational drift constrains the evolutionary rate of prokaryotic plasmids. Molecular Biology and Evolution 10.1093/molbev/msab283

Garoña A, Santer M, Hülter NF, et al (2023) Segregational drift hinders the evolution of antibiotic resistance on polyploid replicons. PLOS Genetics 19(8):e1010829. 10.1371/journal.pgen.1010829

Geoffroy F, Uecker H (2023) Limits to evolutionary rescue by conjugative plasmids. Theoretical Population Biology 154:102–117. 10.1016/j.tpb.2023.10.001

Gillespie DT (1976) A general method for numerically simulating the stochastic time evolution of coupled chemical reactions. Journal of Computational Physics 22(4):403–434. 10.1016/0021-9991(76)90041-3

Gomulkiewicz R, Holt RD (1995) When does evolution by natural selection prevent extinction? Evolution 49(1):201–207. 10.1111/j.1558-5646.1995.tb05971.x

Graham RL, Knuth DE, Patashnik O (1994) Concrete Mathematics, 2nd edn. Addison-Wesley

Gustafsson P, Nordström K (1980) Control of plasmid R1 replication: Kinetics of replication in shifts between different copy number levels. Journal of Bacteriology 141(1):106–110. 10.1128/jb.141.1.106-110.1980

Haldane JBS (1927) A mathematical theory of natural and artificial selection, part v: Selection and mutation. Mathematical Proceedings of the Cambridge Philosophical Society 23(7):838–844. 10.1017/s0305004100015644

Halleran AD, Flores-Bautista E, Murray RM (2019) Quantitative characterization of random partitioning in the evolution of plasmid-encoded traits. BioRxiv preprint, 10.1101/594879

Hernandez-Beltran JCR, Miró Pina V, Siri-Jégousse A, et al (2022) Segregational instability of multicopy plasmids: A population genetics approach. Ecology and Evolution 12(12). 10.1002/ece3.9469

Hülter NF, Wein T, Effe J, et al (2020) Intracellular competitions reveal determinants of plasmid evolutionary success. Frontiers in Microbiology 11. 10.3389/fmicb.2020.02062

Ilhan J, Kupczok A, Woehle C, et al (2018) Segregational drift and the interplay between plasmid copy number and evolvability. Molecular Biology and Evolution 36(3):472–486. 10.1093/molbev/msy225

Ishii K, Hashimoto-Gotoh T, Matsubara K (1978) Random replication and random assortment model for plasmid incompatibility in bacteria. Plasmid 1(4):435–445. 10.1016/0147-619x(78)90002-1

Ismail E, Blom J, Bultreys A, et al (2014) A novel plasmid pEA68 of Erwinia amylovora and the description of a new family of plasmids. Archives of Microbiology 196(12):891–899. 10.1007/s00203-014-1028-5

Kiyosawa H, Hughes JE, Podgorski GJ, et al (1993) Small circular plasmids of the eukaryote Dictyostelium purpureum define two novel plasmid families. Plasmid 30(2):106–118. 10.1006/plas.1993.1038

Martin G, Aguilée R, Ramsayer J, et al (2013) The probability of evolutionary rescue: towards a quantitative comparison between theory and evolution experiments. Philosophical Transactions of the Royal Society B 368(1610):20120088. 10.1098/rstb.2012.0088

Mode CJ (1971) Multitype Branching Processes. Modern Analytic and Computational Methods in Science and Mathematics, American Elsevir Publishing Company, Inc.

Nordström K (2006) Plasmid r1—replication and its control. Plasmid 55(1):1–26. 10.1016/j.plasmid.2005.07.002

Novick RP (1987) Plasmid incompatibility. Microbiological Reviews 51(4):381–395. 10.1128/mr.51.4.381-395.1987

Novick RP, Hoppensteadt F (1978) On plasmid incompatibility. Plasmid 1(4):421–434. 10.1016/0147-619x(78)90001-x

Orr HA, Unckless RL (2008) Population extinction and the genetics of adaptation. The American Naturalist 172(2):160–169. 10.1086/589460

Orr HA, Unckless RL (2014) The population genetics of evolutionary rescue. PLoS Genetics 10(8):e1004551. 10.1371/journal.pgen.1004551

Portnoy DA, Martinez RJ (1985) Role of a plasmid in the pathogenicity of Yersinia species. In: Goebel W (ed) Genetic Approaches to Microbial Pathogenicity. No. 118 in Current Topics in Microbiology and Immunology, Springer-Verlag, p 29–51, 10.1007/978-3-642-70586-13

Prangishvili D, Albers SV, Holz I, et al (1998) Conjugation in archaea: Frequent occurrence of conjugative plasmids in Sulfolobus. Plasmid 40(3):190–202. 10.1006/plas.1998.1363

Projan SJ, Monod M, Narayanan CS, et al (1987) Replication properties of pIM13, a naturally occurring plasmid found in Bacillus subtilis, and of its close relative pE5, a plasmid native to Staphylococcus aureus. Journal of Bacteriology 169(11):5131–5139. 10.1128/jb.169.11.5131-5139.1987

Rodriguez-Beltran J, Hernandez-Beltran JCR, DelaFuente J, et al (2018) Multicopy plasmids allow bacteria to escape from fitness trade-offs during evolutionary innovation. Nature Ecology & Evolution 2(5):873–881. 10.1038/s41559-018-0529-z

Rodríguez-Beltrán J, DelaFuente J, León-Sampedro R, et al (2021) Beyond horizontal gene transfer: the role of plasmids in bacterial evolution. Nature Reviews Microbiology 19(6):347–359. 10.1038/s41579-020-00497-1

San Millan A, Escudero JA, Gutierrez B, et al (2009) Multiresistance in Pasteurella multocida is mediated by coexistence of small plasmids. Antimicrobial Agents and Chemotherapy 53(8):3399–3404. 10.1128/aac.01522-08

San Millan A, Santos-Lopez A, Ortega-Huedo R, et al (2015) Small-plasmid-mediated antibiotic resistance is enhanced by increases in plasmid copy number and bacterial fitness. Antimicrobial Agents and Chemotherapy 59(6):3335–3341. 10.1128/aac.00235-15

San Millan A, Escudero JA, Gifford DR, et al (2016) Multicopy plasmids potentiate the evolution of antibiotic resistance in bacteria. Nature Ecology & Evolution 1(1). 10.1038/s41559-016-0010

Santer M, Uecker H (2020) Evolutionary rescue and drug resistance on multicopy plasmids. Genetics 215(3):847–868. 10.1534/genetics.119.303012

Santer M, Kupczok A, Dagan T, et al (2022) Fixation dynamics of beneficial alleles in prokaryotic polyploid chromosomes and plasmids. Genetics 222(2). 10.1093/genetics/iyac121

Santos-Lopez A, Bernabe-Balas C, Ares-Arroyo M, et al (2016) A naturally occurring single nucleotide polymorphism in a multicopy plasmid produces a reversible increase in antibiotic resistance. Antimicrobial Agents and Chemotherapy 61(2). 10.1128/aac.01735-16

Schensted IV (1958) Model of subnuclear segregation in the macronucleus of ciliates. The American Naturalist 92(864):161–170. 10.1086/282023

Sewastjanow BA (1975) Verzweigungsprozesse. R. Oldenbourg Verlag

Silver S (1992) Plasmid-determined metal resistance mechanisms: Range and overview. Plasmid 27(1):1–3. 10.1016/0147-619x(92)90001-q

Smillie C, Garcillán-Barcia MP, Francia MV, et al (2010) Mobility of plasmids. Microbiology and Molecular Biology Reviews 74(3):434–452. 10.1128/mmbr.00020-10

Smith BA, Dougherty K, Clark M, et al (2021) Experimental evolution of the megaplasmid pMPPla107 in Pseudomonas stutzeri enables identification of genes contributing to sensitivity to an inhibitory agent. Philosophical Transactions of the Royal Society B 377(1842):20200474. 10.1098/rstb.2020.0474

Strang G (2009) Introduction to Linear Algebra, 4th edn. Wellesley-Cambridge Press

Summers DK, Sherratt DJ (1984) Multimerization of high copy number plasmids causes instability: ColE1 encodes a determinant essential for plasmid monomerization and stability. Cell 36(4):1097–1103. 10.1016/0092-8674(84)90060-6

Summers DK, Beton CWH, Withers HL (1993) Multicopy plasmid instability: the dimer catastrophe hypothesis. Molecular Microbiology 8(6):1031–1038. 10.1111/j.1365-2958.1993.tb01648.x

Sun L, Alexander HK, Bogos B, et al (2018) Effective polyploidy causes phenotypic delay and influences bacterial evolvability. PLOS Biology 16(2):e2004644. 10.1371/journal.pbio.2004644

Tazzyman SJ, Bonhoeffer S (2014) Plasmids and evolutionary rescue by drug resistance. Evolution 68(7):2066–2078. 10.1111/evo.12423

Uhlin BE, Nordström K (1975) Plasmid incompatibility and control of replication: copy mutants of the R-factor R1 in Escherichia coli K-12. Journal of Bacteriology 124(2):641–649. 10.1128/jb.124.2.641-649.1975

Wardell GE, Hynes MF, Young PJ, et al (2021) Why are rhizobial symbiosis genes mobile? Philosophical Transactions of the Royal Society B 377(1842):20200471. 10.1098/rstb.2020.0471

Yu W, Gillies K, Kondo JK, et al (1996) Loss of plasmid-mediated oligopeptide transport system in lactococci: Another reason for slow milk coagulation. Plasmid 35(3):145–155. 10.1006/plas.1996.0017

Zielenkiewicz U, Cegłowski P (2001) Mechanisms of plasmid stable maintenance with special focus on plasmid addiction systems. Acta Biochimica Polonica 48(4):1003– 1023. 10.18388/abp.20013863

