## Supplementary figures for "Evolutionary rescue of bacterial populations by heterozygosity on multicopy plasmids"

### Supplementary Material to "Evolutionary rescue of bacterial populations by heterozygosity on multicopy plasmids"

Stochastic simulations were performed to verify the results of the numerical analysis. The branching process was simulated in discrete time, with the number and type of daughters of each cell in every generation being sampled from the same progeny distribution as in the analytic model, except that the assumptions on mutation were relaxed. Mutations were allowed to occur at every plasmid replication in every cell division, as opposed to the analytic model where we assumed that there is at most one mutation per host generation, and never multiple mutation events in the same lineage. The simulation programs are available as a further supplement.

Establishment probability was calculated by simulating populations starting from a pair of cells sampled from the distribution of pairs of daughters of a homozygote wild-type cell conditional on a mutation occurring; the population was considered to have gone extinct if the number of cells reached zero, and the mutation to have established if the number of heterozygote cells reached 100000. The use of this threshold was validated after the estimated establishment probability was calculated by verifying that the probability that at least one of the simulations that resulted in an establishment was a false positive (given the calculated establishment probability) was less than 0.1 %. The calculated establishment probabilities are shown in Figure S1; the 95 % confidence intervals for the estimated probabilities are narrow enough that no error bars are shown in the figure, as they would be obscured by the symbols. The difference between the simulations and the numerical approximations is shown in Figure S2; in general, the numerical results are very close to the simulation results, within the natural stochastic variation of the simulation results.

Rescue probability was calculated by simulating populations starting from a homozygote wild-type population of a given initial size $N_0 = 100000$. The population was considered to have gone extinct if the number of cells reached zero, and to have been rescued if the number of heterozygote cells reached $N_0$. The calculated rescue probabilities are shown in Figure S3; the 95 % confidence intervals for the estimated probabilities are narrow enough that no error bars are shown in the figure, as they would be obscured by the symbols. The difference between the simulations and the numerical approximations is shown in Figure S4; again, the differences are within the stochastic variability of the simulations.

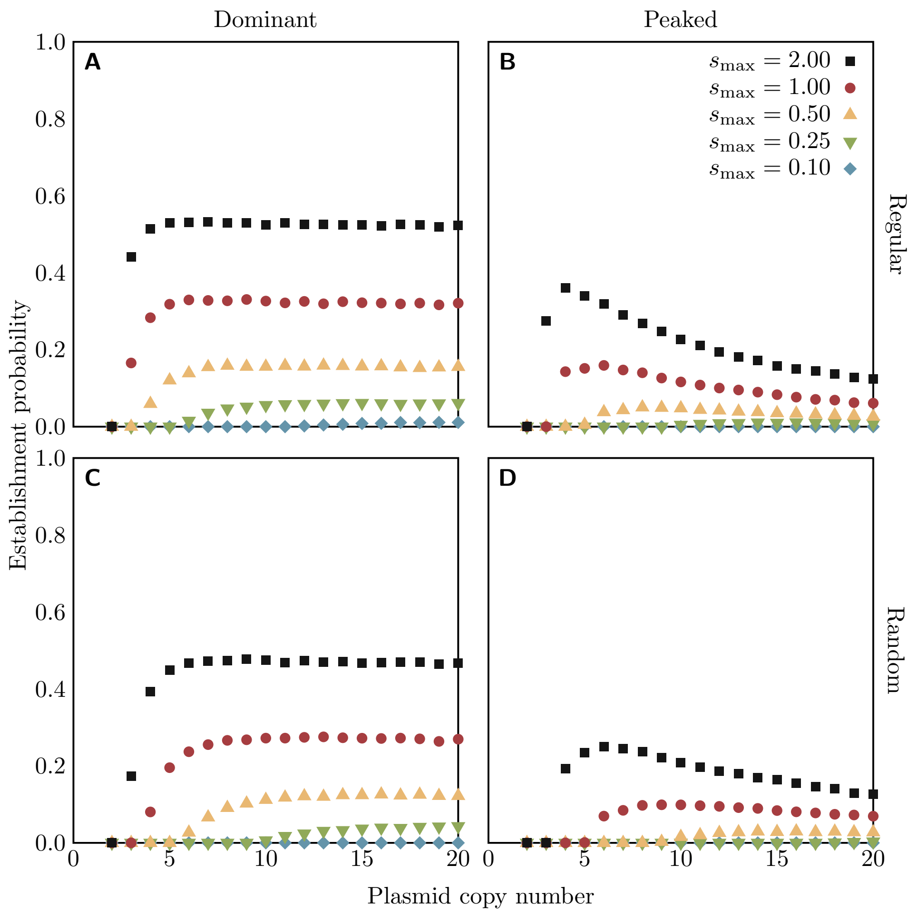


Figure S1: Establishment probabilities of rescue mutations on a plasmid of a given copy number in the heterozygote advantage scenario, under the regular (top row) or random (bottom row) replication assumption, calculated from stochastic simulations. Colours of points indicate the fitness $s_{\text{max}}$ of all heterozygotes (with the dominant fitness function, left column) or the maximum fitness of heterozygotes (with the peaked fitness function, right column). For all cases, $s_{\text{min}} = -0.1$ and $u = 10^{-6}$.

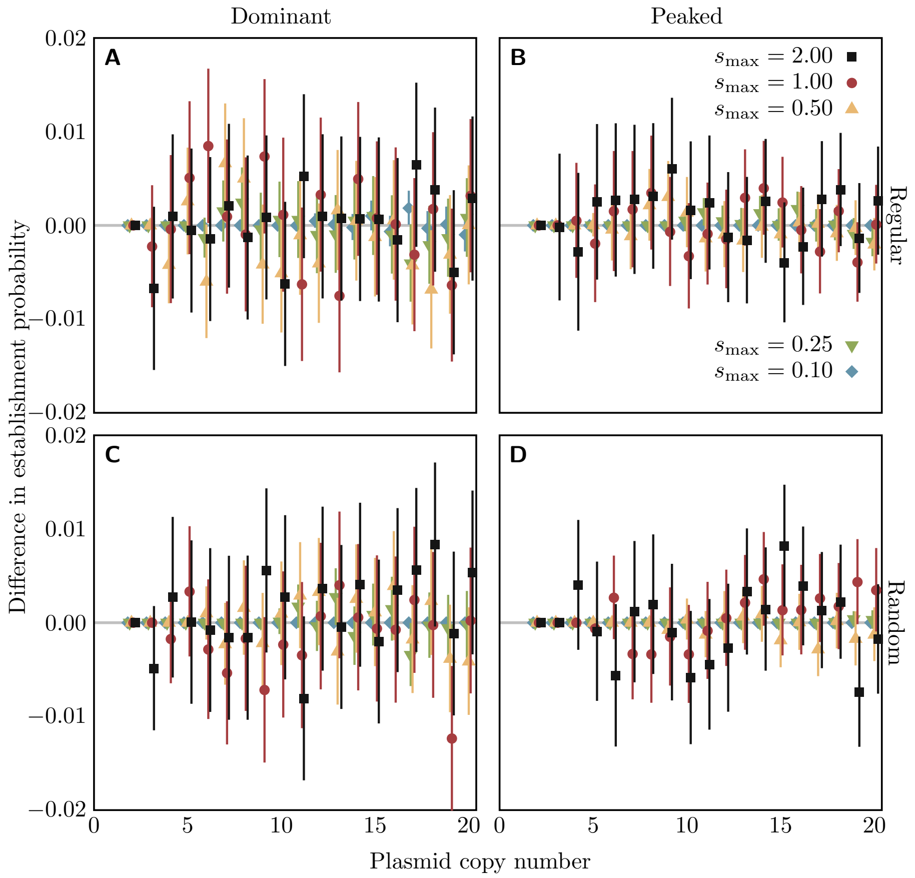


Figure S2: Difference between the stochastic simulation and numerically derived establishment probabilities, with 95 % confidence intervals from the likelihood-ratio test. Simulation parameters described in the caption to Figure S1.

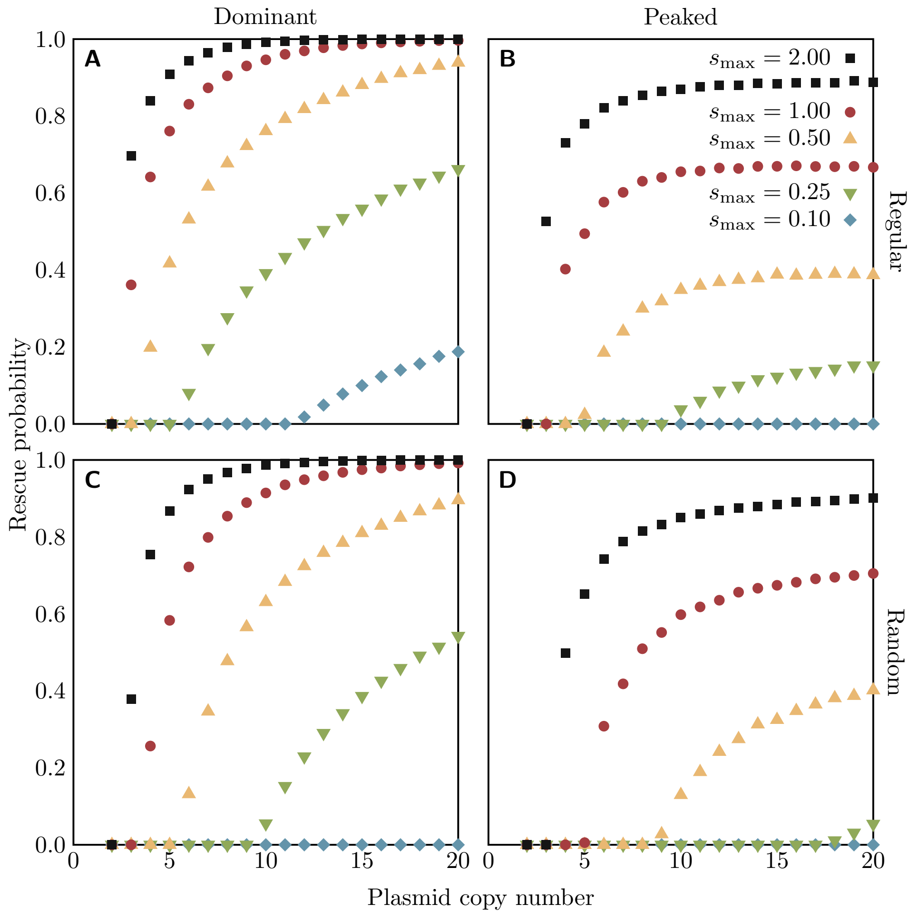


Figure S3: Probabilities of a bacterial population being rescued by a mutation on a plasmid of a given copy number in the heterozygote advantage scenario, under the regular (top row) or random (bottom row) replication assumption, calculated from stochastic simulations. Colours of points indicate the fitness $s_{\max}$ of all heterozygotes (with the dominant fitness function, left column) or the maximum fitness of heterozygotes (with the peaked fitness function, right column). For all cases, $s_{\min} = -0.1$, $u = 10^{-6}$, and $N_0 = 10^5$.

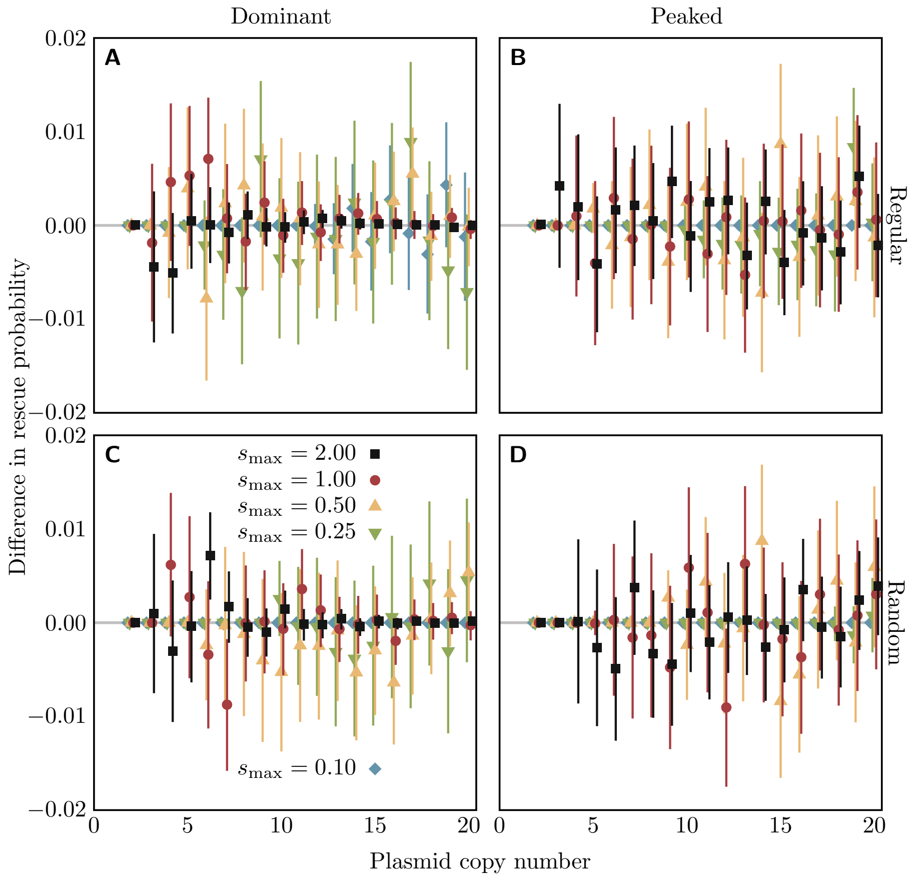


Figure S4: Difference between the stochastic simulation and numerically derived rescue probabilities, with 95 % confidence intervals from the likelihood-ratio test. Simulation parameters described in the caption to Figure S3.
