## Supplementary material for "Evolutionary rescue of bacterial populations by heterozygosity on multicopy plasmids": Numerical and simulation code: Cointegrates.hs

```
{-# LANGUAGE RecordWildCards #-}
-- | The implementation of the model with cointegration.
module Cointegrates (
    CointegrateModel,
    cointegrateModel,
    CopyNumber,
    copyNumber
) where

import Prelude hiding ((<>))

import Control.Arrow ((&&&))
import Data.Bifunctor (first)
import Data.List (findIndex, foldl', tails)
import Data.Maybe (fromJust)
import Numeric (expm1)
import Numeric.LinearAlgebra (Matrix, Vector, assoc, diag, fromList, size, tr,
    (!), (<>), (#>))

import EvolutionaryRescue

-- | A parameterized instance of the model with cointegration.
data CointegrateModel = CointegrateModel {
    -- | The index (into the various matrices) corresponding to the type of the
    -- initial cell of the process.
    start :: Int,
    -- | The reproduction function \(P(a \to \{b,c\})\) in list form: the
    -- @i@th entry in this list is a list of tuples @(p, (j,k))@, where @p@ is
    -- the probability of obtaining daughters with cell types @j@ and @k@ from a
    -- cell with cell type @i@.
    reprList :: [[(Double, (Int, Int))]],
    -- | The matrix the @(i,j)@ element of which is the probability of getting a
    -- cell of type @j@ from a cell of type @i@ by cointegration and resolution.
    cointResolvMat :: Matrix Double,
    -- | The reproduction probability for each cell type. Note that variables
    -- with names starting with ρ are also used for the resolution probabilities
    -- below: be careful not to confuse them.
    ρ :: Vector Double,
    -- | The quantity \(unN_0(1+s_\mathrm{min})/s_\mathrm{min}\), needed to
    -- calculate the rescue probability.
    rescCoeff :: Double,
    -- | The quantity \(1/s_\mathrm{min}\), needed to calculate the mean time
    -- to emergence.
    etCoeff :: Double
}

instance RescueModel CointegrateModel where
    typeCount = size . ρ
    fdf (CointegrateModel{..}) q = (objective, jacobian)
      where objective = 1 - ρ + ρ*(cointResolvMat #> fromList r)
            r = fmap (foldl' (\acc (p, (a, b)) -> acc + p * (q!a) * (q!b)) 0)
                reprList
            jacobian = diag ρ <> cointResolvMat <> core
            core = assoc (size cointResolvMat) 0 (do
                (parent, children) <- zip [0..] reprList
                (prob, (child1, child2)) <- children
                if child1 == child2
                    then [((parent, child1), 2*prob*(q!child2))]
                    else [((parent, child1), prob*(q!child2)),
                        ((parent, child2), prob*(q!child1))])
    expectedOffspring (CointegrateModel{..}) =
        diag ρ <> tr cointResolvMat <> assoc (size cointResolvMat) 0 (do
            (parent, children) <- zip [0..] reprList
            (prob, (child1, child2)) <- children
            if child1 == child2
                then [((parent, child1), 2*prob)]
                else [((parent, child1), prob), ((parent, child2), prob)])
    mutantExtinction = Left . uncurry e . (start &&& typeCount)
    rescueProb cm pest = - expm1 (pest * rescCoeff cm)
    meanEmergenceTime rm pest = etCoeff rm * exp (pest * rescCoeff rm) /
        expm1 (pest * rescCoeff rm) * expIntSeries (-pest * rescCoeff rm)

-- | Create a new instance of the 'CointegrateModel' with the given parameters.
-- @cointegrateModel smax smin uN κf ρf ('copyNumber' n)@ is a model with
-- 
-- * regular replication,
-- * the possibility of plasmids cointegrating and cointegrates resolving
--   (cointegration occurs at most once per generation, with probability @κf p@,
--   where @p@ is the number of plasmids in the cell, and resolution occus at
--   most once per generation with probability @ρf m@, where @m@ is the number
--   of multimers in the cell),
-- * a dominant fitness function with maximum value @smax@ and minimum value
--   @smin@,
-- * copy number @n@, and
-- * product \(uN_0\) of mutation rate and initial wild-type population size of
--   @uN@.
cointegrateModel :: Double -> Double -> Double -> (Int -> Double) ->
    (Int -> Double) -> CopyNumber -> CointegrateModel
cointegrateModel smax smin uN κf ρf (CN n cts rs cmat rmat) =
    CointegrateModel{..}
  where start = fromJust (findIndex (startingType n) cts)
        reprList = rs
        κvec = fromList (fmap (κf . plasmidCount) cts)
        cointMat = diag (1 - κvec) + diag κvec <> cmat
        ρvec = fromList (fmap (ρf . multimerCount) cts)
        resolvMat = diag (1 - ρvec) + diag ρvec <> rmat
        cointResolvMat = resolvMat <> cointMat
        s = fromList (fmap (\ct -> if doomed ct then smin else smax) cts)
        ρ = (1 + s) / (2 + s)
        rescCoeff = uN * (1 + smin) * fromIntegral n / smin
        etCoeff = recip smin

-- | The copy number dependent data, intended to be cached for use in multiple
-- calls to 'cointegrateModel'.
data CopyNumber = CN Int -- ^ The copy number.
    [CellType] -- ^ The list of possible cell types at this copy number.
    [[(Double, (Int, Int))]] -- ^ Reproduction probabilities.
    (Matrix Double) -- ^ Cointegration probability matrix.
    (Matrix Double) -- ^ Resolution probability matrix.

-- | Calculate all of the copy number specific data for the given copy number.
copyNumber :: Int -> CopyNumber
copyNumber n = CN n cts (reproduction cts) cmat rmat
  where cts = celltypes n
        cmat = assoc (length cts,length cts) 0 (concat
            (zipWith (\r -> fmap (\(p, c) -> ((r,c),p)))
            [0..] (cointegration cts)))
        rmat = assoc (length cts,length cts) 0 (concat
            (zipWith (\r -> fmap (\(p, c) -> ((r,c),p)))
            [0..] (resolution cts)))

--------------------------------------------------------------------------------

-- Reproduction

-- | Calculate the reproduction probabilities from the given list of cell types.
-- The @i@th element of the returned list is a list of @(p,(j,k))@ pairs, where
-- @i@, @j@, and @k@ represent cell types as indices into the given list, and
-- @p@ is the probability of producing @j@- and @k@-type daughters from an
-- @i@-type cell.
reproduction :: [CellType] -> [[(Double, (Int, Int))]]
reproduction cts = fmap (\parent ->
    let res = filter ((0 /=) . fst) (fmap (first (parent →)) children) in
        let denom = sum (fmap fst res) in
            fmap (\(count, to) -> (fromIntegral count / fromIntegral denom, to)) res) cts
  where children = do
            (ct1,idx1):rest <- tails (zip cts [0..])
            (ct2,idx2) <- (ct1,idx1):rest
            return ((ct1,ct2), (idx1,idx2))

-- | Return the number of ways to produce the given pair of daughter cells from
-- the given parent cell. The mechanism of the calculation is to iterate through
-- all of the plasmid types, comparing the numbers of each type in the three
-- input cell types.
(→) :: CellType -> (CellType, CellType) -> Int
m → (d1, d2) = extract (foldl' reprodFoldStep (IdenticalSoFar 1)
    (zipWith3 joinCellTypes m d1 d2))
  where extract (IdenticalSoFar res) = res
        extract (Differing res) = 2*res
        extract Unpossible = 0
        joinCellTypes (_, pn) (_, n1) (_, n2) = (pn,n1,n2)

-- | This type is used to track the state while iterating through plasmid types
-- and comparing parent and daughter cells to determine the reproduction
-- probabilities. The state is either 'IdenticalSoFar' if the two daughter cell
-- types have been identical so far, 'Differing' if they have differed, and
-- 'Unpossible' if it has been determined that these daughter cell types cannot
-- come from the given parent cell types. In the former two cases, the argument
-- is the number of ways to segregate the plasmids of the types considered so
-- far.
data ReprodFoldState = IdenticalSoFar Int
                     | Differing Int
                     | Unpossible

-- | A single step of the iteration through all plasmid types. The second
-- argument gives the number of plasmids of the current type in the parent cell,
-- the first daughter, and the second daughter respectively.
reprodFoldStep :: ReprodFoldState -> (Int, Int, Int) -> ReprodFoldState
reprodFoldStep (IdenticalSoFar res) (pn, n1, n2) = case (2*pn == n1 + n2, n1 == n2) of
    (True, True) -> IdenticalSoFar (res * ((2*pn) `choose` n1))
    (True, False) -> Differing (res * ((2*pn) `choose` n1))
    (False, _) -> Unpossible
reprodFoldStep (Differing res) (pn, n1, n2) = if 2*pn == n1 + n2
    then Differing (res * ((2*pn) `choose` n1))
    else Unpossible
reprodFoldStep Unpossible _ = Unpossible

-- Cointegration and Resolution

-- | From a list of cell types, return the distribution of possible outcomes of
-- a single cointegration event. The @i@th element of the returned list is a
-- list of @(p,j)@ pairs, where @i@ and @j@ represent cell types as indices into
-- the given list, and @p@ is the probability of producing a @j@-type cell from
-- an @i@-type cell by a single cointegration event.
cointegration :: [CellType] -> [[(Double, Int)]]
cointegration cts = fmap (\initial ->
    let res = filter ((0 /=) . fst) (zip (fmap (initial ⇒) cts) [0..]) in
        let denom = sum (fmap fst res) in
            fmap (\(count, to) -> (fromIntegral count / fromIntegral denom, to)) res) cts

-- | From a list of cell types, return the distribution of possible outcomes of
-- a single resolution event. The @i@th element of the returned list is a list
-- of @(p,j)@ pairs, where @i@ and @j@ represent cell types as indices into the
-- given list, and @p@ is the probability of producing a @j@-type cell from an
-- @i@-type cell by a single resolution event.
resolution :: [CellType] -> [[(Double, Int)]]
resolution cts = fmap (\initial ->
    let res = filter ((0 /=) . fst) (zip (fmap (⇐ initial) cts) [0..]) in
        let denom = sum (fmap fst res) in
            fmap (\(count, to) -> (fromIntegral count / fromIntegral denom, to)) res) cts

-- | Return the number of ways that the second cell type can be produced from
-- the first by a single cointegration event. This is calculated by iterating
-- through all of the plasmid types and comparing the number of plasmids of that
-- type in the initial and final cell type respectively to check that the
-- transition is possible and count the number of pairs of plasmids of the
-- initial type.
(⇒) :: CellType -> CellType -> Int    
(⇒) = fmap (extract . foldl' cointFoldStep NothingSoFar) . zipWith joinCellTypes
  where extract (Done res) = res
        extract _ = 0
        joinCellTypes (pt, n1) (_, n2) = (pt, n1, n2)

-- | Return the number of ways that the second cell type can be produced from
-- the first by a single resolution event. This is calculated by iterating
-- through all of the plasmid types and comparing the number of plasmids of that
-- type in the initial and final cell type respectively to check that the
-- transition is possible and count the number of plasmids of the initial type.
(⇐) :: CellType -> CellType -> Int
(⇐) = fmap (extract . foldl' resolvFoldStep NothingSoFar) . zipWith joinCellTypes
  where extract (Done res) = res
        extract _ = 0
        joinCellTypes (pt, n2) (_, n1) = (pt, n1, n2)

-- | This type is used for the calculation of the possibility of cointegration
-- or resolution between two plasmid types, but works differently in each case.
-- In both cases, we iterate through the possible plasmid types comparing the
-- number of copies of that plasmid type in the original and final cell type.
-- For cointegration, the original plasmid types are the separate plasmids, the
-- final type is the cointegrate, and we calculate the number of ways this
-- cointegration can occur. For resolution, the original plasmid type is the
-- cointegrate, the final types are the resolvants, and we calculate the number
-- of plasmids of the original type. This type represents our state while we
-- loop through the plasmid types, and its values have slightly different
-- interpretations in the cointegration and resolution
-- cases.
--
-- * 'NothingSoFar', in both cases means that the numbers of each plasmid type
--   have been identical in the two cell types up to this point.
--
-- * For cointegration, 'HaveOriginal' means one of the original plasmid types
--   has been found, and is given as the first argument. For resolution, it
--   means the (only) original plasmid type has been found. In both cases, the
--   'Int' argument is the number of plasmids of the type that has been found.
--
-- * For cointegration, 'HaveFinal' means the plasmid type of the resulting
--   cointegrate has been found, and is given by the argument. For resolution,
--   it means one of the resolvant plasmid types has been found, and is given by
--   the argument.
--
-- * For cointegration, 'NeedFinal' means both original plasmid types have been
--   found, and we are now looking for the cointegrate type, which must be the
--   type given by the first argument. The second 'Int' is the calculated number
--   of ways for cointegration to occur. For resolution, it means the
--   cointegrate type and one resolvant type have been found, and we are looking
--   for the second resolvant, which must be of the type given by the argument.
--   The 'Int' is the copy number of the original type.
--
-- * For cointegration, 'NeedOriginal' means we have found the cointegrate type
--   and one of the original types, and are now looking for the other original
--   type, which is given in the first argument. The 'Int' is the number of
--   plasmids of the original type which has been found. For resolution, it
--   means that both resolvants have been found, and we are now looking for the
--   original cointegrate type, given by the first argument. The second argument
--   is meaningless.
--
-- * 'Done' means that all the plasmid types have been found. The argument is
--   the result.
--
-- * 'Impossible' means that it has been determined that the desired transition
--   between cell types is impossible.
data CointFoldState = NothingSoFar
                    | HaveOriginal PlasmidType Int
                    | HaveFinal PlasmidType
                    | NeedFinal PlasmidType Int
                    | NeedOriginal PlasmidType Int
                    | Done Int
                    | Impossible

-- | A single step of the iteration through all plasmid types. The second
-- argument gives the plasmid type being currently considered, and its abundance
-- in the initial and final cell types.
cointFoldStep :: CointFoldState -> (PlasmidType, Int, Int) -> CointFoldState
cointFoldStep NothingSoFar (pt, before, after) = case after - before of
    1 -> HaveFinal pt
    0 -> NothingSoFar
    -1 -> HaveOriginal pt before
    -2 -> let (l,a) = pt in NeedFinal (2*l,2*a) ((before * (before - 1)) `div` 2)
    _ -> Impossible
cointFoldStep (HaveOriginal (l1,a1) n1) (pt, before, after) = case after - before of
    1 -> let (lf,af) = pt in NeedOriginal (lf - l1, af - a1) n1
    0 -> HaveOriginal (l1,a1) n1
    -1 -> let (l2,a2) = pt in NeedFinal (l1 + l2, a1 + a2) (n1 * before)
    _ -> Impossible
cointFoldStep (HaveFinal (lf,af)) (pt, before, after) = case after - before of
    0 -> HaveFinal (lf,af)
    -1 -> let (l1,a1) = pt in NeedOriginal (lf - l1, af - a1) before
    -2 -> let (li,ai) = pt in if (lf == 2*li) && (af == 2*ai)
        then Done ((before * (before - 1)) `div` 2)
        else Impossible
    _ -> Impossible
cointFoldStep (NeedFinal fin res) (pt, before, after) = case after - before of
    1 -> if fin == pt then Done res else Impossible
    0 -> NeedFinal fin res
    _ -> Impossible
cointFoldStep (NeedOriginal pt1 n2) (pt, before, after) = case after - before of
    0 -> NeedOriginal pt1 n2
    -1 -> if pt1 == pt then Done (before * n2) else Impossible
    _ -> Impossible
cointFoldStep (Done res) (pt, before, after) = case after - before of
    0 -> Done res
    _ -> Impossible
cointFoldStep Impossible _ = Impossible

-- | A single step of the iteration through all plasmid types. The second
-- argument gives the plasmid type being currently considered, and its abundance
-- in the initial and final cell types.
resolvFoldStep :: CointFoldState -> (PlasmidType, Int, Int) -> CointFoldState
resolvFoldStep NothingSoFar (pt, before, after) = case after - before of
    2 -> let (l,a) = pt in NeedOriginal (2*l, 2*a) 0
    1 -> HaveFinal pt
    0 -> NothingSoFar
    -1 -> HaveOriginal pt before
    _ -> Impossible
resolvFoldStep (HaveOriginal (l0,a0) n0) (pt, before, after) = case after - before of
    2 -> let (lf,af) = pt in if (l0 == 2*lf) && (a0 == 2*af)
        then Done n0
        else Impossible
    1 -> let (lf,af) = pt in NeedOriginal (l0 - lf, a0 - af) n0
    0 -> HaveOriginal (l0,a0) n0
    _ -> Impossible
resolvFoldStep (HaveFinal (l1,a1)) (pt, before, after) = case after - before of
    1 -> let (l2, a2) = pt in NeedOriginal (l1 + l2, a1 + a2) 0
    0 -> HaveFinal (l1,a1)
    -1 -> let (l0,a0) = pt in NeedFinal (l0 - l1, a0 - a1) before
    _ -> Impossible
resolvFoldStep (NeedFinal fin n0) (pt, before, after) = case after - before of
    1 -> if fin == pt then Done n0 else Impossible
    0 -> NeedFinal fin n0
    _ -> Impossible
resolvFoldStep (NeedOriginal pt0 _) (pt, before, after) = case after - before of
    0 -> NeedOriginal pt0 0
    -1 -> if pt0 == pt then Done before else Impossible
    _ -> Impossible
resolvFoldStep (Done res) (pt, before, after) = case after - before of
    0 -> Done res
    _ -> Impossible
resolvFoldStep Impossible _ = Impossible

-- Other Biology

-- | Determine whether the given cell type is a homozygote type (and therefore
-- doomed to extinction).
doomed :: CellType -> Bool
doomed = (\(as, bs) -> (as == 0) || (bs == 0)) .
    foldl' (\(as, bs) ((l, a), n) -> (as + n * a, bs + n * (l - a))) (0,0)

-- | The total number of plasmids (physical molecules) in the given cell type.
plasmidCount :: CellType -> Int
plasmidCount = sum . fmap snd

-- | The total number of multimers (plasmids with more than one copy of the
-- plasmid genome) there are in the given cell.
multimerCount :: CellType -> Int
multimerCount  = sum . fmap (\((l, _), n) -> if l > 1 then n else 0)

-- | @startingType n celltype@ is 'True' if @celltype@ is the starting cell type
-- for a model with copy number set to @n@; that is it is the type with @n@
-- plasmid monomers, exactly one of which is mutant.
startingType :: Int -> CellType -> Bool
startingType n = all (\(pt, count) -> case pt of
    (1,0) -> count == (n - 1)
    (1,1) -> count == 1
    _ -> count == 0)

-- Cell type generation

-- | A possible plasmid in the cell, represented as a @(l,a)@ pair, where @l@ is
-- the total number of loci on the plasmid (i.e. the number of monomers it
-- contains), and @a@ is the number of those that are mutant.
type PlasmidType = (Int, Int)

-- | A possible cell type in the model, represented as a map from 'PlasmidType's
-- to the number of plasmids of that type in the cell.
type CellType = [(PlasmidType, Int)]

-- | Generate the possible plasmid types for a plasmid with the given copy
-- number.
plasmidtypes :: Int -> [PlasmidType]
plasmidtypes n = [ (l,a) | l <- [1..n], a <- [0..l] ]

msets :: Integral a => [a] -> a -> [[a]]
msets [] n | n == 0 = [[]]
           | otherwise = []
msets (w:ws) n | n < 0 = []
               | otherwise = [0..(n `div` w)] >>= \v -> fmap (v:) (msets ws (n - w*v))

-- | Generate all the possible cell types in a model where the plasmid copy
-- number is the given number.
celltypes :: Int -> [CellType]
celltypes n = let pt = plasmidtypes n in fmap (zip pt) (msets (fmap fst pt) n)

printPlasmidType :: PlasmidType -> String
printPlasmidType (loci, aloci) = "(" ++ replicate aloci 'A' ++ replicate (loci - aloci) 'B' ++ ")"

printCelltype :: CellType -> String
printCelltype = concatMap (\(pt,count) -> concat (replicate count (printPlasmidType pt)))

-- Utility

-- | Binomial coefficients as an integer function.
choose :: Int -> Int -> Int
n `choose` k | k > n = 0
             | otherwise =  foldl' go 1 [1..mn]
  where (mn,mx) = if k < n - k then (k, n - k) else (n - k, k)
        go a i = a * (mx + i) `div` i
```
