## Supplementary material for "Evolutionary rescue of bacterial populations by heterozygosity on multicopy plasmids": Numerical and simulation code: common.c

```
/* The code common between the regular and random replication models. */
#include <math.h>
#include <stdio.h>
#include <stdlib.h>

#include "simulation.h"

const char *fitness_usage_message = "There are two fitness function specifications permitted:\n"
	"  flat(smin, smax)\tthe flat, or dominant, fitness function\n"
	"  peaked(smin, smax)\tthe peak fitness function\n";

struct fitness *
alloc_fitness(const char *specification, size_t n)
{
	int res;
	double smin, smax, s;
	struct fitness *ret;
	size_t i;

	ret = calloc(n + 1, sizeof(struct fitness));

	res = sscanf(specification, " flat ( %lf , %lf ) ", &smin, &smax);
	if (res == 2) {
		ret[0].lambda = 1.0 + smin;
		ret[0].rho = (1.0 + smin) / (2.0 + smin);
		for (i = 1; i < n; i++) {
			ret[i].lambda = 1.0 + smax;
			ret[i].rho = (1.0 + smax) / (2.0 + smax);
		}
		ret[n].lambda = 1.0 + smin;
		ret[n].rho = (1.0 + smin) / (2.0 + smin);
		return ret;
	}

	res = sscanf(specification, " peaked ( %lf , %lf ) ", &smin, &smax);
	if (res == 2) {
		for (i = 0; i <= n; i++) {
			s = smax - (smax - smin) * ((double) abs(2*i - n)) / ((double) n);
			ret[i].lambda = 1.0 + s;
			ret[i].rho = (1.0 + s) / (2.0 + s);
		}
		return ret;
	}

	free(ret);
	return NULL;
}

void
free_mode(double **m, size_t n)
{
	size_t i;

	for (i = 0; i <= n; i++)
		free(m[i]);

	free(m);
}
```
