## Supplementary material for "Evolutionary rescue of bacterial populations by heterozygosity on multicopy plasmids": Numerical and simulation code: EvolutionaryRescue.hs

```
-- | This module contains the common code that can be shared between all the
-- evolutionary rescue models we use. The idea is that each model defines a type
-- @m@ that represents a particular model, an instance @'RescueModel' m@, and a
-- function @buildModel :: /some parameters/ -> m@. The user creates a new model
-- instance with the desired parameters using this function, and then calls the
-- 'establishmentProb', 'rescueProb', and 'finalDist' functions from this model
-- on it to get the desired results.
module EvolutionaryRescue (
    RescueModel(..),
    establishmentProb,
    finalDist,
    e,
    expIntSeries
) where

import Control.Arrow ((***))
import Numeric.LinearAlgebra (Matrix, Vector, cmap, eig, fromList, ident, konst,
    maxIndex, norm_1, realPart, scalar, sumElements, toColumns, tr, (!), (<\>),
    (#>), (<.>))

-- | A type which is an instance of the 'RescueModel' class represents a
-- branching process model of evolutionary rescue. A model @a@ (i.e. a value of
-- a type which is an instance of 'RescueModel') represents a multitype
-- branching process with @'typeCount' a@ types and an offspring distribution
-- generating function given by @'fdf' a@. The branching process may start off
-- from single cell of known type, or from a probability distribution on single
-- cells, or from a probability distribution on pairs of cells (represented as a
-- quadratic form): the starting conditions are given by @'mutantExtinction' a@.
-- Finally, the rescue probability can be calculated from a known establishment
-- probability with @'rescueProb' a@.
class RescueModel a where
    -- | The number of distinct types in the multitype branching process model
    -- (this is the size of the vectors passed to the 'fdf' function).
    typeCount :: a -> Int
    -- | Evaluate the generating function of the offspring distribution of the
    -- process and its Jacobian at the given point.
    fdf :: a -> Vector Double -> (Vector Double, Matrix Double)
    -- | The matrix of expected number of offspring for the branching process,
    -- the matrix whose \(i,j\) element is the expected number of offspring of
    -- type \(j\) from an individual of type \(i\).
    expectedOffspring :: a -> Matrix Double
    -- | How to calculate the establishment probability of the model from the
    -- extinction probabilities of the individual types. The establishment
    -- probability is 1 minus a quantity which is either a linear combination of
    -- the extinction probabilities, and this function returns @Left v@, where
    -- @v@ is the vector of weights of the linear combination, or a quadratic
    -- form in the extinction probabilities, and this function returns a matrix
    -- representation @m@ of the form as @Right m@. Starting the branching
    -- process from a single cell of type @i@ is a special case of a linear
    -- combination of extinction probabilities: this function should return
    -- @Left ('e' i ('typeCount' m))@.
    mutantExtinction :: a -> Either (Vector Double) (Matrix Double)
    -- | Calculate the rescue probability for the model, given the establishment
    -- probability.
    rescueProb :: a -> Double -> Double
    -- | Calculate the mean time of the occurrence of the first successfully
    -- estabishing rescue mutation, given the establishment probability.
    meanEmergenceTime :: a -> Double -> Double

-- | Given @f@ which evaluates a function and its Jacobian, find a root thereof
-- with Newton's method. Assumes convergence when the difference in estimates of
-- the root has L1 norm less than @tol@.
newton :: Double -> (Vector Double -> (Vector Double, Matrix Double))
    -> Vector Double -> Vector Double
newton tol f = loop
  where loop x = let (obj, jac) = f x in let δx = -(jac <\> obj) in
            if norm_1 δx < tol
                then x + δx
                else loop (x + δx)

-- | Calculate the establishment probability for the given model, with precision
-- up to the given tolerance @tol@ (technically, this is the tolerance for the
-- total error on all the extinction probabilities, not the establishment
-- probability itself).
establishmentProb :: RescueModel a => Double -> a -> Double
establishmentProb tol rm = 1 - case mutantExtinction rm of
    Left v -> v <.> extinctionProbs
    Right m -> extinctionProbs <.> (m #> extinctionProbs)
  where extinctionProbs = newton tol fdf' (konst 0.5 (typeCount rm))
        -- We want to find a fixed point of fdf, which will be a zero of fdf'
        fdf' q = let (obj,jac) = fdf rm q in
            (obj - q, jac - ident (typeCount rm))

-- | Calculate the distribution of individuals among types in the limit of
-- infinite time. If the establishment probability is zero, so that this
-- distribution doesn't exist, then returns the zero vector.
finalDist :: RescueModel a => a -> Vector Double
finalDist = uncurry leading . (cmap realPart *** cmap realPart) .
    eig . tr . expectedOffspring
  where dist x = x / scalar (sumElements x)
        leading :: Vector Double -> Matrix Double -> Vector Double
        leading λs xs = let i = maxIndex λs in if (λs ! i) > 1
            then dist (toColumns xs !! i)
            else 0 * λs

-- | A standard basis vector for \(\mathbf{R}^n\). @e i n@ is the vector of
-- length @n@ with @i@th element equal to 1 and all other elements zero.
e :: Int -> Int -> Vector Double
e i n = fromList (replicate i 0 ++ [1] ++ replicate (n-i-1) 0)

-- | The exponential integral function
-- \[
--   f(x) = \int_0^x \frac{e^t - 1}{t} \,dt
--     = \sum_{k=1}^\infty \frac{x^k}{k\cdot k!}.
-- \]
expIntSeries :: Double -> Double
expIntSeries x = x * evalPoly (take 200 coeffs)
  where evalPoly [] = 0
        evalPoly (a:as) = a + x*evalPoly as

coeffs :: [Double]
coeffs = loop 1 1
  where loop k acc = (recip (fromInteger (k * k * acc))):(loop (k + 1) (k * acc))
```
