## Supplementary material for "Evolutionary rescue of bacterial populations by heterozygosity on multicopy plasmids": Numerical and simulation code: generate-figures.sh

```
./regular-probs -s 20241031 -e -r 100000 \
	'flat(-0.1,2.00)' \
	'flat(-0.1,1.00)' \
	'flat(-0.1,0.50)' \
	'flat(-0.1,0.25)' \
	'flat(-0.1,0.10)' \
	'peaked(-0.1,2.00)' \
	'peaked(-0.1,1.00)' \
	'peaked(-0.1,0.50)' \
	'peaked(-0.1,0.25)' \
	'peaked(-0.1,0.10)' > probs-reg.dat
./random-probs -s 20241031 -e -r 100000 \
	'flat(-0.1,2.00)' \
	'flat(-0.1,1.00)' \
	'flat(-0.1,0.50)' \
	'flat(-0.1,0.25)' \
	'flat(-0.1,0.10)' \
	'peaked(-0.1,2.00)' \
	'peaked(-0.1,1.00)' \
	'peaked(-0.1,0.50)' \
	'peaked(-0.1,0.25)' \
	'peaked(-0.1,0.10)' > probs-rand.dat

./regular-traj -s 20241031 -n  5 -o 0.2 'flat(-0.1,1.0)'    > tr-reg.dat
./regular-traj -s 20241031 -n 10 -o 0.2 'flat(-0.1,1.0)'   >> tr-reg.dat
./regular-traj -s 20241031 -n 15 -o 0.2 'flat(-0.1,1.0)'   >> tr-reg.dat
./regular-traj -s 20241031 -n 20 -o 0.2 'flat(-0.1,1.0)'   >> tr-reg.dat
./regular-traj -s 20241031 -n  5 -o 0.2 'peaked(-0.1,1.0)' >> tr-reg.dat
./regular-traj -s 20241031 -n 10 -o 0.2 'peaked(-0.1,1.0)' >> tr-reg.dat
./regular-traj -s 20241031 -n 15 -o 0.2 'peaked(-0.1,1.0)' >> tr-reg.dat
./regular-traj -s 20241031 -n 20 -o 0.2 'peaked(-0.1,1.0)' >> tr-reg.dat
./random-traj  -s 20241031 -n  5 -o 0.2 'flat(-0.1,1.0)'    > tr-rand.dat
./random-traj  -s 20241031 -n 10 -o 0.2 'flat(-0.1,1.0)'   >> tr-rand.dat
./random-traj  -s 20241031 -n 15 -o 0.2 'flat(-0.1,1.0)'   >> tr-rand.dat
./random-traj  -s 20241031 -n 20 -o 0.2 'flat(-0.1,1.0)'   >> tr-rand
./random-traj  -s 20241031 -n  5 -o 0.2 'peaked(-0.1,1.0)' >> tr-rand.dat
./random-traj  -s 20241031 -n 10 -o 0.2 'peaked(-0.1,1.0)' >> tr-rand.dat
./random-traj  -s 20241031 -n 15 -o 0.2 'peaked(-0.1,1.0)' >> tr-rand.dat
./random-traj  -s 20241031 -n 20 -o 0.2 'peaked(-0.1,1.0)' >> tr-rand.dat

./plots
./plots-traj tr-reg.dat tr-rand.dat
./plots-supp probs-reg.dat probs-rand.dat
```
