## Supplementary material for "Evolutionary rescue of bacterial populations by heterozygosity on multicopy plasmids": Numerical and simulation code: Makefile

```
CFLAGS=-std=c99 -Wall -pedantic -march=native -O3 -g -D _POSIX_C_SOURCE=200809L
LDLIBS=-lgsl -lopenblas -lm

random-probs: random.o common.o probabilities.o
	$(CC) $(LDFLAGS) -o $@ random.o common.o probabilities.o $(LDLIBS)
regular-probs: regular.o common.o probabilities.o
	$(CC) $(LDFLAGS) -o $@ regular.o common.o probabilities.o $(LDLIBS)
random-traj: random.o common.o trajectories.o
	$(CC) $(LDFLAGS) -o $@ random.o common.o trajectories.o $(LDLIBS)
regular-traj: regular.o common.o trajectories.o
	$(CC) $(LDFLAGS) -o $@ regular.o common.o trajectories.o $(LDLIBS)

common.o: common.c simulation.h
probabilities.o: probabilities.c simulation.h
random.o: random.c simulation.h
regular.o: regular.c simulation.h
trajectories.o: trajectories.c simulation.h
```
