## Supplementary material for "Evolutionary rescue of bacterial populations by heterozygosity on multicopy plasmids": Numerical and simulation code: plots-supp.hs

```
{-# LANGUAGE LambdaCase, OverloadedStrings, TupleSections #-}
module Main where

import Prelude hiding (readFile)

import Control.Applicative (some, (<|>))
import Data.Attoparsec.Text.Lazy (Parser, char, decimal, double, parseOnly,
    skipSpace)
import Data.Text.IO (readFile)
import Diagrams
import Diagrams.Backend.PGF.CmdLine
import System.Environment (getArgs)
import System.Exit (exitFailure)

import EvolutionaryRescue
import Plots
import Random
import Regular

main :: IO ()
main = do
    (regularSimFile, randomSimFile) <- getArgs >>= \case
        [f1, f2] -> (,) <$> readFile f1 <*> readFile f2
        _ -> putStrLn "Provide two arguments." >> exitFailure
    regularSimData <- case parseOnly (readSimData Regular) regularSimFile of
        Left err -> putStrLn ("Parse error: " ++ err) >> exitFailure
        Right res -> return res
    randomSimData <- case parseOnly (readSimData Random) randomSimFile of
        Left err -> putStrLn ("Parse error: " ++ err) >> exitFailure
        Right res -> return res
    renderPGF "heterozygosity-supp-fig-est.tex" (mkWidth 343)
        (pestFig (regularSimData ++ randomSimData))
    renderPGF "heterozygosity-supp-fig-resc.tex" (mkWidth 343)
        (prescFig (regularSimData ++ randomSimData))
    renderPGF "heterozygosity-supp-fig-est-delta.tex" (mkWidth 343)
        (pestDeltaFig (deltas (regularSimData ++ randomSimData)))
    renderPGF "heterozygosity-supp-fig-resc-delta.tex" (mkWidth 343)
        (prescDeltaFig (deltas (regularSimData ++ randomSimData)))

-- The data to plot

data Prob = Establishment Double Double Double
          | Rescue Int Double Double Double
    deriving (Eq, Ord, Read, Show)

readSimData :: ReplicationMode ->
    Parser [((ReplicationMode, FitnessFunc, Double, Double, Int, Double), Prob)]
readSimData mode = some ((,) <$>
    ((mode,,,,,) <$>
        (skipSpace *> (("flat" *> pure Dominant) <|> ("peaked" *> pure Peaked))) <*>
        (char '(' *> double) <*>
        (char ',' *> double) <*>
        (char ')' *> skipSpace *> int) <*>
        (skipSpace *> double)) <*>
    ((skipSpace *>
        (("rescue(" *> (Rescue <$>
            int <*>
            (char ')' *> skipSpace *> double) <*>
            (skipSpace *> double) <*>
            (skipSpace *> double))) <|>
        ("establishment" *> (Establishment <$>
            (skipSpace *> double) <*>
            (skipSpace *> double) <*>
            (skipSpace *> double)))))))
  where int :: Parser Int
        int = decimal

deltas ::
    [((ReplicationMode, FitnessFunc, Double, Double, Int, Double), Prob)] ->
    [((ReplicationMode, FitnessFunc, Double, Double, Int, Double), Prob)]
deltas = fmap (\((m,f,smin,smax,n,u), prob) -> case (m, prob) of
    (Regular, Establishment lo sim hi) ->
        let mod = regularModel (evalFitFunc f) smax smin 0 n in
        let pest = establishmentProb 1e-6 mod in
        ((m,f,smin,smax,n,u), Establishment (lo-pest) (sim-pest) (hi-pest))
    (Regular, Rescue pop lo sim hi) ->
        let mod = regularModel (evalFitFunc f) smax smin (u*fromIntegral pop) n in
        let pres = rescueProb mod (establishmentProb 1e-6 mod) in
        ((m,f,smin,smax,n,u), Rescue pop (lo - pres) (sim - pres) (hi - pres))
    (Random, Establishment lo sim hi) ->
        let mod = randomModel (evalFitFunc f) smax smin 0 n in
        let pest = establishmentProb 1e-6 mod in
        ((m,f,smin,smax,n,u), Establishment (lo-pest) (sim-pest) (hi-pest))
    (Random, Rescue pop lo sim hi) ->
        let mod = randomModel (evalFitFunc f) smax smin (u*fromIntegral pop) n in
        let pres = rescueProb mod (establishmentProb 1e-6 mod) in
        ((m,f,smin,smax,n,u), Rescue pop (lo - pres) (sim - pres) (hi - pres)))

-- Figures

pestFig ::
    [((ReplicationMode, FitnessFunc, Double, Double, Int, Double), Prob)] ->
    Diagram PGF
pestFig plotdata = finish (labelAxes "Plasmid copy number"
    "Establishment probability" (fourPanelLayout
        (scatterplot copyNumberScale probScale smaxScale)
        ["Dominant", "Peaked"] ["Regular", "Random"]
        [(dreg, mempty), (preg, k), (drand, mempty), (prand, mempty)]))
  where dreg = [(n, pest, smax) |
            ((Regular,Dominant,_,smax,n,_), Establishment _ pest _) <- plotdata]
        preg = [(n, pest, smax) |
            ((Regular,Peaked,_,smax,n,_), Establishment _ pest _) <- plotdata]
        drand = [(n, pest, smax) |
            ((Random,Dominant,_,smax,n,_), Establishment _ pest _) <- plotdata]
        prand = [(n, pest, smax) |
            ((Random,Peaked,_,smax,n,_), Establishment _ pest _) <- plotdata]
        k = key False (fmap (r2 . (94,)) [95,88..]) smaxScale

prescFig ::
    [((ReplicationMode, FitnessFunc, Double, Double, Int, Double), Prob)] ->
    Diagram PGF
prescFig plotdata = finish (labelAxes "Plasmid copy number" "Rescue probability"
    (fourPanelLayout (scatterplot copyNumberScale probScale smaxScale)
        ["Dominant", "Peaked"] ["Regular", "Random"]
        [(dreg, mempty), (preg, k), (drand, mempty), (prand, mempty)]))
  where dreg = [(n, pest, smax) |
            ((Regular, Dominant, _, smax, n, _), Rescue _ _ pest _) <- plotdata]
        preg = [(n, pest, smax) |
            ((Regular, Peaked, _, smax, n, _), Rescue _ _ pest _) <- plotdata]
        drand = [(n, pest, smax) |
            ((Random, Dominant, _, smax, n, _), Rescue _ _ pest _) <- plotdata]
        prand = [(n, pest, smax) |
            ((Random, Peaked, _, smax, n, _), Rescue _ _ pest _) <- plotdata]
        k = key False (fmap (r2 . (94,)) [95,81,74,60,53,46,39]) smaxScale

pestDeltaFig ::
    [((ReplicationMode, FitnessFunc, Double, Double, Int, Double), Prob)] ->
    Diagram PGF
pestDeltaFig plotdata = finish (labelAxes "Plasmid copy number"
    "Difference in establishment probability" (fourPanelLayout
        (scatterplotWithError copyNumberScale deltaScale smaxScale)
        ["Dominant", "Peaked"] ["Regular", "Random"]
        [(dreg, mempty), (preg, k), (drand, mempty), (prand, mempty)]))
  where dreg = [(n, low, pest, high, smax) |
            ((Regular,Dominant,_,smax,n,_), Establishment low pest high) <- plotdata]
        preg = [(n, low, pest, high, smax) |
            ((Regular,Peaked,_,smax,n,_), Establishment low pest high) <- plotdata]
        drand = [(n, low, pest, high, smax) |
            ((Random,Dominant,_,smax,n,_), Establishment low pest high) <- plotdata]
        prand = [(n, low, pest, high, smax) |
            ((Random,Peaked,_,smax,n,_), Establishment low pest high) <- plotdata]
        k = key False (fmap (r2 . (94,)) [95,88,81,18,11,4]) smaxScale

prescDeltaFig ::
    [((ReplicationMode, FitnessFunc, Double, Double, Int, Double), Prob)] ->
    Diagram PGF
prescDeltaFig plotdata = finish (labelAxes "Plasmid copy number"
    "Difference in rescue probability" (fourPanelLayout
        (scatterplotWithError copyNumberScale deltaScale smaxScale)
        ["Dominant", "Peaked"] ["Regular", "Random"]
        [(dreg, mempty), (preg, mempty), (drand, k), (prand, mempty)]))
  where dreg = [(n, low, pest, high, smax) |
            ((Regular,Dominant,_,smax,n,_), Rescue _ low pest high) <- plotdata]
        preg = [(n, low, pest, high, smax) |
            ((Regular,Peaked,_,smax,n,_), Rescue _ low pest high) <- plotdata]
        drand = [(n, low, pest, high, smax) |
            ((Random,Dominant,_,smax,n,_), Rescue _ low pest high) <- plotdata]
        prand = [(n, low, pest, high, smax) |
            ((Random,Peaked,_,smax,n,_), Rescue _ low pest high) <- plotdata]
        k = key False (fmap (r2 . (75,)) [95,88,81,74,11,4]) smaxScale
```
