## Supplementary material for "Evolutionary rescue of bacterial populations by heterozygosity on multicopy plasmids": Numerical and simulation code: plots-traj.hs

```
{-# LANGUAGE LambdaCase, OverloadedStrings, TupleSections #-}
module Main where

import Prelude hiding (readFile)

import Control.Applicative (some, (<|>))
import Data.Attoparsec.Text.Lazy (Parser, char, decimal, double, parseOnly,
    skipSpace)
import Data.Foldable (foldl')
import Data.List (groupBy)
import Data.Text.IO (readFile)
import Diagrams
import Diagrams.Backend.PGF.CmdLine
import System.Environment (getArgs)
import System.Exit (exitFailure)

import EvolutionaryRescue
import Plots
import Random
import Regular

main :: IO ()
main = do
    (regularSimFile, randomSimFile) <- getArgs >>= \case
        [f1, f2] -> (,) <$> readFile f1 <*> readFile f2
        _ -> putStrLn "Provide two arguments." >> exitFailure
    regularSimData <- case parseOnly (readSimData Regular) regularSimFile of
        Left err -> putStrLn ("Parse error: " ++ err) >> exitFailure
        Right res -> return res
    randomSimData <- case parseOnly (readSimData Random) randomSimFile of
        Left err -> putStrLn ("Parse error: " ++ err) >> exitFailure
        Right res -> return res
    renderPGF "heterozygosity-fig-traj.tex" (mkWidth 337)
        (trajFig (organizeData regularSimData ++ organizeData randomSimData))

-- The data to plot

readSimData :: ReplicationMode ->
    Parser [((ReplicationMode, FitnessFunc, Double, Double, Int, Double),
        (Int, Double, Double))]
readSimData mode = some ((,) <$>
    ((mode,,,,,) <$>
        (skipSpace *> (("flat" *> pure Dominant) <|> ("peaked" *> pure Peaked))) <*>
        (char '(' *> double) <*>
        (char ',' *> double) <*>
        (char ')' *> skipSpace *> int) <*>
        (skipSpace *> double)) <*>
    ((,,) <$>
        (skipSpace *> int) <*>
        (skipSpace *> (("inf" *> pure (read "Infinity")) <|> double)) <*>
        (skipSpace *> double)))
  where int :: Parser Int
        int = decimal

organizeData ::
    [((ReplicationMode, FitnessFunc, Double, Double, Int, Double), (Int, Double, Double))] ->
    [(ReplicationMode, FitnessFunc, Int, Source, [(Double, Double)])]
organizeData l = do
    (params, sims) <- simTrajectories l
    let (m, f, _, _, n, _) = params
    let meanOcc = numericalData params
    [(m, f, n, SimulationMean, meanAlong (filter rescued sims)),
        (m, f, n, Decoration, [(meanOcc,0), (meanOcc,100000)])] ++
        fmap (m,f,n,Simulation,) sims
  where fstEq (a,_) (b,_) = a == b
        fstEq' (a,_,_) (b,_,_) = a == b
        simTrajectories = fmap (\((paras,pt):rest) ->
            (paras,traj (pt:fmap snd rest))) . groupBy fstEq
        traj = fmap (filter ((< 120) . fst) . fmap (\(_,t,p) -> (t,p))) .
            groupBy fstEq'
        rescued = (>= 90000) . snd . last

meanAlong :: [[(Double, Double)]] -> [(Double, Double)]
meanAlong sims = (0, 100000):loop 1
  where interpolate ((t0,p0):(t1,p1):rest) t = if (t0 <= t) && (t <= t1)
            then (p0 + (p1 - p0) * (t - t0) / (t1 - t0), 1)
            else interpolate ((t1,p1):rest) t
        interpolate _ _ = (0, 0)
        mean = (\(x,l) -> x / fromIntegral l) .
            foldl' (\(x,l) (x',l') -> (x + x', l + l')) (0,0)
        meanAt t = mean (fmap ($ t) (fmap interpolate sims))
        loop t = let res = meanAt t in
            case (isNaN res, res >= 100000) of
                (True, _) -> []
                (_, False) -> (t, res):loop (t + 1)
                (_, _) -> [(t, res)]

numericalData :: (ReplicationMode, FitnessFunc, Double, Double, Int, Double) ->
    Double
numericalData (Regular, f, smin, smax, n, u) = 
    let mod = regularModel (evalFitFunc f) smax smin (100000*u) n in
    let pest = establishmentProb 1e-6 mod in
    meanEmergenceTime mod pest
numericalData (Random, f, smin, smax, n, u) = 
    let mod = randomModel (evalFitFunc f) smax smin (100000*u) n in
    let pest = establishmentProb 1e-6 mod in
    meanEmergenceTime mod pest

trajFig :: [(ReplicationMode, FitnessFunc, Int, Source, [(Double, Double)])] ->
    Diagram PGF
trajFig plotdata = finish (labelAxes "Time (mean cell lifespans)"
    "Population size" (fourPanelLayout
        (lineplot timeScale popScale copyNumberScale'')
        ["Dominant", "Peaked"] ["Regular", "Random"]
        [(dreg, k), (preg, mempty), (drand, mempty), (prand, mempty)]))
  where dreg = do
            (Regular, Dominant, n, src, dat) <- plotdata
            return ((n, src), dat)
        preg = do
            (Regular, Peaked, n, src, dat) <- plotdata
            return ((n, src), dat)
        drand = do
            (Random, Dominant, n, src, dat) <- plotdata
            return ((n, src), dat)
        prand = do
            (Random, Peaked, n, src, dat) <- plotdata
            return ((n, src), dat)
        k = key True (fmap (r2 . (90,)) [90,80..]) copyNumberScale''
```
