## Supplementary material for "Evolutionary rescue of bacterial populations by heterozygosity on multicopy plasmids": Numerical and simulation code: Plots.hs

```
{-# LANGUAGE LambdaCase #-}
module Plots (
    -- * General plotting functions, etc.
    finish,
    labelAxes,
    -- ** Plot layouts
    fourPanelLayout,
    colLayout,
    -- ** Plot (panel) types
    Plot,
    scatterplot,
    scatterplotWithError,
    impulseplot,
    lineplot,
    -- ** Scales
    ContScale,
    DiscScale,
    key,
    -- * Specific scales
    probScale,
    copyNumberScale,
    copyNumberScale',
    copyNumberScale'',
    smaxScale,
    smaxScale',
    propScale,
    fracScale,
    modelScale,
    deltaScale,
    timeScale,
    popScale,
    -- * Types to represent the input data
    ReplicationMode(..),
    FitnessFunc(..),
    evalFitFunc,
    -- ** Graph specific
    Source(..)
) where

import Data.Colour.SRGB (Colour, sRGB24)
import Data.Foldable (fold)
import Diagrams hiding (dashing, loc, scale)
import Diagrams.Backend.PGF (PGF)
import Diagrams.TwoD hiding (scale)
import Text.Printf (printf)

import Regular

data ReplicationMode = Regular
                     | Random
    deriving (Eq, Ord, Enum, Bounded, Read, Show)

data FitnessFunc = Dominant
                 | Peaked
    deriving (Eq, Ord, Enum, Bounded, Read, Show)

evalFitFunc :: FitnessFunc -> Double -> Double
evalFitFunc Dominant = flat
evalFitFunc Peaked = Regular.triangle

data Source = Simulation
            | SimulationMean
            | Decoration

finish :: Diagram PGF -> Diagram PGF
finish = frame 5 . fontSize (local 5)

labelAxes :: String -> String -> Diagram PGF -> Diagram PGF
labelAxes xlab ylab plot = hsep 12 [text ylab # rotateBy (1/4),
    vsep 12 [plot # center, text xlab]]

panelLabels :: [Diagram PGF]
panelLabels = fmap (translate (r2 (5, 95)) . text . ("\\sffamily\\bfseries " ++)
    . (:[])) ['A'..'Z']

fourPanelLayout :: Plot f a -> [String] -> [String] ->
    [(f a, Diagram PGF)] -> Diagram PGF
fourPanelLayout plot cols rows = vsep 6 .
    fmap (hsep 6 . fstmap (yscale plot <>)) .
    zipWith (\l ps -> lstmap (l <>) ps) rowLabels .
    lstmap (fmap (xscale plot <>)) .
    fstmap (zipWith (<>) colLabels) .
    splitIntoRows .
    zipWith (<>) panelLabels.
    fmap (\(dat, extra) -> plotFunc plot dat <> extra)
  where splitIntoRows [] = []
        splitIntoRows panels = let (row, rest) = splitAt (length cols) panels in
            row:splitIntoRows rest
        fstmap _ [] = []
        fstmap f (x:xs) = (f x):xs
        lstmap _ [] = []
        lstmap f (x:[]) = (f x):[]
        lstmap f (x:xs) = x:(lstmap f xs)
        colLabels = fmap (translate (r2 (50,106)) . text) cols
        rowLabels = fmap (translate (r2 (106,50)) . rotateBy (-1/4) . text) rows

colLayout :: Plot f a -> [(String, f a, Diagram PGF)] -> Diagram PGF
colLayout plot = vsep 6 .
    lstmap (xscale plot <>) .
    zipWith (<>) (fmap (translate (r2 (5,-50))) panelLabels) .
    fmap (\(lab, dat, extra) ->
        plotFunc plot dat <>
        extra <>
        yscale plot <>
        text lab # rotateBy (-1/4) # translate (r2 (211, 25)))
  where lstmap _ [] = []
        lstmap f (x:[]) = (f x):[]
        lstmap f (x:xs) = x:(lstmap f xs)

data Plot f a = Plot {
    xscale :: Diagram PGF,
    yscale :: Diagram PGF,
    plotFunc :: f a -> Diagram PGF
}

scatterplot :: Foldable f =>
    ContScale x -> ContScale y -> DiscScale p -> Plot f (x, y, p)
scatterplot xs ys ps = Plot {
    xscale = axisScale xs,
    yscale = axisScale ys,
    plotFunc = fold .
        (:[square 100 # alignBL # lw thin, adornment xs, adornment ys]) .
        foldMap (\(x, y, p) -> point ps p #
            fc (col ps p) #
            translate (r2 (scale xs x, scale ys y)))}

scatterplotWithError :: Foldable f =>
    ContScale x -> ContScale y -> DiscScale p -> Plot f (x, y, y, y, p)
scatterplotWithError xs ys ps = Plot {
    xscale = axisScale xs,
    yscale = axisScale ys,
    plotFunc = fold .
        (:[rect 102 100 # alignBL # lw thin, adornment xs, adornment ys]) .
        foldMap (\(x, lo, y, hi, p) -> point ps p #
                fc (col ps p) #
                translate (r2 (scale xs x + 0.5 * dodge ps p, scale ys y)) <>
            fromVertices [p2 (0, scale ys lo),p2 (0, scale ys hi)] #
                translateX (scale xs x + 0.5 * dodge ps p) #
                lc (col ps p) #
                lw (sqrt (2/3)))}

impulseplot :: Foldable f =>
    ContScale x -> ContScale y -> DiscScale g -> Plot f (x, y, g)
impulseplot xs ys gs = Plot {
    xscale = axisScale xs,
    yscale = translateX (-5) (axisScale ys),
    plotFunc = fold .
        (:[rect 210 50 # alignBL # translateX (-5) # lw thin,
            adornment xs, adornment ys]) .
        foldMap (\(x, y, g) -> rect 2 (scale ys y) #
            alignBL #
            translateX (scale xs x + 2 * dodge gs g) #
            fc (col gs g) #
            lw 0)}

lineplot :: Foldable f =>
    ContScale x -> ContScale y -> DiscScale l -> Plot f (l, [(x, y)])
lineplot xs ys ls = Plot {
    xscale = axisScale xs,
    yscale = axisScale ys,
    plotFunc = fold .
        (:[square 100 # alignBL # lw thin, adornment xs, adornment ys]) .
        foldMap (\(line, pts) ->
            fromVertices (fmap (\(x,y) -> p2 (scale xs x, scale ys y)) pts) #
            clipTo (square 100 # alignBL) #
            lc (col ls line) #
            opacity (op ls line) #
            dashingL (dashing ls line) (dodge ls line) #
            lw (local (linewidth ls line)))}

data ContScale a = CS {
    axisScale :: Diagram PGF,
    adornment :: Diagram PGF,
    scale :: a -> Double
}

data DiscScale a = DS {
    vals :: [a],
    label :: a -> String,
    point :: a -> Diagram PGF,
    dodge :: a -> Double,
    col :: a -> Colour Double,
    dashing :: a -> [Double],
    op :: a -> Double,
    linewidth :: a -> Double
}

key :: Bool -> [V2 Double] -> DiscScale a -> Diagram PGF
key linemode locs s = fold
    (zipWith (\loc v -> translate loc (if linemode
        then scaleline #
                lc (col s v) #
                dashingL (dashing s v) 0 #
                lw (local (linewidth s v)) <>
            alignedText 1 0.5 (label s v) # translateX (-6)
        else point s v # fc (col s v) <>
            alignedText 1 0.5 (label s v) # translateX (-5)))
    locs (vals s))
  where scaleline = fromVertices [p2 (-4.5,-4.5), p2 (4.5,4.5)] # lw 1.5

probScale :: ContScale Double
probScale = CS {
    axisScale = mconcat [
        alignedText 1 0.5 "$1.0$" # translate (r2 (-2, 100)),
        alignedText 1 0.5 "$0.8$" # translate (r2 (-2, 80)),
        alignedText 1 0.5 "$0.6$" # translate (r2 (-2, 60)),
        alignedText 1 0.5 "$0.4$" # translate (r2 (-2, 40)),
        alignedText 1 0.5 "$0.2$" # translate (r2 (-2, 20)),
        alignedText 1 0.5 "$0.0$" # translate (r2 (-2, 0))],
    adornment = mempty,
    scale = (100 *)}

copyNumberScale :: ContScale Int
copyNumberScale = CS {
    axisScale = mconcat [
        alignedText 0.5 1 "$20$" # translate (r2 (100, -2)),
        alignedText 0.5 1 "$15$" # translate (r2 (75, -2)),
        alignedText 0.5 1 "$10$" # translate (r2 (50, -2)),
        alignedText 0.5 1 "$5$" # translate (r2 (25, -2)),
        alignedText 0.5 1 "$0$" # translate (r2 (0, -2))],
    adornment = mempty,
    scale = (5 *) . fromIntegral}

copyNumberScale' :: DiscScale Int
copyNumberScale' = DS {
    vals = [2, 3, 4, 5],
    label = printf "$n = %d$",
    point = const (circle (sqrt (6 / pi)) # lw 0),
    dodge = const 0,
    col = \case
        -1 -> sRGB24 0x15 0x15 0x15
        2 -> sRGB24 0xa6 0x3d 0x40
        3 -> sRGB24 0xe9 0xb8 0x72
        4 -> sRGB24 0x90 0xa9 0x59
        5 -> sRGB24 0x64 0x94 0xaa
        _ -> sRGB24 0 0 0,
    dashing = \case
        -1 -> []
        2 -> [6, 2, 2, 2]
        3 -> [2, 2]
        4 -> [6, 2]
        5 -> []
        _ -> [1, 2],
     op = const 1,
     linewidth = const 1.5}

copyNumberScale'' :: DiscScale (Int, Source)
copyNumberScale'' = DS {
    vals = [(5,Simulation), (10,Simulation), (15,Simulation), (20,Simulation)],
    label = printf "$n = %d$" . fst,
    point = const (circle (sqrt (6 / pi)) # lw 0),
    dodge = fromIntegral . (`div` 5) . fst,
    col = \case
        (5, _) -> sRGB24 0xa6 0x3d 0x40
        (10, _) -> sRGB24 0xe9 0xb8 0x72
        (15, _) -> sRGB24 0x90 0xa9 0x59
        (20, _) -> sRGB24 0x64 0x94 0xaa
        _ -> sRGB24 0 0 0,
    dashing = \case
        (_, Simulation) -> []
        (_, SimulationMean) -> [6,1]
        (_, Decoration) -> [1,1],
    op = \case
        (_, Simulation) -> 0.25
        _ -> 1,
    linewidth = const 0.5}

propScale :: ContScale Double
propScale = CS {
    axisScale = mconcat [
        alignedText 0.5 1 "$1.0$" # translate (r2 (200, -2)),
        alignedText 0.5 1 "$0.8$" # translate (r2 (160, -2)),
        alignedText 0.5 1 "$0.6$" # translate (r2 (120, -2)),
        alignedText 0.5 1 "$0.4$" # translate (r2 (80, -2)),
        alignedText 0.5 1 "$0.2$" # translate (r2 (40, -2)),
        alignedText 0.5 1 "$0.0$" # translate (r2 (0, -2))],
    adornment = mempty,
    scale = (200 *)}

fracScale :: ContScale Double
fracScale = CS {
    axisScale = mconcat [
        alignedText 1 0.5 "$0.4$" # translate (r2 (-2, 50)),
        alignedText 1 0.5 "$0.3$" # translate (r2 (-2, 37.5)),
        alignedText 1 0.5 "$0.2$" # translate (r2 (-2, 25)),
        alignedText 1 0.5 "$0.1$" # translate (r2 (-2, 12.5)),
        alignedText 1 0.5 "$0.0$" # translate (r2 (-2, 0))],
    adornment = mempty,
    scale = (125 *)}

smaxScale :: DiscScale Double
smaxScale = DS {
    vals = [2, 1, 0.5, 0.25, 0.1],
    label = printf "$s_\\mathrm{max} = %.2f$",
    point = \case
        2.00 -> square (sqrt 6) # lw 0
        1.00 -> circle (sqrt (6 / pi)) # lw 0
        0.50 -> Diagrams.TwoD.triangle (sqrt (8 * sqrt 3)) # lw 0
        0.25 -> Diagrams.TwoD.triangle (sqrt (8 * sqrt 3)) # reflectY # lw 0
        0.10 -> square (sqrt 6) # rotateBy (1/8) # lw 0
        _ -> mempty,
    dodge = \case
        2.00 -> 2
        1.00 -> 1
        0.50 -> 0
        0.25 -> -1
        0.10 -> -2
        _ -> 0,
    col = \case
        2.00 -> sRGB24 0x15 0x15 0x15
        1.00 -> sRGB24 0xa6 0x3d 0x40
        0.50 -> sRGB24 0xe9 0xb8 0x72
        0.25 -> sRGB24 0x90 0xa9 0x59
        0.10 -> sRGB24 0x64 0x94 0xaa
        _ -> sRGB24 0 0 0,
    dashing = const [],
    op = const 1,
    linewidth = const 1.5}

smaxScale' :: ContScale Double
smaxScale' = CS {
    axisScale = mconcat [
        alignedText 0.5 1 "$3.0$" # translate (r2 (100, -2)),
        alignedText 0.5 1 "$2.5$" # translate (r2 (500/6, -2)),
        alignedText 0.5 1 "$2.0$" # translate (r2 (200/3, -2)),
        alignedText 0.5 1 "$1.5$" # translate (r2 (50, -2)),
        alignedText 0.5 1 "$1.0$" # translate (r2 (100/3, -2)),
        alignedText 0.5 1 "$0.5$" # translate (r2 (100/6, -2)),
        alignedText 0.5 1 "$0.0$" # translate (r2 (0, -2))],
    adornment = mempty,
    scale = (/ 3) . (100 *)}

modelScale :: DiscScale (ReplicationMode, FitnessFunc)
modelScale = DS {
    vals = [(Regular, Dominant), (Regular, Peaked),
        (Random, Dominant), (Random, Peaked)],
    label = \(m, f) -> show m ++ " " ++ show f,
    point = const (circle (sqrt (6 / pi)) # lw 0),
    dodge = \case
        (Regular, Dominant) -> -2
        (Regular, Peaked) -> -1
        (Random, Dominant) -> 0
        (Random, Peaked) -> 1,
    col = \case
        (Regular, Dominant) -> sRGB24 0xa6 0x3d 0x40
        (Regular, Peaked) -> sRGB24 0xe9 0xb8 0x72
        (Random, Dominant) -> sRGB24 0x90 0xa9 0x59
        (Random, Peaked) -> sRGB24 0x64 0x94 0xaa,
    dashing = const [],
    op = const 1,
    linewidth = const 2}

deltaScale :: ContScale Double
deltaScale = CS {
    axisScale = extrudeEnvelope (r2 (-6,0)) (mconcat [
        alignedText 1 0.5 "$ 0.02$" # translate (r2 (-2, 100)),
        alignedText 1 0.5 "$ 0.01$" # translate (r2 (-2, 75)),
        alignedText 1 0.5 "$ 0.00$" # translate (r2 (-2, 50)),
        alignedText 1 0.5 "$-0.01$" # translate (r2 (-2, 25)),
        alignedText 1 0.5 "$-0.02$" # translate (r2 (-2, 0))]),
    adornment = fromVertices [p2 (0,50), p2 (100,50)] #
        lc (sRGB24 0xC0 0xC0 0xC0) #
        lw (local 0.75),
    scale = (50 +) . (5000 *)}

timeScale :: ContScale Double
timeScale = CS {
    axisScale = mconcat [
        alignedText 0.5 1 "$120$" # translate (r2 (100, -2)),
        alignedText 0.5 1 "$ 90$" # translate (r2 (75, -2)),
        alignedText 0.5 1 "$ 60$" # translate (r2 (50, -2)),
        alignedText 0.5 1 "$ 30$" # translate (r2 (25, -2)),
        alignedText 0.5 1 "$  0$" # translate (r2 (0, -2))],
    adornment = mempty,
    scale = (/ 1.2)}

popScale :: ContScale Double
popScale = CS {
    axisScale = extrudeEnvelope (r2 (-8,0)) (mconcat [
        alignedText 1 0.5 "$10^5$" # translate (r2 (-2, 100)),
        alignedText 1 0.5 "$75000$" # translate (r2 (-2, 75)),
        alignedText 1 0.5 "$50000$" # translate (r2 (-2, 50)),
        alignedText 1 0.5 "$25000$" # translate (r2 (-2, 25)),
        alignedText 1 0.5 "$0$" # translate (r2 (-2, 0))]),
    adornment = mempty,
    scale = (/ 1000)}
```
