## Supplementary material for "Evolutionary rescue of bacterial populations by heterozygosity on multicopy plasmids": Numerical and simulation code: plots.hs

```
{-# LANGUAGE TupleSections #-}
module Main where

import Diagrams
import Diagrams.Backend.PGF.CmdLine
import Numeric.LinearAlgebra (toList)

import Cointegrates
import EvolutionaryRescue
import Plots
import Random
import Regular

main :: IO ()
main = do
    renderPGF "heterozygosity-fig-est.tex" (mkWidth 337) pestFig
    renderPGF "heterozygosity-fig-resc.tex" (mkWidth 337) prescFig
    renderPGF "heterozygosity-fig-fd.tex" (mkWidth 337) fdFig
    renderPGF "heterozygosity-fig-coint.tex" (mkWidth 337) cointFig

-- The data to plot

plotdata :: [(ReplicationMode, FitnessFunc, Double, Int, Double, Double, [Double])]
plotdata = do
    rep <- [Regular, Random]
    fit <- [Dominant, Peaked]
    smax <- [2, 1, 0.5, 0.25, 0.1]
    n <- [2..20]
    case rep of
        Regular -> let m = regularModel (evalFitFunc fit) smax (-0.1) 0.1 n in
                   let pest = establishmentProb 1e-6 m in
                   let fd = finalDist m in
                   return (rep, fit, smax, n, pest, rescueProb m pest, toList fd)
        Random -> let m = randomModel (evalFitFunc fit) smax (-0.1) 0.1 n in
                  let pest = establishmentProb 1e-6 m in
                  let fd = finalDist m in
                  return (rep, fit, smax, n, pest, rescueProb m pest, toList fd)

cointplotdata :: [(Double, Double, Int, Double)]
cointplotdata = do
     n <- [2..5]
     let cn = copyNumber n
     κ <- [0, 1e-4, 1e-2]
     smax <- [0.05,0.1..1.95] ++ [1.99] ++ [2.05,2.1..3]
     let m = cointegrateModel smax (-0.1) 0.1 (\i -> if i > 1 then κ else 0) (const 0) cn
     return (κ, smax, n, establishmentProb 1e-6 m)

-- Figures

pestFig :: Diagram PGF
pestFig = finish (labelAxes "Plasmid copy number" "Establishment probability"
    (fourPanelLayout (scatterplot copyNumberScale probScale smaxScale)
        ["Dominant", "Peaked"] ["Regular", "Random"]
        [(dreg, mempty) ,(preg, k), (drand, mempty), (prand, mempty)]))
  where dreg = [(n, pest, smax) | (rep, fit, smax, n, pest, _, _) <- plotdata,
            rep == Regular, fit == Dominant]
        preg = [(n, pest, smax) | (rep, fit, smax, n, pest, _, _) <- plotdata,
            rep == Regular, fit == Peaked]
        drand = [(n, pest, smax) | (rep, fit, smax, n, pest, _, _) <- plotdata,
            rep == Random, fit == Dominant]
        prand = [(n, pest, smax) | (rep, fit, smax, n, pest, _, _) <- plotdata,
            rep == Random, fit == Peaked]
        k = key False (fmap (r2 . (94,)) [95,88..]) smaxScale

prescFig :: Diagram PGF
prescFig = finish (labelAxes "Plasmid copy number" "Rescue probability"
    (fourPanelLayout (scatterplot copyNumberScale probScale smaxScale)
        ["Dominant", "Peaked"] ["Regular", "Random"]
        [(dreg, mempty), (preg, k), (drand, mempty), (prand, mempty)]))
  where dreg = [(n, presc, smax) | (rep, fit, smax, n, _, presc, _) <- plotdata,
            rep == Regular, fit == Dominant]
        preg = [(n, presc, smax) | (rep, fit, smax, n, _, presc, _) <- plotdata,
            rep == Regular, fit == Peaked]
        drand = [(n, presc, smax) | (rep, fit, smax, n, _, presc, _) <- plotdata,
            rep == Random, fit == Dominant]
        prand = [(n, presc, smax) | (rep, fit, smax, n, _, presc, _) <- plotdata,
            rep == Random, fit == Peaked]
        k = key False (fmap (r2 . (94,)) [95,81,74,60,53,46,39]) smaxScale

fdFig :: Diagram PGF
fdFig = finish
    (labelAxes "Fraction of mutant plasmids in a cell" "Fraction of cells"
        (colLayout (impulseplot propScale fracScale modelScale) [
            ("$n = 4$", [(i / 4, p, (m, f)) | (m, f, smax, n, _, _, fd) <- plotdata,
                (i, p) <- zip [0..] fd, smax == 1.0, n == 4], k),
            ("$n = 5$", [(i / 5, p, (m, f)) | (m, f, smax, n, _, _, fd) <- plotdata,
                (i, p) <- zip [0..] fd, smax == 1.0, n == 5], mempty),
            ("$n = 7$", [(i / 7, p, (m, f)) | (m, f, smax, n, _, _, fd) <- plotdata,
                (i, p) <- zip [0..] fd, smax == 1.0, n == 7], mempty),
            ("$n = 11$", [(i / 11, p, (m, f)) | (m, f, smax, n, _, _, fd) <- plotdata,
                (i, p) <- zip [0..] fd, smax == 1.0, n == 11], mempty),
            ("$n = 13$", [(i / 13, p, (m, f)) | (m, f, smax, n, _, _, fd) <- plotdata,
                (i, p) <- zip [0..] fd, smax == 1.0, n == 13], mempty)]))
  where k = key True (fmap r2 [(120, 42), (185, 42), (120, 32), (185, 32)]) modelScale

cointFig :: Diagram PGF
cointFig = finish (labelAxes "$s_\\mathrm{max}$" "Establishment probability"
    (fourPanelLayout (lineplot smaxScale' probScale copyNumberScale') ["",""]
        ["",""] [
            (k0, text "$\\kappa = 0$" # translate (r2 (80,95))),
            (k4, k <> text "$\\kappa = 10^{-4}$" # translate (r2 (80,95))),
            (k2, text "$\\kappa = 10^{-2}$" # translate (r2 (80,95))),
            (kinf, text "$\\kappa = \\infty$" # translate (r2 (80,95)))]))
  where k0 = [
            (5, (0,0):[(smax, pest) | (κ, smax, n, pest) <- cointplotdata,
                κ == 0, n == 5]),
            (4, (0,0):[(smax, pest) | (κ, smax, n, pest) <- cointplotdata,
                κ == 0, n == 4]),
            (3, (0,0):[(smax, pest) | (κ, smax, n, pest) <- cointplotdata,
                κ == 0, n == 3]),
            (2, (0,0):[(smax, pest) | (κ, smax, n, pest) <- cointplotdata,
                κ == 0, n == 2])]
        k4 = [
            (5, (0,0):[(smax, pest) | (κ, smax, n, pest) <- cointplotdata,
                κ == 1e-4, n == 5]),
            (4, (0,0):[(smax, pest) | (κ, smax, n, pest) <- cointplotdata,
                κ == 1e-4, n == 4]),
            (3, (0,0):[(smax, pest) | (κ, smax, n, pest) <- cointplotdata,
                κ == 1e-4, n == 3]),
            (2, (0,0):[(smax, pest) | (κ, smax, n, pest) <- cointplotdata,
                κ == 1e-4, n == 2])]
        k2 = [
            (5, (0,0):[(smax, pest) | (κ, smax, n, pest) <- cointplotdata,
                κ == 1e-2, n == 5]),
            (4, (0,0):[(smax, pest) | (κ, smax, n, pest) <- cointplotdata,
                κ == 1e-2, n == 4]),
            (3, (0,0):[(smax, pest) | (κ, smax, n, pest) <- cointplotdata,
                κ == 1e-2, n == 3]),
            (2, (0,0):[(smax, pest) | (κ, smax, n, pest) <- cointplotdata,
                κ == 1e-2, n == 2])]
        kinf = [(-1, [(x, x/(1 + x)) | x <- [0,0.05..3]])]
        k = key True (fmap (r2 . (30,)) [55,65..]) copyNumberScale'
```
