## Supplementary material for "Evolutionary rescue of bacterial populations by heterozygosity on multicopy plasmids": Numerical and simulation code: probabilities.c

```
#include <gsl/gsl_randist.h>
#include <gsl/gsl_rng.h>
#include <math.h>
#include <stdbool.h>
#include <stdio.h>
#include <stdlib.h>
#include <string.h>
#include <unistd.h>
#include <time.h>

#include "simulation.h"

static void
usage(const char *executable)
{
	fprintf(stderr, "usage: %s [options] FITNESS FUNCTIONS...\n\n"
		"Options:\n"
		"  -u REAL\tmutation rate to evaluate\n"
		"  -n INT\tcopy number to evaluate\n"
		"  -e\t\treport extablishment probability\n"
		"  -r INT\treport rescue probability for the given initial population size\n"
		"  -i INT\tthe number of simulation replications for each set of parameters (default 50000)\n"
		"  -s INT\tset the random number generator seed\n"
		"\n"
		"  The -u, -n, and -r arguments may be specified multiple times.\n"
		"\n"
		"  In the absence of options, the default is to evaluate for:\n"
		"    u = 10⁻⁶,\n"
		"    n = 2 through 20,\n"
		"  reporting only establishment probability.\n"
		"\n"
		"Each fitness function specification should appear in a seperate argument.\n"
		"%s",
		executable, fitness_usage_message);
}

/*
 * Calculate a point estimate (p) and 95% confidence interval (lower, upper) for
 * the probability parameter of a binomial distribution with observation xi and
 * total number of cases ni using the likelihood-ratio test and Newton's method.
 */
static void
conf_interval(unsigned xi, unsigned ni, double *p, double *lower, double *upper)
{
	double x, n, nmx, fixed, ll, lu, tmp, d;

	x = (double) xi;
	nmx = (double) ni - xi;
	n = (double) ni;

	*p = x / n;

	if (xi == 0) {
		*lower = 0.0;
		*upper = -expm1(-1.920729410347061/n);
		return;
	} else if (xi == ni) {
		*lower = exp(-1.920729410347061/n);
		*upper = 1.0;
		return;
	}

	/* We perform the Newton's method in logit space (solving for the logit
	 * of the interval endpoints): ll and lu hold the current estimate of
	 * the logit of the endpoint, while tmp holds the logarithm of the
	 * current estimate of the endpoint. */

	fixed = 1.920729410347061 - x*log(x) - nmx*log(nmx) + n*log(n);

	ll = log(x) - log(nmx) - 1;
	tmp = ll - log1p(exp(ll));
	do {
		d = (fixed - nmx*ll + n*tmp) / (x - n*exp(tmp));
		ll -= d;
		tmp = ll - log1p(exp(ll));
	} while (fabs(d) >= 1e-6);
	*lower = exp(tmp);

	lu = log(x) - log(nmx) + 1;
	tmp = lu - log1p(exp(lu));
	do {
		d = (fixed - nmx*lu + n*tmp) / (x - n*exp(tmp));
		lu -= d;
		tmp = lu - log1p(exp(lu));
	} while (fabs(d) >= 1e-6);
	*upper = exp(tmp);
}

/* One simulation step. */
static void
next_generation(const gsl_rng *rng, size_t n, const struct fitness *f, double **m, const unsigned *curr_gen, unsigned *next_gen)
{
	size_t i, j, k;
	unsigned parents, offspring[PAIRS(n + 1)];

	for (j = 0; j <= n; j++) {
		next_gen[j] = 0;
	}

	for (i = 0; i <= n; i++) {
		parents = gsl_ran_binomial(rng, f[i].rho, curr_gen[i]);
		gsl_ran_multinomial(rng, PAIRS(n + 1), parents, m[i], offspring);
		for (j = 0; j <= n; j++) {
			for (k = 0; k <= j; k++) {
				next_gen[j] += offspring[IDX(j, k)];
				next_gen[k] += offspring[IDX(j, k)];
			}
		}
	}
}

/* Run a simulation instance, and return whether the population is rescued
 * before going extinct. Arguments are a random number generator, the rescue
 * mode, the number of homozygote wild-type cells to start with, the number of
 * single-mutant cells to start with, and the threshold for rescue. */
static bool
rescued(const gsl_rng *rng, size_t n, const struct fitness *f, double **m, unsigned n_wild_type, unsigned n_mutant, unsigned threshold)
{
	double init_dist[PAIRS(n + 1)];
	unsigned init_pop[PAIRS(n + 1)], pop[2][n + 1];
	int curr_pop = 0;
	unsigned total;
	size_t i, j, k;

	memcpy(init_dist, m[0], sizeof(init_dist));
	init_dist[IDX(0,0)] = 0.0;
	/* init_dist is no longer normalized to sum to 1, but that's fine as
	 * gsl_ran_multinomial works with nonnormalized weights. */

	gsl_ran_multinomial(rng, PAIRS(n + 1), n_mutant, init_dist, init_pop);
	memset(pop, 0, sizeof(pop));
	pop[curr_pop][0] = n_wild_type;
	for (j = 0; j <= n; j++) {
		for (k = 0; k <= j; k++) {
			pop[curr_pop][j] += init_pop[IDX(j,k)];
			pop[curr_pop][k] += init_pop[IDX(j,k)];
		}
	}

	do {
		next_generation(rng, n, f, m, pop[curr_pop], pop[!curr_pop]);
		curr_pop = !curr_pop;
		total = 0;
		for (i = 0; i <= n; i++)
			total += pop[curr_pop][i];
		if (total == 0)
			return false;
	} while (total - pop[curr_pop][0] - pop[curr_pop][n] < threshold);

	return true;
}

int
main(int argc, char **argv)
{
	int c, tmpi;
	size_t i, j, k, l, m;
	unsigned sum, threshold;
	double tmpd, p, min, max;
	bool seen_i = false;
	long seed = time(NULL);

	size_t n_mutation_rates = 0;
	double mutation_rates[argc];
	size_t n_copy_numbers = 0;
	size_t copy_numbers[(argc > 19) ? argc : 19];
	size_t n_populations = 0;
	unsigned population_sizes[argc];
	size_t reps = 50000;

	gsl_rng *rng;
	struct fitness *fit;
	double **mod;

	while ((c = getopt(argc, argv, "u:n:er:i:s:")) != -1) {
		switch (c) {
			case 'u':
				tmpd = atof(optarg);
				for (i = 0; i < n_mutation_rates; i++)
					if (mutation_rates[i] == tmpd)
						break;
				mutation_rates[i] = tmpd;
				n_mutation_rates++;
				break;
			case 'n':
				tmpi = atoi(optarg);
				for (i = 0; i < n_copy_numbers; i++)
					if (copy_numbers[i] == tmpi)
						break;
				copy_numbers[i] = tmpi;
				n_copy_numbers++;
				break;
			case 'e':
				for (i = 0; i < n_populations; i++)
					if (population_sizes[i] == 0)
						break;
				population_sizes[i] = 0;
				n_populations++;
				break;
			case 'r':
				tmpi = atoi(optarg);
				for (i = 0; i < n_populations; i++)
					if (population_sizes[i] == tmpi)
						break;
				population_sizes[n_populations] = tmpi;
				n_populations++;
				break;
			case 'i':
				if (!seen_i) {
					reps = atoi(optarg);
					seen_i = true;
				} else {
					fprintf(stderr, "The -i argument can only be specified once.\n");
					exit(1);
				}
				break;
			case 's':
				seed = atol(optarg);
				break;
			default:
				fprintf(stderr, "Unknown argument: %c.", optopt);
				usage(argv[0]);
				exit(1);
		}
	}

	if (optind >= argc) {
		usage(argv[0]);
		exit(1);
	}

	if (n_mutation_rates == 0) {
		n_mutation_rates = 1;
		mutation_rates[0] = 1e-6;
	}
	if (n_copy_numbers == 0) {
		n_copy_numbers = 19;
		for (i = 0; i < 19; i++)
			copy_numbers[i] = i + 2;
	}
	if (n_populations == 0) {
		n_populations = 1;
		population_sizes[0] = 0;
	}

	rng = gsl_rng_alloc(gsl_rng_mt19937);
	gsl_rng_set(rng, seed);

	for (i = 0; i < n_mutation_rates; i++) {
		for (j = 0; j < n_copy_numbers; j++) {
			mod = alloc_mode(copy_numbers[j], mutation_rates[i]);
			for (k = optind; k < argc; k++) {
				fit = alloc_fitness(argv[k], copy_numbers[j]);
				if (fit == NULL) {
					fprintf(stderr, "Invalid fitness function: %s\n", argv[i]);
					usage(argv[0]);
					exit(1);
				}
				for (l = 0; l < n_populations; l++) {
					sum = 0;
					threshold = population_sizes[l] == 0 ? 100000 : population_sizes[l];
					for (m = 0; m < reps; m++)
						sum += rescued(rng, copy_numbers[j], fit, mod, population_sizes[l], population_sizes[l] == 0, threshold);
					conf_interval(sum, reps, &p, &min, &max);
					printf("%s\t%zd\t%e\t", argv[k], copy_numbers[j], mutation_rates[i]);
					if (population_sizes[l] == 0) {
						printf("establishment\t");
						/* Check that probability an establishment is a false positive is less than 0.1% */
						if (sum != 0 && log1p(-exp(-1e-2 / sum)) / log1p(-min) >= threshold)
							printf("FALSE POSITIVES TOO HIGH\n");
						else
							printf("%.5f\t%.5f\t%.5f\n", min, p, max);
					} else {
						printf("rescue(%d)\t%.5f\t%.5f\t%.5f\n", population_sizes[l], min, p, max);
					}
				}
				free(fit);
			}
			free_mode(mod, copy_numbers[j]);
		}
	}

}
```
