## Supplementary material for "Evolutionary rescue of bacterial populations by heterozygosity on multicopy plasmids": Numerical and simulation code: Random.hs

```
{-# LANGUAGE RecordWildCards, DeriveGeneric, DeriveAnyClass #-}
-- | The implementation of the random replication model, in which plasmids are
-- selected to be replicated randomly, creating a Pólya urn process.
module Random (
    RandomModel,
    randomModel,
) where

import Numeric (expm1, log1p)
import Numeric.LinearAlgebra (Matrix, Vector, assoc, cmap, diag, fromList,
    fromRows, konst, linspace, scalar, (<.>), (#>))
import Numeric.SpecFunctions (logChoose)
import Statistics.Distribution (logProbability)
import Statistics.Distribution.Hypergeometric (hypergeometric)

import EvolutionaryRescue

-- | A parameterized instance of the random replication model.
data RandomModel = RandomModel {
    -- | The plasmid copy number.
    n :: Int,
    -- | The \(i\)th element of this list is the matrix whose \(j,k\) element is
    -- \(P(i \to \{j,k\})\), not including the factor of 2 in the off-diagonal
    -- elements.
    reprProbs :: [Matrix Double],
    -- | The reproduction probability for each cell type.
    ρ :: Vector Double,
    -- | The quadratic form relating the extinction probability of the entire
    -- model to the extinction probabilities of the individual types. The
    -- \(i,j\) element of this matrix is the probability that the mutation first
    -- appears in a pair of cells with \(i\) and \(j\) mutant plasmids
    -- respectively.
    startForm :: Matrix Double,
    -- | The quantity \(unN_0(1+s_\mathrm{min})/s_\mathrm{min}\), needed to
    -- calculate the rescue probability.
    rescCoeff :: Double,
    -- | The quantity \(1/s_\mathrm{min}\), needed to calculate the mean time
    -- to emergence.
    etCoeff :: Double
}

instance RescueModel RandomModel where
    typeCount = (1+) . n
    fdf (RandomModel{..}) q = (objective, jacobian)
      where objective = 1 - ρ
                + ρ * fromList (fmap (\m -> q <.> m #> q) reprProbs)
            jacobian = diag ρ <> fromRows (fmap (\m -> 2*m #> q) reprProbs)
    expectedOffspring (RandomModel{..}) =
        diag ρ <> fromRows (fmap (#> konst 2 (n + 1)) reprProbs)
    mutantExtinction = Right . startForm
    rescueProb rm pest = - expm1 (pest * rescCoeff rm)
    meanEmergenceTime rm pest = etCoeff rm * exp (pest * rescCoeff rm) /
        expm1 (pest * rescCoeff rm) * expIntSeries (-pest * rescCoeff rm)

-- | Create a new rescue model with the given parameters. @randomModel domf smax
-- smin uN n@ is model with
--
-- * random replication,
-- * a fitness function with maximum value @smax@, minimum value @smin@, and
--   interpolation between them according to @domf@, a function
--   \([0,1]\to[0,1]\) that maps the proportion of mutant plasmids to the
--   fitness normalized between @smin@ and @smax@,
-- * copy number @n@, and
-- * product \(uN_0\) of mutation rate and initial wild-type population size of
--   @uN@.
randomModel :: (Double -> Double) -> Double -> Double -> Double -> Int
    -> RandomModel
randomModel domf smax smin uN n = RandomModel{..}
  where s = scalar (smax-smin) * cmap domf (linspace (n+1) (0,1)) + scalar smin
        ρ = (1 + s) / (2 + s)
        lRepDenom = lc (2*n - 1) n
        ln = log (fromIntegral n)
        -- In the following, i always represents the initial number of mutant
        -- plasmids, k the number after replication but before host cell
        -- division, and j the number in a daughter cell (and therefore j - k
        -- the number in the other daughter cell).
        reprProbs = fmap (\i -> assoc (n+1,n+1) 0 $ do
            k <- [i..n+i]
            let lRepNum = lc (2*n - k - 1) (n - k + i) + lc (k - 1) (k - i)
            let d = hypergeometric k (2*n) n
            j <- [max 0 (k-n)..min n k]
            return ((j,k-j), exp (lRepNum-lRepDenom+logProbability d j))) [0..n]
        startForm = assoc (n+1, n+1) 0 $ do
            k <- [1..n]
            let logRepPart = lc (2*n - k - 1) (n - 1)
                    + log (fromIntegral (2*n + (n-1)*k))
                    - ln - lRepDenom
                    - log (fromIntegral k) - log1p (fromIntegral k)
            let d = hypergeometric k (2*n) n
            j <- [0..k `div` 2]
            let lp = logRepPart + logProbability d j
            [((j, k - j), exp lp), ((k - j, j), exp lp)]
        rescCoeff = uN * (1 + smin) * fromIntegral n / smin
        etCoeff = recip smin

-- | Corrected version of the 'logChoose' function, which correctly evaluates
-- \(\log\binom{x}{0} = 0\).
lc :: Int -> Int -> Double
lc _ 0 = 0
lc n k = logChoose n k
```
