## Supplementary material for "Evolutionary rescue of bacterial populations by heterozygosity on multicopy plasmids": Numerical and simulation code: README

These files constitute the software used to produce the results in Dewan and
Uecker (2024), "Evolutionary rescue of bacterial populations by heterozygosity
on multicopy plasmids."

				    CONTENTS

The files can be divided into three general groups, in addition to this file and
generate-figures.sh, which contain documentation.

Numerical solutions
A library written in Haskell that calculate the establishment and rescue
probabilities, final distribution of cell types, and mean occurrence time of the
first successful rescue mutation from exact numerical equations. This group
contains:

EvolutionaryRescue.hs - General framework and common code for all models
Regular.hs          }
Random.hs           } - Specific models
Cointegrates.hs     }

Stochastic simulations
Programs written in C that simulate the rescue process to estimate
establishment and rescue probabilities, and provide population size
trajectories. This group contains:

simulation.h    - common header file for all components
probabilities.c - a program to estimate establishment and rescue probabilities
trajectories.c  - a program to sample population size trajectories during rescue
common.c        - code common to all models
regular.c       - code specific to the regular replication model
random.c        - code specific to the random replication model
Makefile        - build script

Plotting programs
Programs written in Haskell that use the output from the above programs to
produce the figures from the paper. This group contains:

Plots.hs      - an unnecessarily complicated graphing library
plots.hs      - generates Figures 3, 4, 6, and 7
plots-traj.hs - generates Figure 5
plots-supp.hs - generates the figures from the supplementary material

			       COMPILING AND USE

The Haskell programs can be compiled with GHC in the accustomed manner: the
figures in the paper were generated with GHC version 8.8.4. The Haskell
libraries on which the numerical solutions depend (together with the versions
used for the paper) are:
	base-4.13.0.0
	hmatrix-0.20.0.0
	math-functions-0.3.4.0
	statistics-0.15.2.0
The plotting programs further depend on:
	attoparsec-0.13.2.4
	colour-2.3.5
	diagrams-lib-1.4.3
	diagrams-pgf-1.4.2.1
	hmatrix-gsl-0.19.0.1
	text-1.2.4.0
The library is documented in Haddock comments.

To build the simulation programs, run
	make regular-probs random-probs regular-traj random-traj
The simulations depend on the GNU Scientific Library (version 2.8 used for the
simulations in the paper) and libopenblas (version 0.3.27).

To see usage documentation for the simulation programs, run them without
arguments. The plotting programs are run as
	./plots
	./plots-traj <output file from regular-traj> <output file from random-traj>
	./plots-supp <output file from regular-probs> <output file from random-probs>

The commands used to generate the figures for the paper are contained in the
file generate-figures.sh
