## Supplementary material for "Evolutionary rescue of bacterial populations by heterozygosity on multicopy plasmids": Numerical and simulation code: regular.c

```
#include <gsl/gsl_randist.h>
#include <stdlib.h>
#include <string.h>

#include "simulation.h"

double **
alloc_mode(size_t n, double u)
{
	size_t i, m;
	unsigned j, k;
	double mut_dist[2*n+1];
	double **res;

	res = calloc(n + 1, sizeof(double *));

	for (i = 0; i <= n; i++) {
		/* First calculate the distribution of the number of mutant
		 * plasmids after plasmids have replicated and any mutations
		 * have occurred. */
		memset(mut_dist, 0, sizeof(mut_dist));
		for (m = 0; m <= n - i; m++)
			mut_dist[2*i + m] = gsl_ran_binomial_pdf(m, u, n - i);
		/* Then use mut_dist and the hypergeometric distribution for
		 * segregation to calculate the distribution of pairs of
		 * daughters. */
		res[i] = calloc(PAIRS(n + 1), sizeof(double));
		for (j = 0; j <= n; j++) {
			for (k = 0; k <= j; k++) {
				res[i][IDX(j, k)] = mut_dist[j + k] * gsl_ran_hypergeometric_pdf(j, j + k, 2*n - j - k, n);
				if (j != k)
					res[i][IDX(j,k)] *= 2;
			}
		}
	}

	return res;
}
```
