## Supplementary material for "Evolutionary rescue of bacterial populations by heterozygosity on multicopy plasmids": Numerical and simulation code: Regular.hs

```
{-# LANGUAGE RecordWildCards #-}
-- | The implementation of the regular replication model, in which every plasmid
-- in the cell is duplicated before host reproduction.
module Regular (
    RegularModel,
    regularModel,
    -- ** Fitness functions
    flat,
    triangle,
    quadratic,
    singleMutant
) where

import Numeric (expm1)
import Numeric.LinearAlgebra (Matrix, Vector, assoc, cmap, diag, fromList,
    fromRows, linspace, scalar, (!), (#>))
import Statistics.Distribution (probability)
import Statistics.Distribution.Hypergeometric (hypergeometric)

import EvolutionaryRescue

-- | A parameterized instance of the regular replication model.
data RegularModel = RegularModel {
    -- | The copy number of the plasmid.
    n :: Int,
    -- | Matrix whose \(i,j\) element is \(P(i \to \{j, 2i - j\})\), not
    -- including the factor of 2 in the off-diagonal elements, which will be
    -- accounted for later.
    reprProbs :: Matrix Double,
    -- | The reproduction probability for each cell type.
    ρ :: Vector Double,
    -- | The quantity \(unN_0(1+s_\mathrm{min})/s_\mathrm{min}\), needed to
    -- calculate the rescue probability.
    rescCoeff :: Double,
    -- | The quantity \(1/s_\mathrm{min}\), needed to calculate the mean time
    -- to emergence.
    etCoeff :: Double
}

instance RescueModel RegularModel where
    typeCount = (1+) . n
    fdf (RegularModel{..}) q = (objective, jacobian)
      where objective = 1 - ρ + (common #> q)
            jacobian = 2*common
            common = diag ρ <> (reprProbs * qcomp)
            -- qcomp[i,j] = q[2*i-j], or zero if that index is out of bounds
            qcomp = assoc (n+1,n+1) 0 (do
                i <- [0..n]
                j <- [max 0 (2*i-n)..min n (2*i)]
                return ((i,j), q!(2*i-j)))
    expectedOffspring (RegularModel{..}) = diag ρ <> (2 * reprProbs)
    mutantExtinction = Left . e 1 . (1+) . n
    rescueProb rm pest = - expm1 (pest * rescCoeff rm)
    meanEmergenceTime rm pest = etCoeff rm * exp (pest * rescCoeff rm) /
        expm1 (pest * rescCoeff rm) * expIntSeries (-pest * rescCoeff rm)
        

-- | Create a new rescue model with the given parameters. @regularModel domf
-- smax smin uN n@ is model with
--
-- * regular replication,
-- * a fitness function with maximum value @smax@, minimum value @smin@, and
--   interpolation between them according to @domf@, a function
--   \([0,1]\to[0,1]\) that maps the proportion of mutant plasmids to the
--   fitness normalized between @smin@ and @smax@,
-- * copy number @n@, and
-- * product \(uN_0\) of mutation rate and initial wild-type population size of
--   @uN@.
regularModel :: (Double -> Double) -> Double -> Double -> Double -> Int
    -> RegularModel
regularModel domf smax smin uN n = RegularModel{..}
  where s = scalar (smax - smin) * cmap domf (linspace (n+1) (0,1)) + scalar smin
        ρ = (1 + s) / (2 + s)
        reprProbs = fromRows (fmap row [0..n])
        row i = let d = hypergeometric (2*i) (2*n) n in
            fromList (fmap (probability d) [0..n])
        rescCoeff = uN * (1 + smin) * fromIntegral n / smin
        etCoeff = recip smin

flat, triangle, quadratic, singleMutant :: Double -> Double
-- | The dominant fitness function, which is at a minimum for homozygotes and a
-- maximum for heterozygotes.
flat x | x == 0 || x == 1 = 0
       | otherwise        = 1
-- | The peaked fitness function, which interpolates linearly between minimum
-- fitness at 0 and 1, and maximum fitness at 0.5.
triangle = (1 +) . negate . abs . subtract 1 . (2 *)
-- | A quadratic fitness function, with minimum fitness at 0 and 1 and maximum
-- fitness at 0.5.
quadratic x = 4 * x * (1 - x)
-- | A fitness function without heterozygote advantage: cells with only
-- wild-type plasmids have minimum fitness, all others have maximum fitness.
singleMutant x | x == 0 = 0
               | otherwise = 1
```
