## Supplementary material for "Evolutionary rescue of bacterial populations by heterozygosity on multicopy plasmids": Numerical and simulation code: simulation.h

```
#ifndef SIMULATION_H_
#define SIMULATION_H_

#include <stddef.h>

/* The number of unordered pairs of numbers of mutant plasmids for the given
 * copy number; that is, the number of pairs (j,k) with 0 <= j < n and
 * 0 <= k <= j. */
#define PAIRS(n) ((n) * ((n) + 1) / 2)
/* Convert an ordered pair of integers (with j >= k) into an array index. */
#define IDX(j, k) ((j) * ((j) + 1) / 2 + (k))

/* The fitness data for a given type. */
struct fitness {
	double lambda; /* Reproduction rate. */
	double rho; /* Probability of reproducing before dying. */
};

/* Dynamically allocate a fitness function with the given name; returns NULL in
 * event of error. */
extern struct fitness *alloc_fitness(const char *, size_t);
/* A usage message for the user, describing the possible fitness functions. */
extern const char *fitness_usage_message;

/* Dynamically allocate a replication mode with the given copy number and
 * mutation rate. */
extern double **alloc_mode(size_t n, double u);
/* Free any resources associated with the given replication mode. */
extern void free_mode(double **, size_t);
#endif
```
