## Supplementary material for "Evolutionary rescue of bacterial populations by heterozygosity on multicopy plasmids": Numerical and simulation code: trajectories.c

```
#include <assert.h>
#include <gsl/gsl_randist.h>
#include <gsl/gsl_rng.h>
#include <math.h>
#include <stdio.h>
#include <string.h>
#include <unistd.h>
#include <time.h>

#include "simulation.h"

static void
usage(const char *executable)
{
	fprintf(stderr, "usage: %s [options] FITNESS FUNCTION\n\n"
		"Options:\n"
		"  Simulation parameters:\n"
		"    -u REAL\tmutation rate to evaluate (default 10⁻⁶)\n"
		"    -n INT\tcopy number to evaluate (default 5)\n"
		"    -p INT\tinitial population size (default 10⁵)\n"
		"  Control of the simulation:\n"
		"    -s INT\tset the random number generator seed\n"
		"  Output:\n"
		"    -i INT\tthe number of replicate trajectories (default 20)\n"
		"    -o TIME\thow often to report results in units of mean-time-to-death (default 10.0)\n"
		"\n"
		"%s",
		executable, fitness_usage_message);
}

static double
event(const gsl_rng *rng, size_t n, const struct fitness *f, double **m, unsigned *pop)
{
	size_t i, j, k;
	double res, tmp, total_rate = 0.0;
	unsigned offspring[PAIRS(n + 1)];

	for (i = 0; i <= n; i++)
		total_rate += pop[i] * (f[i].lambda + 1.0);

	res = gsl_ran_exponential(rng, 1.0 / total_rate);
	tmp = total_rate * gsl_rng_uniform(rng);
	for (i = 0; i <= n; i++) {
		tmp -= pop[i] * f[i].lambda;
		if (tmp < 0.0) {
			pop[i]--;
			gsl_ran_multinomial(rng, PAIRS(n + 1), 1, m[i], offspring);
			for (j = 0; j <= n; j++) {
				for (k = 0; k <= j; k++) {
					pop[j] += offspring[IDX(j, k)];
					pop[k] += offspring[IDX(j, k)];
				}
			}
			return res;
		}
		tmp -= 1.0 * pop[i];
		if (tmp < 0.0) {
			pop[i]--;
			return res;
		}
	}
	/* Should only get here if the population has died out. */
	assert(total_rate == 0.0);
	return INFINITY;
}

int
main(int argc, char **argv)
{
	int c;
	size_t i, j;
	double now, next_output;
	unsigned total;

	double mutation_rate = 1e-6;
	size_t copy_number = 5;
	unsigned init_pop = 100000;
	size_t reps = 20;
	long seed = time(NULL);
	double output_rate = 10.0;

	gsl_rng *rng;
	struct fitness *f;
	double **mode;

	while ((c = getopt(argc, argv, "u:n:p:i:s:o:")) != -1) {
		switch (c) {
			case 'u':
				mutation_rate = atof(optarg);
				break;
			case 'n':
				copy_number = atoi(optarg);
				break;
			case 'p':
				init_pop = atoi(optarg);
				break;
			case 'i':
				reps = atoi(optarg);
				break;
			case 's':
				seed = atol(optarg);
				break;
			case 'o':
				output_rate = atof(optarg);
				break;
			default:
				fprintf(stderr, "Unknown argument: %c.", optopt);
				usage(argv[0]);
				exit(1);
		}
	}

	rng = gsl_rng_alloc(gsl_rng_mt19937);
	gsl_rng_set(rng, seed);

	if (optind + 1 != argc) {
		usage(argv[0]);
		exit(1);
	}

	f = alloc_fitness(argv[optind], copy_number);
	if (f == NULL) {
		fprintf(stderr, "Invalid fitness function: %s\n", argv[optind]);
		exit(1);
	}
	mode = alloc_mode(copy_number, mutation_rate);

	unsigned pop[copy_number + 1];

	for (i = 0; i < reps; i++) {
		memset(pop, 0, (copy_number + 1) * sizeof(unsigned));
		pop[0] = init_pop;
		now = 0.0;
		printf("%s\t%zd\t%e\t%zd\t%.8f\t%u\n", argv[optind], copy_number, mutation_rate, i, 0.0, init_pop);
		next_output = 0.0;
		do {
			next_output += output_rate;

			do {
				now += event(rng, copy_number, f, mode, pop);
			} while (now < next_output);

			total = 0;
			for (j = 0; j <= copy_number; j++)
				total += pop[j];
			printf("%s\t%zd\t%e\t%zd\t%.8f\t%u\n", argv[optind], copy_number, mutation_rate, i, now, total);
		} while (total > 0 && total < reps * init_pop);
	}
}
```
